# Heterobifunctional Protein Binders Enable Cell Type-Specific Killing Through In-cell Enrichment

**DOI:** 10.1101/2025.05.16.654562

**Authors:** Ahmed Bulldan, Min Zheng, Christian Meyners, Patrick Purder, Johannes Krieger, Johannes Dreizler, Thomas M. Geiger, Max Repity, Marte Høen Lein, Ingrid Quist-Løkken, Noel Tewes, Martin Schwalm, Sarah Schlesiger, Sebastien Moniot, Stefan Knapp, Ingo V. Hartung, Toril Holien, Alexander Loewer, Felix Hausch

## Abstract

Non-catalytic heterobifunctional molecules promise to expand the range of therapeutic options by establishing complexes between key target proteins and accessory presenter proteins equipped with additional properties. Here, we systematically investigate the rational design of such molecules, explore the biochemical basis of complex formation and determine how they achieve cellular efficacy using the endogenously expressed immunophilin FKBP12 and the transcriptional regulator BRD4 as paradigms. We present classes of bifunctional molecules that enable selective, FKBP12-dependent killing of specific cell types at subnanomolar concentrations and allow to differentiate between closely similar bromodomains. We propose that the strongly potentiated efficacy of these bifunctional compounds is based on cellular enrichment through binding to the highly abundant presenter protein FKBP12, a mechanism we term “CellTrap”. Our findings substantiate the concept that highly expressed, non-essential proteins can be repurposed as selective recruiters to expand therapeutic windows of existing small-molecule inhibitors, opening new avenues for designing targeted drugs with improved cell-type specificity.

## Introduction

Proximity-inducing drugs are an emerging therapeutic modality with the potential to unlock gain-of-function pharmacology, cell-type specificity and difficult-to-drug proteins. Precedented in nature by molecular glues like FK506 and Rapamycin^1^, the field recently gained substantially preclinical and clinical traction with the use of designed heterobifunctional molecules as chemical inducers of protein-protein interactions (Fig. 1). This is most prominently exemplified by proteolysis targeting chimeras (PROTACs), which co-engage E3 ubiquitin ligases and target proteins to catalytically induce the degradation of the latter^2^. Inspired by PROTACs, several additional concepts of event-driven heterobifunctionals have emerged using phosphatases (PhosTACs, PhoRC), kinases (PHICS), deubiquitinating enzymes (DUBTACs), ribozymes (RIBOTACs), autophagic vesicles (AUTACs), lysosomal compartments (LYTACs), or transcriptional/epigenetic chemical inducers of proximity (TCIPs)^3^ as hijacked effector proteins^4,5^.

**Figure 1.**
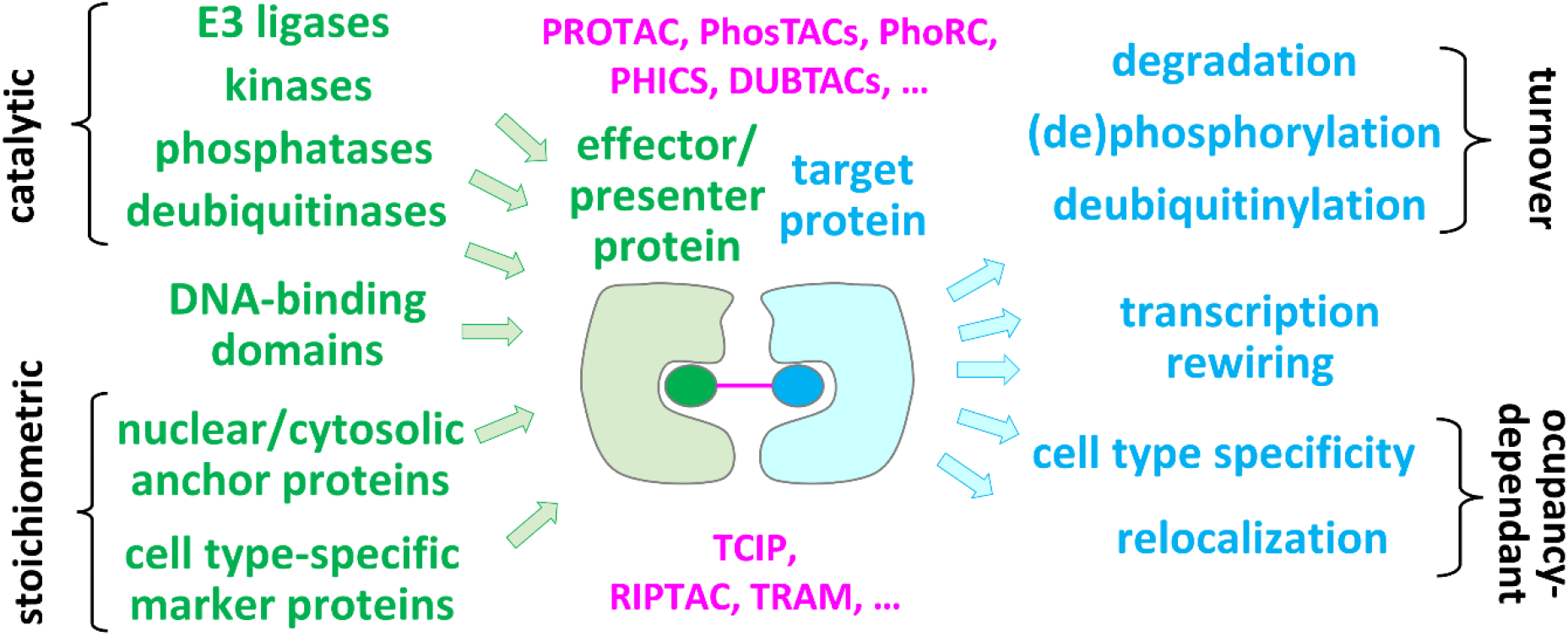
Examples of heterobifunctional drug modalities. Bifunctional molecules comprised of a ligand for the target protein (blue) and a ligand for an effector protein (green) are tethered by a linker (magenta) and allow recruitment of the latter to induce *de novo* functional effects. Catalytic effectors can operate at sub-stoichiometric drug levels whereas non-catalytic bifunctionals require persistent target occupancy.

More recently, heterobifunctional molecules have been proposed as therapeutics that do not engage enzymatic machineries of effector proteins but rely on sustained target protein occupancy (Fig. 1). These offer intriguing new modes of action. For example, exploiting presenter proteins (i.e., proteins that bind to one part of the heterobifunctional molecules) that are specifically expressed in diseased cells allows to increase therapeutic window of existing inhibitors of essential proteins. This can be realized by heterobifunctional RIPTACs (Regulated Induced Proximity-Targeting Chimeras)^6^, the first of which has recently entered clinical trials^7^ A related concept proposes to alter the subcellular localization of target proteins by anchoring them to cell compasrtment-specific proteins (TRAMs)^8^. Both approaches have been validated by proof-of-concept studies using exogenous overexpressed proteins but examples for effectively targeting endogenous proteins remain limited^9^. First clinical experience with non-degrading complex-inducing therapeutics has been gained with the immunosuppressants FK506, Rapamycin and Cyclosporin A and related tricomplex-inducing molecular glues^10–12^ that target difficult-to-drug proteins. However, the development of novel tricomplex inhibitors required a decade-long discovery effort with limited rational guidance. Bifunctional molecules that can build on pre-existing ligands for presenter and target proteins can be assembled and explored much faster.

Due to their larger size and higher complexity, bifunctional modalities are in general burdened by unfavorable pharmacological properties (e.g., reduced cell permeability, high cellular efflux, low solubility and resulting poor oral bioavailability). These challenges could be overcome for catalytically acting PROTACs (which require only low intracellular drug concentrations) by property-focused design which enabled reaching therapeutically active drug exposures in patients after oral dosing^13^. However, these pharmacokinetic hurdles remain relevant for bifunctionals that lack catalytical enhancement and therefore will require higher intracellular drug concentrations. Today, there is a lack of systematic studies on the behaviour and performance of this class of molecules and in turn their pharmacological prospect remains unclear.

To explore the potential of non-catalytic bifunctionals, we selected FKBP12 and BRD4 as paradigm presenter and target proteins. FKBP12 is the key presenter protein for the natural products FK506 and rapamycin^14^ and enable their molecular glue mechanism as immunosuppressants. Moreover, FKBP12 and engineered versions thereof are abundantly used as adaptor proteins for chemically induced protein-protein interactions in cells^1,15,16^. BRD4 is a member of the BET family of epigenetic reader proteins. It regulates the basal transcriptional machinery, mediates protein interactions in superenhancers and plays a crucial role in lineage specification^17^. Among the genes regulated by BRD4 are disease-associated genes such as the oncogene MYC and KEAP1. Inhibition of BRD4 has therefore been suggested as mono- or combination therapy against several tumor entities. However, most clinical studies have so far been hampered by dose-limiting side effects.

Using these model proteins, we investigated the various dimensions that determine the potency and efficacy of non-catalytic bifunctionals. Most significantly, we found that the cell killing potency of standard BRD4 inhibitors can be substantially potentiated in a cell-type specific manner by designed bifunctional molecules comprising highly tunable drug-like FKBP12 binders. We provide a first mathematical model rationalizing our findings to provide a basis for systematically unlocking the potential of this exciting new drug modality.

## Results

### Synthesis of dual BRD4- and FKBP12-binding conjugates

To generate BRD4-FKBP12 bifunctional molecules, we used the prototypic BET inhibitor (+)- JQ1^18^ and [4.3.1]bicyclic high-affinity ligands for FKBP12^19–22^. For both ligands the structure-activity relationship is well understood^15,23^, the binding mode is supported by co-crystal structures, and several exit vectors for conjugation have been validated^24^. We first generated a small library of BRD4/FKBP12-targeting heterobifunctional compounds by exploring 2 exit vectors for (+)-JQ1, 13 different amine/azide- or carboxy/azide-bifunctional linkers of varying length, and 9 different FKBP12 ligand variants, each containing an alkyne (Fig. 2A). Our linker selection comprised flexible, partially and fully rigidified designs including those which have been used in PROTACs with drug-like overall properties.

**Figure 2.**
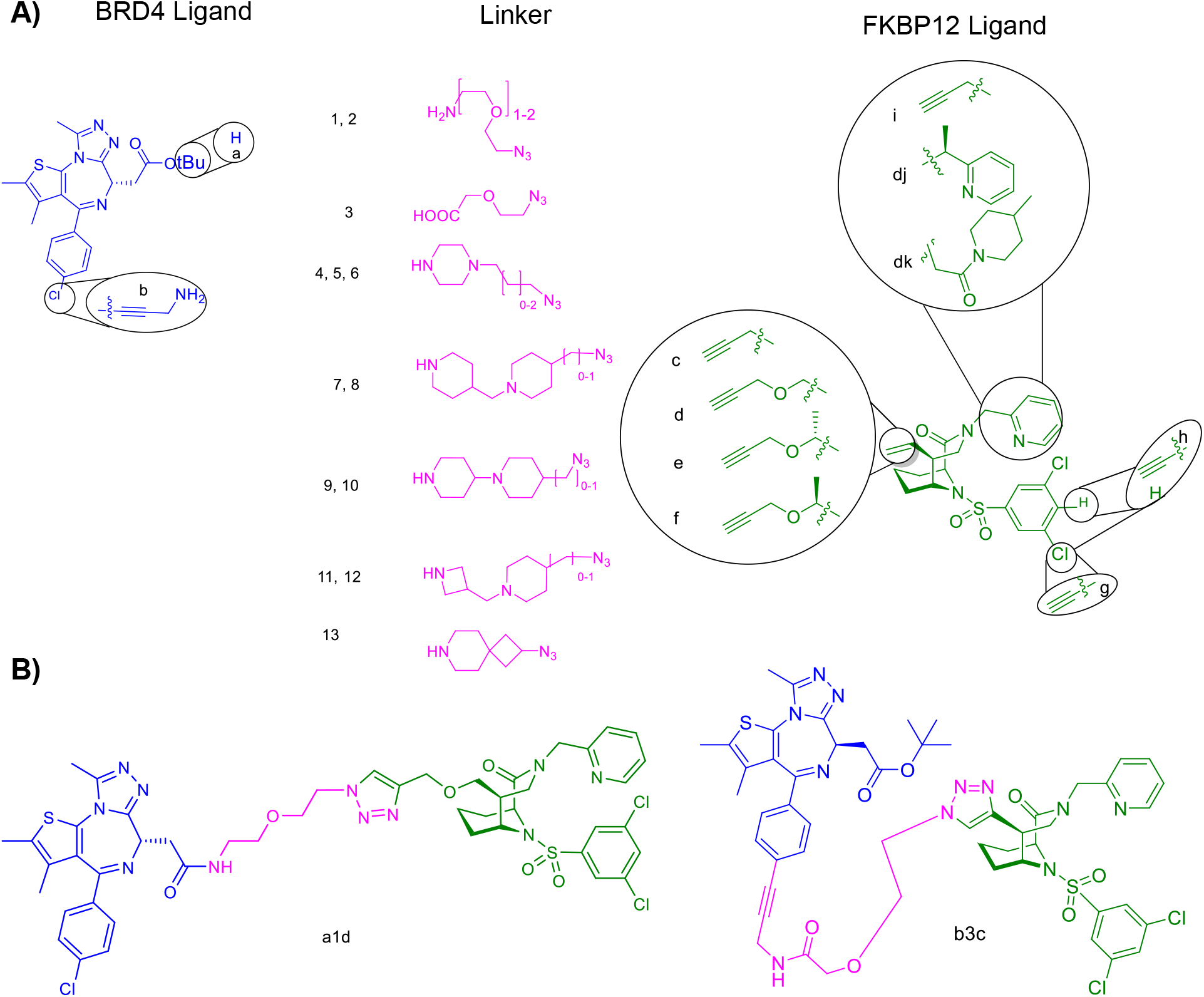
Chemical assembly of the bifunctional BRD4-FKBP12 ligand conjugates. **A**) Structures of BRD4 ligands, amine/azide or acid /azide-bifunctionalized linkers and alkyne-functionalized FKBP ligands. The attachment points and attachment functionalities are highlighted in the circles with an alphabetical identifier, the linkers are identified by numbers. Note that FKBP12 ligands dj and dk combine the exit vector d and a replacement of the pyridine moiety. **B**) Structures of exemplary BRD4-FKBP12 ligand conjugates a1d and b3c.

The (+)-JQ1 derivatives were first coupled to the bifunctional linkers. The resulting intermediates were then conjugated via their azide moieties to the alkyne-functionalized FKBP12 ligands by copper-catalyzed click reaction (Fig. 2B). Overall, 31 BRD4/FKBP12-targeting heterobifunctionals were obtained in this way.

### Biochemical characterization of BRD4-FKBP12 heterobifunctional ligands

With these bifunctionals in hand, we first characterized the binding to FKBP12^25^ and to the isolated BD1 and BD2 domains of BRD4 (**SI Fig.1**) by competitive fluorescence polarization (FP) assays. All compounds displayed high to very high affinity for FKBP12 (<5nM for exit vectors c-f, <10nM for exit vector i, <100nM for exit vector g and h, Tab 1 and **SI Tab 1**), in line with previous observations^24^. The binding to BRD4^BD1^ and BRD4^BD2^ was substantially weaker for most conjugates (mean Ki = 909 nM), with no differences between BRD4^BD1^ and BRD4^BD2^. To assess simultaneous binding of FKBP12 and BRD4, we repeated the BRD4 binding assays in the presence of 15 µM FKBP12. This further compromised BRD4 binding in several cases (**SI Tab. 1**). However, for compounds a1d, a1dj, a13c, and b3c we observed a >25-fold positive cooperativity (Fig. 3, SI Fig. 2, and Tab. 1), i.e., >25-fold tighter binding to BRD4 in the presence of high concentrations of FKBP12. When pre-bound to FKBP12, these compounds bound to BRD4 as good as (+)-JQ1 or even better (for b3c). Of note, compound a13c exhibited high selectivity for BRD4^BD1^*vs* BRD4^BD2^ in an FKBP12-assisted manner (Table 1, Fig. 3B).

**Table 1.**
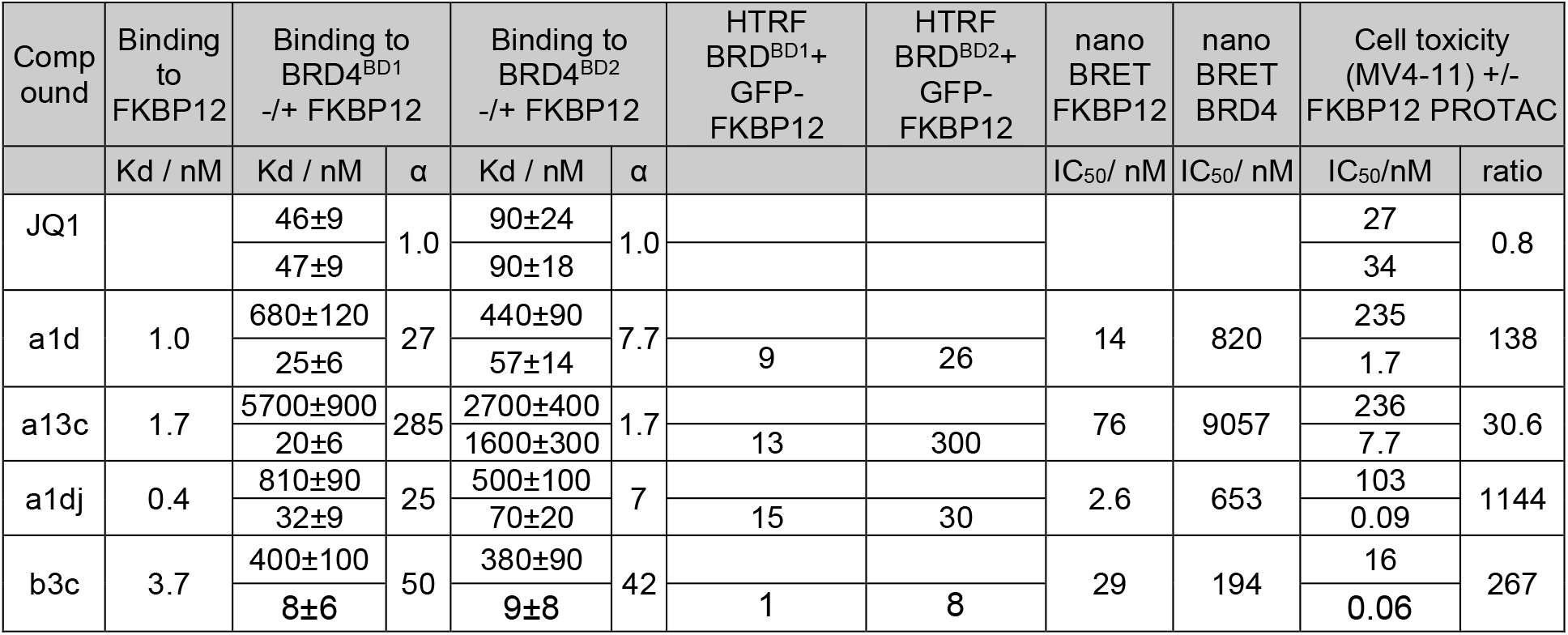
Characterization of BRD4-FKBP12 bifunctional ligands. Binding of FKBP12-BRD4 ligand conjugates to purified BRD4 BD1 or BD2 bromo domains (BRD4^BD1^ and BRD4^BD2^) was determined in a competitive fluorescence polarization assay in absence or presence of 15 µM FKBP12 using 2 nM of a fluorescently labeled JQ1-analog as a tracer. Binding to FKBP12 was determined in a competitive fluorescence polarization assay. Direct formation of the ternary complex was determined by a homogeneous time-resolved FRET (HTRF) assay using constant concentrations of eGFP-FKBP12 (5 µM) and His-BRD4^BD1 or BD2^ (100 nM) and serial dilution of bifunctionals. NanoBRET data were determined in intact HEK293 cells by expression of NanoLuc-fusion proteins and using fluorescently labeled FKBP12 or BRD4 ligands. Cell toxicity was determined in MV4-11 cells after five days of treatment.

**Fig. 3.**
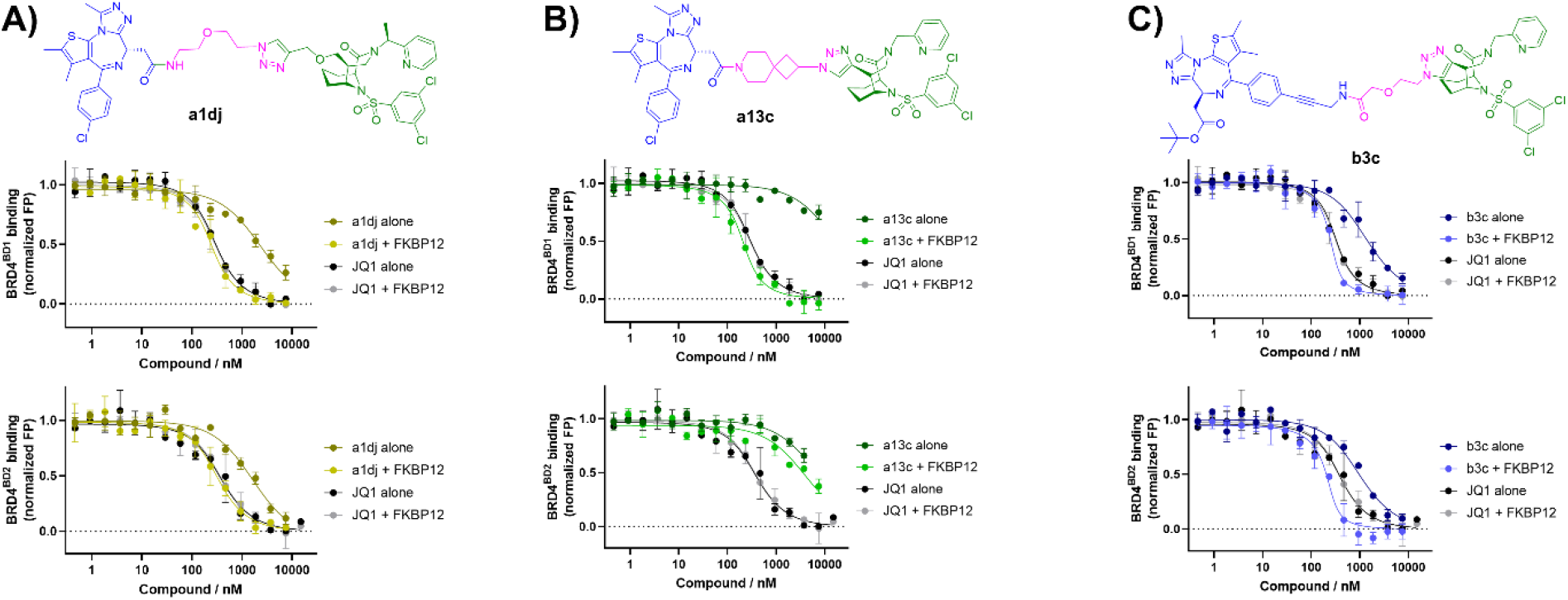
Positive cooperativity and selectivity of BRD4-FKBP12 bifunctional ligands. The binding to BRD4^BD1^ and BRD4^BD2^ of a1dj, a13c, b3c and JQ1 as reference was determined alone or in the presence of 15 µM FKBP12 by a competitive FP assay using a fluorescent JQ1 analog as tracer. Data points are indicated as mean from two replicates and standard deviations are indicated as error bars. The solid lines represent data fitted to a competitive binding model.

We also investigated ternary complex formation of the best compounds by a homogeneous time-resolved FRET assay (HTRF, **SI Fig. 3**). This confirmed the enhanced affinities of the FKBP12-compound complexes towards BRD4 as well as the BD1/BD2 selectivity of a13c. **(SI Tab. 2)**. Notably, we observed different upper plateaus for the FKBP12-BRD4^BD1^ complexes induced by b3c vs. those induced by a1d, a1dj, or a13c **(SI Fig. 3)**. This indicated the induction of ternary complexes with different architectures, i.e., with different relative orientation of the GFP tag on FKBP12 vs the His tag (complexed in the HTRF assay with a Terbium-labelled antibody) on BRD-BRD4^BD1^.

### Structure of the ternary complexes

To gain more insights into the underlying binding modes, we co-crystallized the ternary complexes of FKBP12:a1d:BRD4^BD1^ (Fig. 4) and FKBP12:b3c:BRD4^BD1^ **(SI Fig. 4A)**^26^. For a1d, two slightly different binding modes were observed, which were present as two different complexes in the crystallographic unit (Fig. 4A,B). The binding of the [4.3.1] bicyclic sulfonamide to FKBP12 and the JQ1-moiety to BRD4^BD1^ were almost identical to the respective binary complexes **(SI Fig. 5)**, as expected. The two ternary binding modes differ by an approx. 30° rotation of FKBP12 and a tilting relative to BRD4^BD1^ and are based on a partially distinct FKBP12-BRD4^BD1^ interaction patterns (Fig. 4D-G and **SI Fig. 6A,B**).

**Fig. 4.**
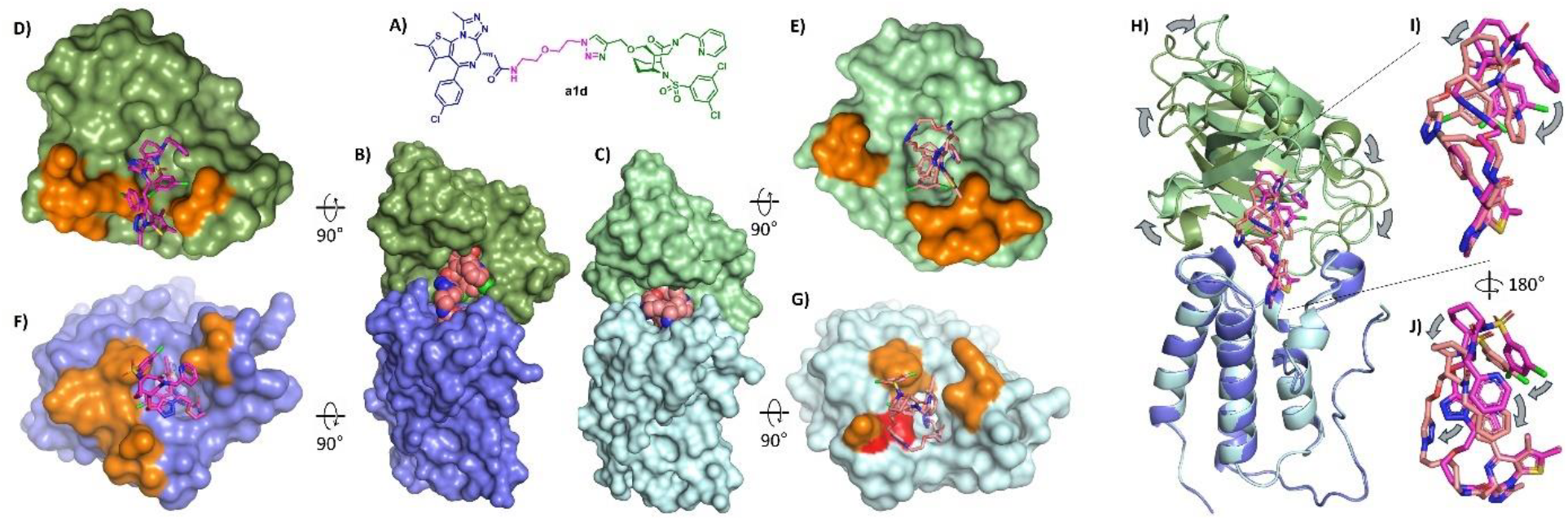
Structure of the FKBP12:a1d:BRD4^BD1^ ternary complex (PDB-ID 9QW8) B-J) FKBP12 is colored light/dark green, BRD4^BD1^ is colored cyan/dark blue, and compound a1d is colored pale pink/magenta. D-G) Amino residues forming ternary protein-protein contacts are highlighted in orange and amino acids contacting the linker are colored in red. **A)** Chemical structure of a1d. **B,C)** Two different ternary FKBP12:a1d:BRD4^BD1^ complexes are present in the crystallographic unit cell (sub-complexes a and b). **D,E)** a1d bound to FKBP12 in the two crystallographic units. BRD4^BD1^ is not shown for clarity. **F,G)** a1d bound to BRD4^BD1^ in the two crystallographic units. FKBP12 is not shown for clarity. **H-J)** Superposition of the two ternary complexes, aligned on the BRD4^BD1^ subunits. Repositing of FKBP12 and between the two subunits is indicated by arrows. **I,J)** Magnification of a1d in the two FKBP12-BRD4^BD1^-bound conformations, extracted from H) and viewed from two different angles.

**Fig. 5.**
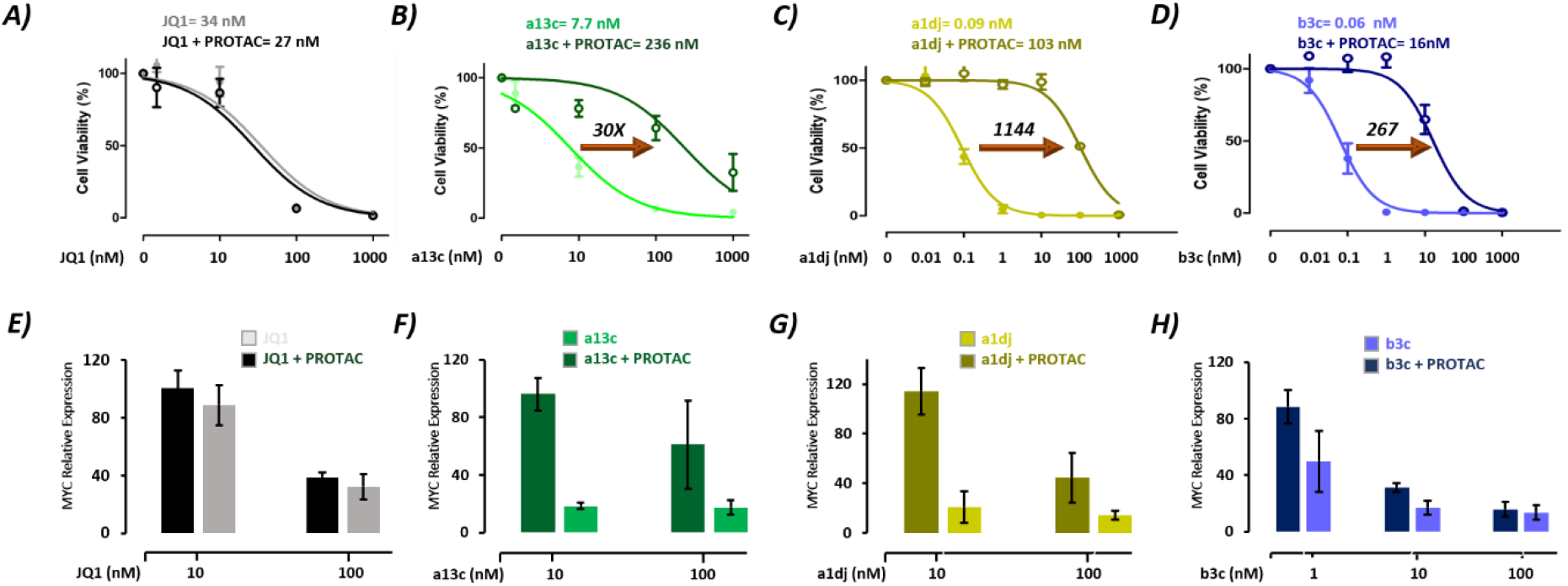
FKBP12-assisted toxicity by BRD4-targeting bifunctionals in MV4-11 cells. **A-D)** Cell toxicity for MV4-11 cells treated for five days with the indicated concentrations of (A) JQ1, (B) a13c, (C) a1dj, and (D) b3c. All experiments were performed in triplicate, either in the presence or absence of FKBP12 PROTAC. Means and standard errors are shown for each measurement. Lines indicate dose-dependent reduction of MYC expression. IC_50_ values in the presence or absence of FKBP12 PROTAC are indicated above each panel. The arrows indicate the fold change **E-H)** Quantitative analysis of MYC mRNA levels in MV4-11 cells following treatment with (E) JQ1, (F) a13c, (G) a1dj, and (H) b3c for six hours. These experiments were conducted in the presence or absence of PROTAC. Error bars represent the mean ± standard deviation (SD) from three independent experiments.

**Fig. 6.**
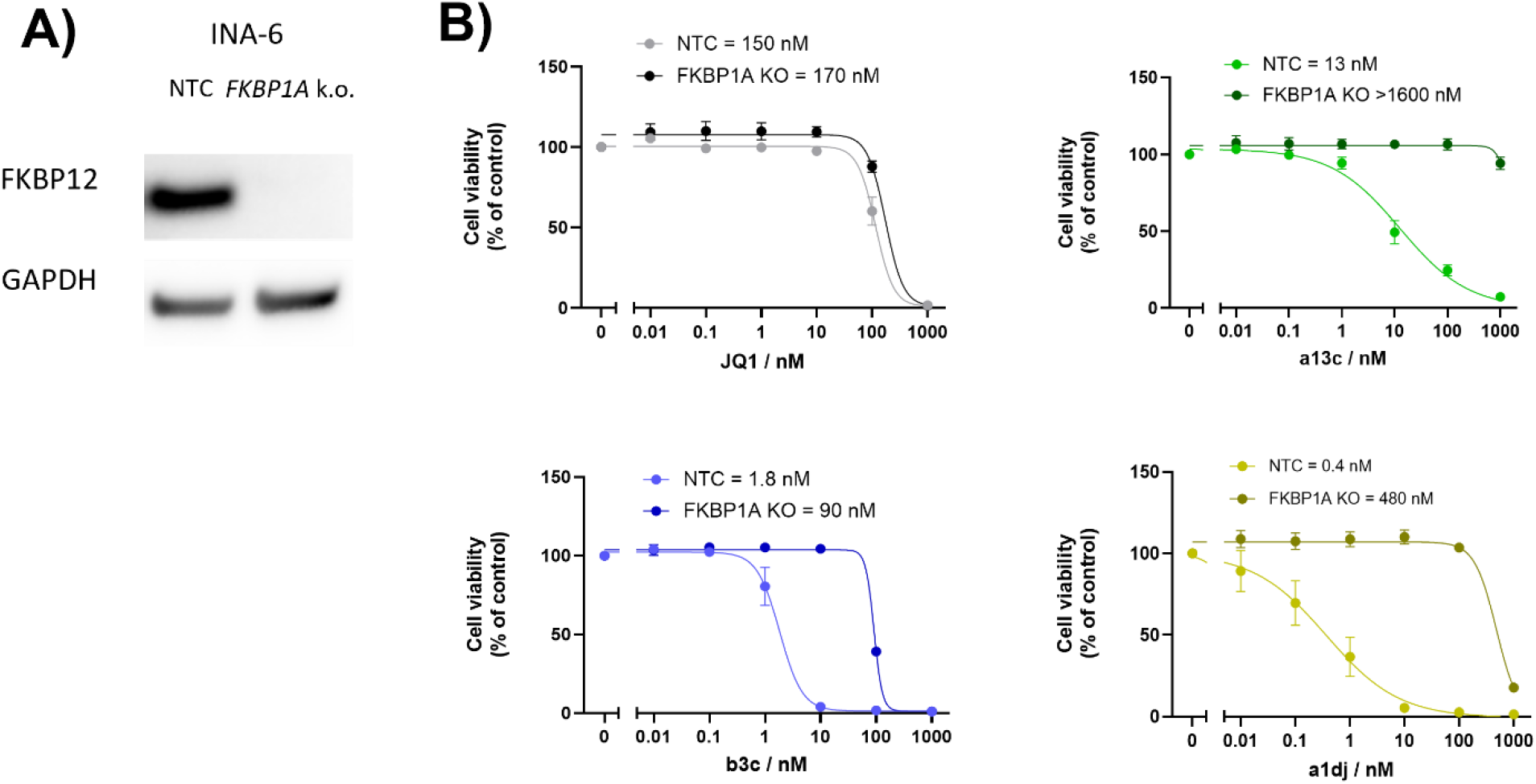
FKBP12-enhanced cell toxicity in myeloma cell lines. **A)** Western Blot analysis of INA-6 cells after FKBP12 knockout (FKBP1A gene) or of non-transfected control cells (NTC). **B)** Cell toxicity of JQ1 (grey), a13c (green), a1dj (yellow), and b3c (blue) in INA-6 cells after FKBP12 knockout or of non-targeted control (NTC) cells.

In addition to the protein-protein interactions, several ‘crossover’ ligand-protein contacts were observed, from the [4.3.1] bicyclic sulfonamide (i.e., from the FKBP12 ligand) to BRD4^BD1^ and from the JQ1-like structure to FKBP12 **(SI Fig. 6C-F)**. In one complex, the linker is involved in interactions with BRD4^BD1^ **(SI Fig. 6F)**. In both complexes substantial intramolecular contacts between the [4.3.1] bicyclic sulfonamide and JQ1-part were observed and both ligands interacted with the linker **(SI Fig. 6G,H)**. In both complexes, a methyl group in the α-position of the pyridyl group, which distinguishes a1d from a1dj, could be spatially accommodated. Notably, in the FKBP12:b3c:BRD4^BD1^ complex^26^ the relative FKBP12-BRD4^BD1^ orientation was drastically different compared to the FKBP12:a1d:BRD4^BD1^ complexes (SI Fig. 4B,C), with a substantially larger distances between the N-termini of FKBP12 and BRD4^BD1^. This is consistent with the reduced HTRF signals for the FKBP12:b3c:BRD4^BD1^ complex compared those for a1d or a1dj **(SI Fig. 3)**, since this assay relies on N-terminal FRET pairs.

Taken together, multiple solutions appear to exist to generate additional binding energy in the ternary complexes, which is crucial for positive cooperativity. This relies on direct protein-protein contacts, protein-non-cognate ligand ‘crossover’ contacts, and linker-protein contacts. Moreover, intramolecular contacts within the bifunctional molecule seemed to be important to stabilize the bioactive conformation.

The generated FKBP12-BRD4 ligand conjugates were rather large molecules (>1000 g/mol), which often have reduced cellular permeability. To assess the potential for cellular application, we investigated intracellular FKBP12 and BRD4 occupancy by nanoBRET assays ^27^. Gratifyingly, several FKBP12-BRD4 bifunctionals bound to FKBP12 or BRD4 in cells to a similar extent as the unconjugated precursor molecules (compound FK[4.3.1]-16h^28^ or compound 19-(S^Me^)^21^for FKBP12 and JQ1 or BRD4), incl. a1d, a1dj, a13c, and b3c (Tab. 1 and **SI Tab. 1**). In turn, we set out to explore the performance of these compounds in functional cellular assays.

### BRD4-FKBP12 bifunctional ligands display potent cytotoxicity depending on cellular FKBP12 levels

First, we evaluated the impact of selected BRD4-FKBP12 bifunctionals on cell viability in MV4-11 cells. This cell line is derived from a hematopoietic malignancy and was chosen based on their known sensitivity to BRD4 inhibition^29^ and high FKBP12 protein level **(SI Fig. 7A)**. In MV4-11 cells, we observed a dose-dependent decrease in cell viability for all compounds tested. Notably, BRD4-FKBP12 bifunctionals showed higher efficacies than JQ1 despite lower binding affinities than JQ alone (Tab. 1), with a1dj and b3c reaching sub-nanomolar IC50 values (Fig. 5A-D and **SI Fig. 8A**).

**Fig. 7.**
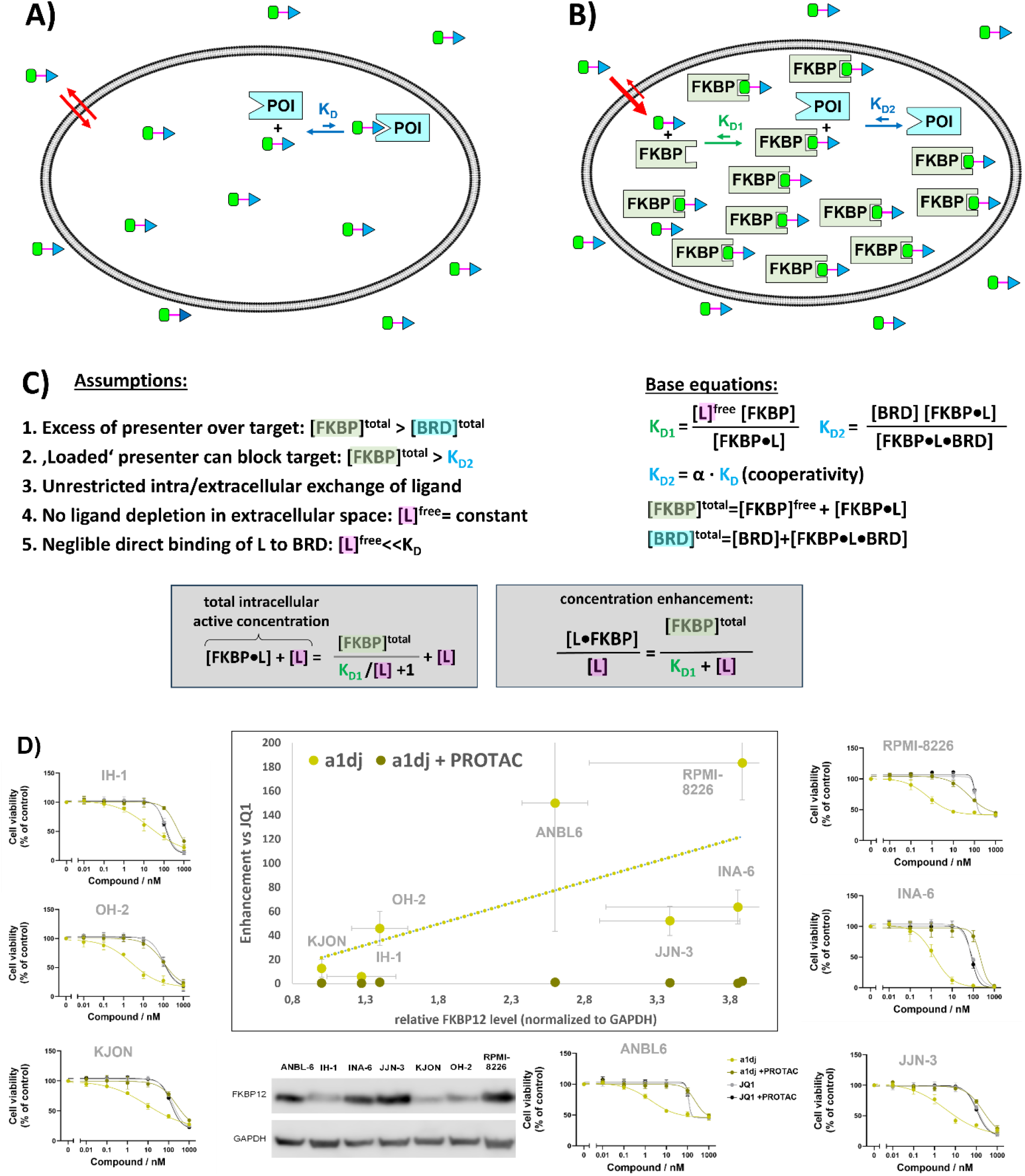
FKBP12-assisted intracellular trapping (CellTrap effect) assuming unrestricted equilibration of unbound drug between free extra- and intracellular drugs and neglecting sub-cellular partitioning of the presenter protein. **A)** Classical ‘binary’ pharmacology, driven by the affinity to the protein of interest. **B)** Presenter protein-assisted pharmacology driven by K_D1_, K_D2_, and the concentration of the presenter protein. Red arrows reflect intracellular drug accumulation due to trapping by intracellular FKBP12. **C)** Key numbers and concentration relationships of presenter protein-assisted pharmacology, exemplified for FKBP12 and BRD4. L represents the FKBP12-BRD4 bifunctional ligand. **D)** Cell toxicity of a1dj (yellow) in a panel of myeloma cell lines in the presence or absence of FKBP12 PROTAC. Toxicity is substantially enhanced compared to JQ1 in a FKBP12-dependent manner and the magnitude of enhancement correlates with FKBP12 expression levels (middle graph). FKBP12 and GAPDH levels were quantified from Western Blots (representative experiment shown in the bottom, see also SI Fig. 10C-E).

**Fig 8.**
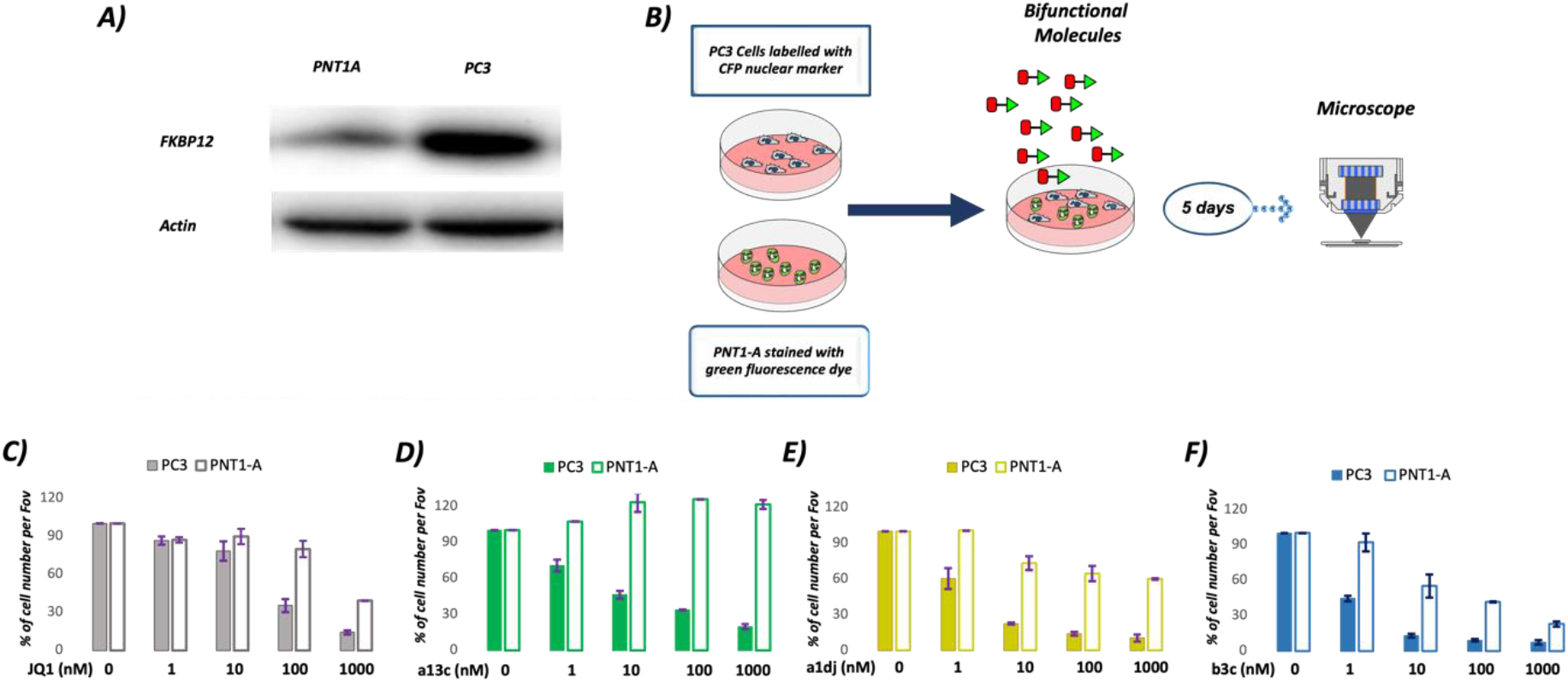
Selective cytotoxicity towards PC3 carcinoma cells over non-transformed PNT1A cells. **A**) Schematic representation of the co-culture live cell microscopy experiment workflow. **B**) Western blot analysis showing FKBP12 protein levels in PNT1-A and PC3 cell lines. **C**–**F**) Cell count analysis in PNT1-A and PC3 cells following treatment for five days with various concentrations of (**C**) JQ1, (**D**) a13c, (**E**) a1dj, and (**F**) b3c. Error bars represent the mean ± standard deviation (SD) from two independent live cell microscopy experiments.

To validate the requirement of ternary complex formation for BRD4 inhibition, we depleted FKBP12 in MV4-11 cells using a specific PROTAC before performing viability assays^24^ (**SI Fig. 7**). As expected, JQ1-induced decreases in cell viability were independent of FKBP12 status (Fig. 5A). BRD4-FKBP12 bifunctionals, however, lost their potency in the absence of FKBP12, as indicated by a strong increase of measured IC50 values for cell survival in the absence of FKBP12 (Fig. 5B-D and **SI Fig. 8A**). This effect was most striking for a1dj, where the ratio of IC50 values in the presence and absence of FKBP12 exceeded three orders of magnitude (Tab. 1).

To corroborate that the observed effect of BRD4-FKBP12 bifunctionals on cell viability is based on BRD4 inhibition, we quantified the expression of the known BRD4 target gene MYC at concentrations where we previously observed cytotoxic effects. As expected, JQ1 induced moderate downregulation of MYC at 10 nM and stronger effects at 100 nM, independent of FKBP12 status (Fig. 5E). Again, BRD4-FKBP12 bifunctionals exhibited stronger repression of MYC than JQ1 (Fig. 5F-H and **SI Fig. 8C**), with b3c inducing notable effects at concentrations as low as 1nM (Fig. 5H). Importantly, the reduction in MYC expression was attenuated by FKBP12 depletion for the bifunctionals. Due to its high efficacy, rescue of MYC expression by b3c was restricted to 1nM. At 10 and 100 nM, this compound induced MYC downregulation even after depletion of FKBP12, which was consistent with the previously observed FKBP12-independent decrease of cell viability at these concentrations (Fig. 5D).

Next, we tested cell viability and MYC expression in THP1 cells, which displayed noticeably lower FKBP12 levels than MV4-11 cells (**SI Fig. 7**). In THP1 cells, JQ1 showed comparable cytotoxic activity as in MV4-11 (SI Fig. 9A, IC50 THP1: 70 nM; IC50 MV4-11: 34 nM). The efficacies of FKBP12-BRD4 bifunctionals, however, were strikingly lower in THP1 compared to MV4-11 (10-60 fold), with a13c being even less effective than JQ1 (**SI Fig. 8B** and **SI Fig. 9 B-D**). Moreover, depleting FKBP12 led to a more modest decrease in IC50 values for all bifunctionals tested. Interestingly, downregulation of MYC upon BRD4 inhibition was more pronounced in these cells compared to MV4-11 (**SI Fig. 8D** and **SI Fig. 9E-H**), with similar FKBP12-dependency as observed before.

The observed residual activity of our bifunctional compounds in FKBP12-depleted cells was stronger than predicted from biochemically measured binding to BRD4 alone (see Table 1). To explore if this was due to remaining low levels of FKBP12, we turned to a previously established FKBP12 knock-out in the human myeloma cell line INA-6 (Fig. 6A, see Methods for details). INA-6 cells showed moderate JQ1 sensitivity independent of the FKBP12 status (Fig. 6B). As previously observed for MV4-11 and THP1, FKBP12-BRD4 bifunctionals showed increased efficacies in wild-type cells, reaching sub-nanomolar IC50 values for a1dj. In knock-out cells, however, cytotoxic effects were hardly observable for a13c and a1dj. Only the strong BRD4 binder b13c induced reduced cell viability at concentration of 100 nM and higher in FKBP12-negative cells.

Our results so far demonstrate that FKBP12-BRD4 bifunctionals inhibit BRD4 function in cells depending on the FKBP12 status. Notably, the efficacies of our compounds were orders of magnitude higher compared to JQ1. This could not be explained by enhanced biochemical binding, as the FKBP12-assisted biochemical affinities were only 2-5-fold higher (Tab. 1). We hypothesized that the enhanced efficacy in cells is based on an ‘upstream’ binding of bifunctional compounds to the highly abundant protein FKBP12 upon cell entry (Fig. 7B vs 7A). This shifts the equilibrium between extra- and intracellular compound concentrations and thereby increases the effective intracellular concentration. FKBP12 binding therefore acts as a cellular trap, which enriches bifunctional compounds in the cells and leads to higher intracellular concentrations compared to monofunctional compounds such as JQ1.

To functionally test this hypothesis, we first developed a mathematical model reflecting intracellular FKBP12 binding and BRD4 inhibition. This model accounted for enhanced cooperative binding of ligands when prebound to FKBP12 (KD2 vs KD, Fig. 7C) and predicted an increased intracellular concentration of ligand species available for BRD4 inhibition (free ligand + FKBP12-bound ligand), which positively correlates with FKBP12 levels. In consequence, effective concentrations were predicted to be reachable at very low concentrations, which were dominated by the FKBP12-ligand complex as the active species **(SI Fig. 10)**.

To validate this, we determined FKBP12 levels in a panel of human myeloma cells and measured cell survival upon treatment with JQ1 and a1dj (Fig. 7D). Indeed, we observed higher increases of a1dj efficacies compared to JQ1 in cells with higher FKBP12 levels. Importantly, the cytotoxicity of a1dj was almost identical to JQ1 upon depletion of FKBP12 in all cell lines tested (Fig. 7D). Such correlation between enhanced efficacies and FKBP12 levels were also observed for a13c and, to a lesser extent, for b3c **(SI Fig. 11)**.

### Selective cytotoxicity of a13c in co-cultured prostate-derived cell lines

The cellular trapping of bifunctional compounds and their dependency on FKBP12 levels suggest that they can be employed to preferentially deplete cells with high FKBP12 levels from co-cultures, offering the potential for an unforeseen therapeutic window. To test this hypothesis, we turned to prostate cells were strong differences in FKBP12 levels were observed for non-transformed PNT1A cells compared to PC3 carcinoma cells (Fig. 8A). We first measured cell survival upon treatment with JQ1 and FKBP12-BRD4 bifunctionals. PNT1A cells showed in general only modest sensitivity to BRD4 inhibition **(SI Fig. 12)**. As FKBP12 expression was already low in these cells, we observed almost no alteration of cell survival upon FKBP12 depletion. Noticeably, a13c did not affect PNT1A viability even at concentrations of up to 1µM. In PC3 cells, however, BRD4 inhibition induced strong cytotoxic effects, which were FKBP12-dependent for all tested bifunctionals **(SI Fig. 13)**.

To investigate selective cell killing according to FKBP12 levels, we performed co-culture experiments of PNT1A and PC3 cells. We applied differential fluorescent labels to quantify both cell types after treatment with JQ1, a13c, a1dj and b3c (Fig. 8B). For JQ1, we observed preferential killing of carcinoma cells at high concentrations as expected from viability measurements of individual cell types (Fig. 8C, **SI Fig. 12** and **SI Fig. 13**). The concentration range inducing selective cell killing was enlarged for the bifunctionals a1dj and b3c, showing noticeable effects on PC3 cells at 1 to 10nM (Fig. 8E-F). As PNT1A cells were unaffected by a13c treatment at all concentrations, the most striking effect was observed for this bifunctional: the population of PC3 cells could be efficiently depleted from the co-culture while PNT1A cells continued to proliferate (Fig. 8D). Notably, the selective cytotoxic effect of bifunctionals was reversed upon FKBP12 depletion (**SI Fig 14**). This demonstrates that even moderate differences in endogenous presenter protein levels can be used advantageously to induce cell specific toxicity, offering unprecedented opportunities for enhanced therapeutic windows to treating human malignancies.

## Discussion

Drugs that induce proximity between a target protein and an endogenous effector or presenter protein offer numerous unprecedented opportunities for refined pharmacology. Synthetically, proximity-based drugs can be readily implemented by “Lego-like” assembly of heterobifunctional molecules.

We show for BRD4 and FKBP12 as a target-presenter pair that it was possible to readily obtain heterobifunctional molecules with positive cooperativity. For this, multiple structural solutions exist, three of which we could resolve at atomic detail. Whether the propensity for positive cooperativity is specific to FKBP12, which has a legacy as a one of nature’s preferred presenter, remains to be seen.

Our results also show that heterobifunctionals, even in their most basic setup, can readily provide selective ligands that can distinguish between highly similar protein binding pockets, here BRD4^BD1^ vs BRD4^BD2^. This likely is due to the large FKBP12-a13c binary complex, which is predicted to engage an extended surface on BRD4^BD1^, incl. residues which differ from BRD4^BD2^. Similar effects have repeatedly been observed for PROTACs^30,31^ and are likely a general feature of proximity-based pharmacology. Like for PROTACs, the occurrence of selectivity is hard to predict and has to be explored experimentally, underscoring the importance of rapidly exploring linker diversity and linker attachment chemistry.

A striking and in this magnitude unexpected finding was the dramatically enhanced potencies of the bifunctionals compared to JQ1. Specifically, the cellular potencies were orders of magnitude higher for the bifunctionals than expected by their biochemical BRD4 affinities. This was strictly dependent on FKBP12 and correlated with the expression level of the presenter protein. We attribute this effect to the intracellular trapping of heterobifunctionals by the highly expressed FKBP12, which we termed ‘CellTrap’ effect (Fig. 7). Assuming an undepletable extracellular reservoir and unrestricted equilibration of free ligand, the latter will unidirectionally diffuse into cells and there bind to FKBP12 until the unbound FKBP12 levels get depleted and approach KD1 (Fig. 7B) This follows saturation binding and at [L] = KD1 half of the intracellular FKBP12 pool is ligand-bound. For [L] < KD1, intracellular concentration enhancement scales linearly with presenter protein levels and anti-proportionally with the affinity of the presenter protein (Fig. 7C). At these sub-saturating concentrations of the ligand with respect to the presenter protein, the presenter protein is only partially occupied. This is the preferred concentration range for presenter protein-mediated pharmacology, as side effects due to presenter protein blockade are minimized.

The key corollary in this model is that the CellTrap effect is most pronounced for highly expressed proteins combined with ultrahigh-affinity ligands. This can explain the extraordinary cellular potencies observed in this work. Intracellular buffering has been proposed to contribute to the efficacies of the FKBP12-based molecular glues FK506 and Rapamycin^1,32^. Intracellular enrichment was also observed for CypA-based molecular glues^33^ and in drug sequestering strategies^32,34^, but never at this magnitude.

The striking potency enhancement enabled by the CellTrap effect resulted in a large concentration window where the efficacy of the heterobifunctionals were strictly dependent on FKBP12 as a presenter protein. Moreover, FKBP12-high expressing cell lines were clearly more susceptible to FKBP12-BRD4 heterobifunctionals compared to FKBP12-low expression cell lines. This offers an unprecedented layer of cell type specificity, driven by the presenter protein. We demonstrated this by the preferential killing of PC3 cells over non-transformed PNT1-A cells in co-culture experiments. This is conceptually similar to the RIPTAC concept^6^, which however has so far only been shown with overexpressed presenter proteins. Our results demonstrate that even moderate differences in presenter protein expression can be used to achieve cell-type specificity.

Taken together, our findings show that remarkable cellular potencies and presenter protein-dependencies can be achieved with heterobifunctionals containing high-affinity ligands for high-expressing proteins. We postulate that this may compensate for unfavorable physicochemical properties imparted by the high molecular weight. Indeed, their limited baseline activity in the absence of the CellTrap effect contributes to the observed cell-type specificity, potentially creating favorable therapeutic windows. Furthermore, like other bifunctional modalities, the co-engagement of two proteins enables otherwise difficult-to-achieve selectivity and allows to differentiate cells based on the co-engaged presenter protein. Compared to PROTACs, non-degrading heterobifunctionals might be the preferred choice when degradation of the entire target proteins is difficult, e.g. for poorly degradable target proteins, or not desirable, e.g. due to excessive toxicity. Although not observed in this work, we note that engaging the target with a presenter-compound pre-complex might be functionally different than blocking the target with a small molecule alone. We expect our findings to be similarly relevant to molecular glues in addition to heterobifunctional molecules. Our findings support CellTrap-assisted heterobifunctionals as therapeutically attractive options, underscore the potential of highly expressed proteins as presenters, and call for developing high affinity ligands for them.

## Methods

### Fluorescence polarization assays

Binding of compounds to FKBP12 and to the isolated BD1 and BD2 domains of BRD4 was determined in a competitive fluorescence polarization assay. For FKBP12 the assay was performed as described by Kozany et al. using the tracer developed by Pomplun et al^25,28^. A similar assay routine was developed for BRD4. In brief, a serial dilution of the compounds in DMSO was prepared using a Biomek FX^P^ pipetting robot (Bekman Coulter). The compounds were transferred to black 384-well plates (Greiner 781076) filled with assay buffer (20 mM HEPES-NaOH, pH 8.0, 150 mM NaCl, 0.002% TX-100 and 0.05% BSA) or assay buffer supplemented with FKBP12 (final concentration 15 µM) and the tracer JQ1-FL (final concentration 2 nM) as well as the respective BD1 or BD2 domain of BRD4 (final concentration 200 nM) were added. After equilibration of 30 min at room temperature the assay plates were read using a Tecan Spark plate reader using excitation/emission wavelengths of 535/590 nm (FKBP12) or 485/535 nm (BRD4). Data analysis was performed with GraphPad Prism 8 software using the fitting equation developed by Wang^35^.

### HTRF assays

Compound induced ternary complex formation was followed by a HTRF assay. This assay relies on the establishment of an HTRF signal between a terbium-labeled antibody binding to the His6-tag of the respective BD1 or BD2 domain of BRD4 and EGFP-labeled FKBP12 upon close proximity. Therefore, a serial dilution of the respective compound in EGFP-FKBP12 (final concentration 5 µM) containing assay buffer (20 mM HEPES-NaOH, pH 8.0, 150 mM NaCl, 0.002% TX-100 and 0.05% BSA) was placed in a white 384-well assays plate (ProxiPlate 384-shallow well Plus, Revvity) and a mixture of BRD4^BD1^ or BRD4^BD2^ (final concentration 100 nM) and 2-fold terbium-labeled anti-6xHis antibody (Revvity) was added. After equilibration of 30 min at room temperature the fluorescence intensity at 535 and 620 nm upon excitation at 340 nm was determined on a Tecan Spark plate reader using a lag time of 150 µs and an integration time of 500 µs. The HTRF signal was determined by forming the ratio 535 nm/620 nM and the data was fitted to a binary binding model using GraphPad Prism 8 software.

### Protein crystallization

In order to generate the FKBP12:**a1d**:BRD4^BD1^ and the FKBP12:**b3c**:BRD4^BD1^ complexes for protein crystallization trials, 2 ml of a solution containing 140 µM BRD4^BD1^, 140 µM **a1d** or **b3c** and 150 µM FKBP12 were prepared in 20 mM HEPES pH 7.5, 150 mM NaCl, 5% DMSO and concentrated to about 400 µl. The samples were separated on a size exclusion column (Superdex 75 10/100 GL increase, Cytiva) and fractions containing the ternary complex were pooled, concentrated to about 10 mg/ml, flash frozen in liquid nitrogen and stored at -80°C.

The crystallization of the FKBP12:**a1d**:BRD4^BD1^ complex was performed at room temperature using the hanging drop vapour-diffusion method, equilibrating mixtures of 1 µl protein complex and 1 µl reservoir against 500 µl reservoir solution consisting of 22% PEG3350, 0.2 M NaCl and 0.1 M Tris-HCl pH 8.5. Crystals were fished, cryo protected with reservoir solution supplemented with 10% glycol and flash frozen in liquid nitrogen.

The crystallization of the FKBP12:**b3c**:BRD4^BD1^ complex was performed at room temperature using the sitting drop vapour-diffusion method, equilibrating mixtures of 0.5 µl protein complex and 0.5 µl reservoir against 30 µl reservoir solution consisting of 25% PEG3350, 0.2 M NH4SCN and 0.1 M Tris-HCl pH 8.8. Crystals were fished, cryo protected with reservoir solution supplemented with 10% glycol and flash frozen in liquid nitrogen.

The crystallographic experiments were performed at the BL14.1 at the Helmholtz-Zentrum BESSYII synchrotron, Berlin, Germany^36^ and the beamline ID23-1 of the European Synchrotron Radiation Facility (ESRF) in Grenoble, France and the raw data can be accessed at https://doi.org/10.15151/ESRF-DC-2127908021^37^. Detailed information on the structure solution and the crystallographic data are provided in the supplemental information.

### Cell lines

PC3 cells were kindly provided by Dr. G. Scheiner-Bobis (Justus Liebig University Giessen, Germany), THP-1 and MV4-11 acute myeloid leukemia cells by Dr. S. Knapp (Goethe University Frankfurt, Germany). PNT1-A prostate-derived normal epithelial cells and RPMI-8226 HMCL were purchased from ATCC. HEK293T and PC3 cells were cultured in Dulbecco’s Modified Eagle Medium (DMEM; Gibco, Darmstadt, Germany) supplemented with 10% fetal bovine serum (FBS) and 1% penicillin-streptomycin (P/S). PNT1-A, THP-1 and MV4-11 cells were maintained in RPMI-1640 medium (Gibco). THP-1 cells were supplemented with 15% FBS, while MV4-11, PNT1-A, ANBL-6, INA-6 and JJN-3 cells were grown in 10% FBS. Media included 1% P/S (Merck, Darmstadt, Germany) and 1% GlutaMAX (Gibco).

The human myeloma cell lines (HMCL) ANBL-6 (kind gift from Dr. D. Jelinek, Mayo Clinic, Rochester, MN, USA), INA-6 (kind gift from Dr. M. Gramatzki, University of Erlangen-Nurnberg, Germany), and JJN-3 (kind gift from Dr. J. Ball, University of Birmingham, UK) were grown in 10% fetal bovine serum (FBS) with 1 ng/mL interleukin (IL)-6 (Gibco, Thermo Fisher Scientific, Waltham, MA, USA) added for ANBL-6 and INA-6. RPMI-8226 (ATCC, Rockville, MD, USA) was grown in 15% FBS in RPMI. IH-1, KJON, and OH-2 were established in our laboratory (PMID: 39567630)^38^ and were grown in 10% human serum (HS, Department of Immunology and Transfusion Medicine, St. Olav’s University hospital, Trondheim, Norway) in RPMI, supplemented with IL-6 (2 ng/mL). For experiments, 2% HS in RPMI was used, with IL-6 (1 ng/mL) added for all cell lines, except JJN-3 and RPMI-8226.

### NanoBRET

The assay was performed as described previously [PMID: 38637703]^39^. In brief, full-length BRD4 was obtained as plasmid cloned in frame with a terminal NanoLuc-fusion. Plasmids were transfected into HEK293T cells using FuGENE HD (Promega, E2312), and proteins were allowed to express for 20 h. Serially diluted inhibitor and BRD Tracer-02 (Promega, TracerDB ID: T000022) at the Tracer KD concentration taken from TracerDB (tracerdb.org [PMID: 38969708])^40^ was pipetted into white 384-well plates (Greiner 781207) using an ECHO acoustic dispenser (Labcyte). The corresponding protein-transfected cells were added and reseeded at a density of 2 × 10^5^ cells/mL after trypsinization and resuspending in Opti-MEM without phenol red (Life Technologies). The system was allowed to equilibrate for 2 h at 37 °C/5 % CO2 prior to bioluminescence resonance energy transfer (BRET) measurements. To measure BRET, NanoBRET NanoGlo Substrate was added as per the manufacturer’s protocol, and filtered luminescence was measured on a PHERAstar plate reader (BMG Labtech) equipped with a luminescence filter pair (450 nm BP filter (donor) and 610 nm LP filter (acceptor)). Competitive displacement data were then graphed using GraphPad Prism 9 software using a normalized 3-parameter curve fit with the following equation: Y = 100/(1 + 10(X − log IC50)).

### Western Blot

To assess FKBP12 levels, 5 × 105 PC3, PNT1-A, and HMCL cells were seeded into 6-cm plates and cultured for two days. Suspension cell lines MV4-11 and THP-1 were cultured in T25 flasks under similar conditions. For FKBP12 degradation, all cell lines were treated with an FKBP12-specific PROTAC (5a1) 200 nM for either 6 hours or 4 hours in the case of HMCL. Following treatment, cells were washed with PBS, harvested, and lysed for 30 min on ice. The lysis buffer used contained 1% IGEPAL CA-630 (Sigma Aldrich, St Louis, MO, USA), 150 mM NaCl, 50 mM Tris–HCl (pH 7.5), protease inhibitor cocktail Complete Mini (Roche, Basel, Switzerland), 50 mM NaF, and 1 mM Na3VO4. Protein concentrations were determined using the Bradford assay. 20 µg protein was loaded on 10% SDS-PAGE gels alongside a pre-stained molecular weight marker (Bio-Rad, Feldkirchen, Germany). Proteins were transferred to either nitrocellulose or PVDF membranes (Merck, Darmstadt, Germany) using a wet transfer system at 100 V for 1.5 hours. Membranes were blocked with 5% skim milk in TBS-T (TBS with 0.1% Tween-20) for 1 hour at room temperature and subsequently incubated overnight at 4°C with FKBP12 primary antibodies. Primary antibodies used were FKBP12 (RRID: AB_2102847, #sc-133067, Santa Cruz, TX, USA), FKBP12 (HUABIO: HBO-ET1703-55, BIOZOL, Eching, Germany), Actin (SIGMA-ALDRICH, A5316, Germany), and GAPDH (RRID: AB_2107448, Cat# Ab8245, Abcam, Cambridge, UK). After three washes with TBS-T, membranes were incubated with HRP-conjugated secondary antibodies for 1 hour at room temperature. Protein bands were detected using an ECL kit and visualized with either Fusion FX gel imaging system or ChemiDoc Imager (Bio-Rad) and quantified using Image Studio software (LI-COR Biosciences, Cambridge, UK).

### Quantitative RT-PCR

Total RNA was extracted using the Monarch Total RNA Isolation Kit. RNA concentration and purity were assessed using a NanoDrop spectrophotometer. cDNA synthesis was performed using reverse transcription. Quantitative PCR (qPCR) was conducted on an Applied Biosystems StepOne Real-Time PCR system using the following primers: MYC forward: TTGGTGAAGCTAACGTTGAGG; MYC reverse: TATTCTGCCCATTTGGGGACA, Actin forward: TAGAAGCATTTGCGGTGGACGATG; Actin reverse: GGCACCCAGCACAATGAAGATCAA-

### Cell viability assay

Cell viability and cytotoxicity were assessed using the CellTiter-Glo Luminescent Viability Assay. HMCL, PC3, and PNT1-A cells were seeded into 96-well plates at densities of 1 × 10^4^, 1 × 10^3^, and 4 × 10^3^ cells per well, respectively. For FKBP12 degradation experiments, the cells were pre-treated with an FKBP12-specific PROTAC (5a1) 200 nM for either 6 hours or 4 hours in the case of HMCL, before adding the compounds or corresponding amount of DMSO, each in a total volume of 100 µl full growth medium. The plates were incubated for either 5 consecutive days or two days for HMCL cells. Suspension cell lines MV4-11 and THP-1 were cultured in T25 flasks for 2 days prior to the experiment. FKBP12 degradation was induced following the same procedure described for adherent cells. After incubation, the cells were counted, and 4 × 10^3^ cells were seeded per well in 96-well plates. Experimental treatments, including the appropriate concentrations of various compounds and necessary controls, were added to each well in 100 µl full growth medium. After the incubation period, The full growth medium containing the compounds was aspirated from all cell lines. The CellTiter-Glo 2.0 reagent (Promega, Madison, WI, USA) was diluted 1:1 with PBS and added to each well. For MV4-11 and THP-1 suspension cells, the reagent was instead diluted 1:1 with the existing full growth medium in each well. Luminescence was measured using a TECAN microplate reader (TECAN, Switzerland).

### CRISPR/Cas9 knock-out of *FKBP1A*

INA-6 FKBP1A KO cells were made previously^41^. In brief, CRISPR sgRNAs were designed using Benchling (https://www.benchling.com/). Oligo pairs encoding sgRNA targeting *FKBP1A* (GGGCGCACCTTCCCCAAGCG) or non-targeting control (TGAATCGTAACCTCGCCATT) were annealed and ligated into lentiCRISPR v2 (gift from Feng Zhang, Addgene plasmid #52961; http://n2t.net/addgene:52961; RRID:Addgene_52961). The plasmids were cotransfected with third generation lentiviral packaging plasmids using Genejuice (Novagen, Merck Life Science AS, Oslo, Norway) in 293T packaging cells (Open Biosystems, Thermo Fisher Scientific). The filtered supernatant was used to generate the knockout cells by treating INA-6 with lentiviral particles in the presence of polybrene (8 µg/mL). Fresh medium was added after 4 h. Puromycin (0.5 µg/mL) was added after 48 h to select for cells with successful integration. Loss of *FKBP1A* was confirmed with immunoblotting and the cells were expanded and used for experiments.

### Live cell microscopy

PC3 prostate cancer cells and PNT1-A normal prostate epithelial cells were co-cultured and analyzed using live-cell microscopy. PC3 cells were labeled with a CFP fluorescent nuclear marker, while PNT1-A cells were stained with Cytopainter Green dye. Equal numbers of both cell types were seeded into 24-well plates under the specified experimental conditions. After 5 days of incubation, the culture medium was replaced with microscopy medium to optimize imaging conditions. Images were acquired using the Nikon Eclipse live-cell microscopy system. Cell segmentation and quantification were performed using a combination of CELLPOSE and custom MATLAB scripts.

## Acknowledgements

Diffraction data have been collected on BL14.1 at the BESSY II electron storage ring operated by the Helmholtz-Zentrum Berlin, Germany and on ID23-1 at the European Synchrotron Radiation Facility (ESRF) in Grenoble, France. We thank the HZB for the allocation of synchrotron radiation beamtime and we would particularly like to acknowledge the help and support of Manfred Weiss and the whole MX team during the experiment. We would like to thank the staff of the ESRF and EMBL Grenoble for assistance and support in using beamline ID23-1 under proposal number MX-2555.

