## Supporting Information for "Heterobifunctional Protein Binders Enable Cell Type-Specific Killing Through In-cell Enrichment"

1. **SI Table 1: Characterization of BRD4-FKBP12-bifunctional ligands.** Binding of Straps to purified BRD4 BD1 or BD2 bromo domains was determined in a competitive fluorescence polarization assay in absence or presence of 15 µM FKBP12 using 2 nM of a fluorescently labeled JQ1-analog as a tracer. Binding of Straps to FKBP12 was determined in a competitive fluorescence polarization assay. NanoBRET data was determined in intact cells by expression of NanoLuc-fusion proteins and using fluorescently labeled FKBP1[2] or BRD4 ligands.

| Compound | BRD4 ligand | Linker | FKBP12 ligand | Binding to FKBP12 | Binding to BRD4 BD1 -/+ FKBP12 | | Binding to BRD4 BD2 -/+ FKBP12 | | NanoBRET FKBP12 | NanoBRET BRD4 |
|---|---|---|---|---|---|---|---|---|---|---|
| | | | | Kd / nM | Kd / nM | α | Kd / nM | α | $IC_{50}$/ nM | $IC_{50}$/ nM |
| JQ1 | | | | | 46±9<br>47±9 | 1.0 | 90±24<br>90±18 | 1.0 | | |
| a1d | 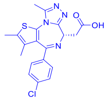 | 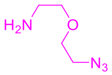 | 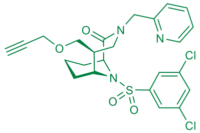 | 1.0 | 680±120<br>25±6 | 27 | 440±90<br>57±14 | 7.7 | 14 | 820 |
| a1c | 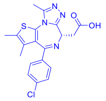 | 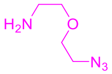 | 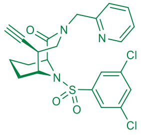 | 1.0 | 1300±200<br>3900±500 | 0.3 | 590±110<br>1500±260 | 0.4 | 22 | 940 |
| a2c | 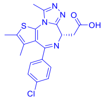 | 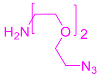 | 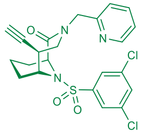 | 0.6 | 440±70<br>250±40 | 1.8 | 510±110<br>1100±140 | 0.5 | 17 | 620 |

| | | | | | | | | | | |
|---|---|---|---|---|---|---|---|---|---|---|
| a1i | 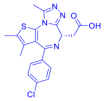 | 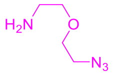 | 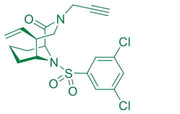 | 10<br> | 1800±270<br>3900±500 | 0.5 | 910±160<br>2900±400 | 0.3 | 200 | 560 |
| a2i | 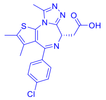 | 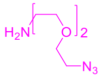 | 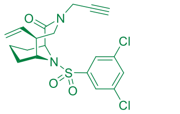 | 8.7<br> | 330±40<br>290±30 | 1.1 | 140±30<br>960±120 | 0.1 | 180 | 790 |
| a1h | 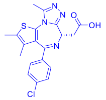 | 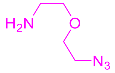 | 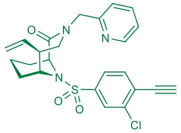 | 50<br> | 100±18<br>1700±200 | 0.06 | 52±13<br>1100±200 | 0.05 | 530 | 270 |
| a2h | 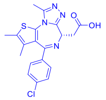 | 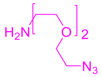 | 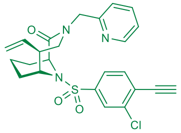 | 2.4<br> | 210±18<br>2400±200 | 0.09 | 46±10<br>1300±200 | 0.04 | 170 | 300 |
| a1g | 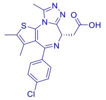 | 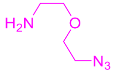 | 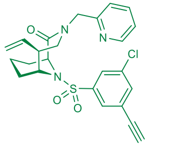 | 46<br> | 2800±500<br>>20000 | | 1700±300<br>9000±2000 | 0.2 | 230 | 1500 |
| a2g | | | | 76 | 1200±140 | 0.4 | 580±90 | 0.4 | | 3066 |

| | | | | | | | | | | |
|---|---|---|---|---|---|---|---|---|---|---|
| | 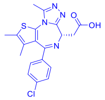 | 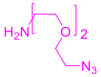 | 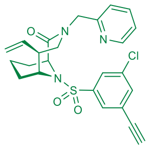 | | 3100±30<br>0 | | 1500±20<br>0 | | | |
| a4d | 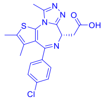 | 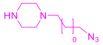 | 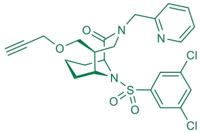 | 1.1 | 500±60<br>380±50 | 1.3 | 1200±20<br>0<br>1200±10<br>0 | 1 | 89 | none |
| a5d | 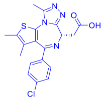 | 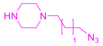 | 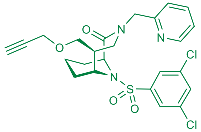 | 1.2 | 740±100<br>300±50 | 2.5 | 1100±20<br>0<br>600±100 | 1.8 | 93 | 1080 |
| a6d | 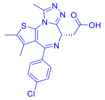 | 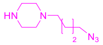 | 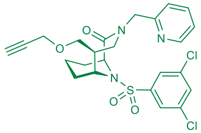 | 0.9 | 600±80<br>400±50 | 1.5 | 1400±30<br>0<br>800±200 | 1.8 | 141 | 787 |
| a7c | 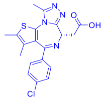 | 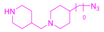 | 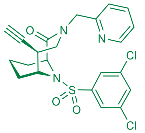 | 3.5 | 520±80<br>180±30 | 2.9 | 700±100<br>900±200 | 0.8 | 138 | 665 |
| a7d | 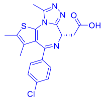 | 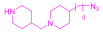 | 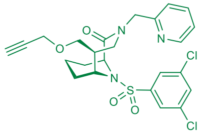 | 2.3 | 680±70<br>310±50 | 2.2 | 700±100<br>660±90 | 1 | 209 | 1115 |

| | | | | | | | | | | |
|---|---|---|---|---|---|---|---|---|---|---|
| a8c | 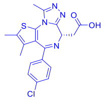 | 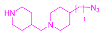 | 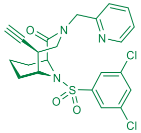 | 2.1 | 170±20<br>150±20 | 1.1 | 190±30<br>160±30 | 1.2 | 178 | 541 |
| a8d | 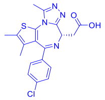 | 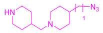 | 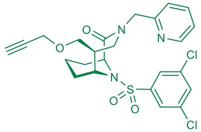 | 1.8 | 160±30<br>84±16 | 1.9 | 130±30<br>120±30 | 1 | 228 | 357 |
| a15d | 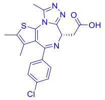 | 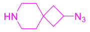 | 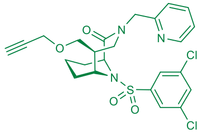 | 2.6 | 3000±700<br>420±60 | 7.1 | 5400±1300<br>520±80 | 10 | 157 | 5218 |
| a13c | 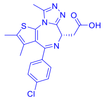 | 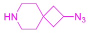 | 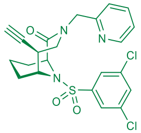 | 1.7 | 5700±900<br>20±6 | 285 | 2700±400<br>1600±300 | 1.7 | 76 | 9057 |
| a1dj | 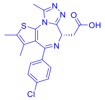 | 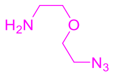 | 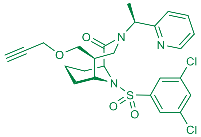 | 0.4 | 810±90<br>32±9 | 25 | 500±100<br>70±20 | 7 | 2.6 | 653 |
| a1dk | 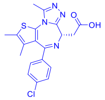 | 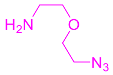 | 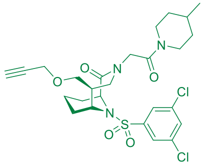 | 0.47 | 520±90<br>200±40 | 2.6 | 320±60<br>800±100 | 0.4 | 9.1 | 1189 |

| | | | | | | | | | | |
|---|---|---|---|---|---|---|---|---|---|---|
| a1e | 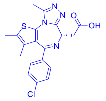 | 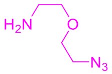 | 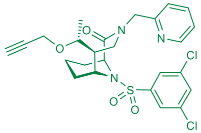 | 8.9 | 70±10<br>48±7 | 1.5 | 33±9<br>60±10 | 0.6 | 103 | 390 |
| a1f | 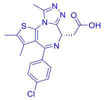 | 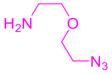 | 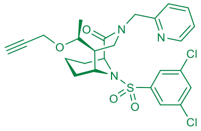 | 1.6 | 180±20<br>140±14 | 1.3 | 50±14<br>230±30 | 0.2 | 24 | 560 |
| b3d | 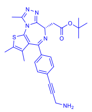 | 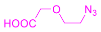 | 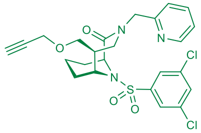 | 1.0 | 68±9<br>200±30 | 0.3 | 110±30<br>1100±100 | 0.1 | 15 | 266 |
| b3c | 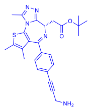 | 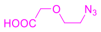 | 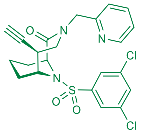 | 3.7 | 400±100<br>8±6 | 50 | 380±90<br>9±8 | 42 | 29 | 194 |
| a10c | 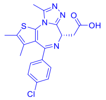 | 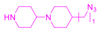 | 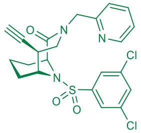 | 0.3 | 120±30<br>900±100 | 0.1 | 170±30<br>1800±300 | 0.09 | 288 | 497 |
| a10d | 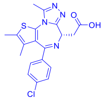 | 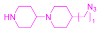 | 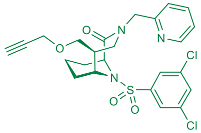 | 1.6 | 140±30<br>100±30 | 1.4 | 170±30<br>80±20 | 2.1 | 241 | none |
| a9c | | 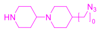 | | 2.2 | 280±40 | 1.6 | 310±40 | 0.5 | 107 | 155 |

| | | | | | | | | | | |
|---|---|---|---|---|---|---|---|---|---|---|
| | 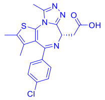 | | 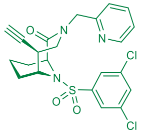 | | 180±20 | | 650±90 | | | |
| a9d | 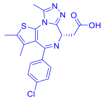 | 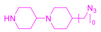 | 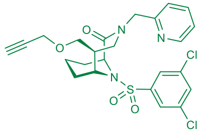 | 1.7 | 340±40<br>140±10 | 2.4 | 500±90<br>450±70 | 1.1 | 169 | 364 |
| a12c | 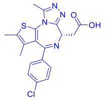 | 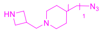 | 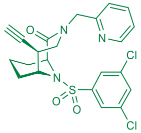 | 1.8 | 390±50<br>230±20 | 1.7 | 800±100<br>700±90 | 1.1 | 280 | 748 |
| a12d | 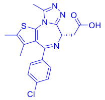 | 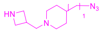 | 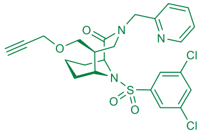 | 2.3 | 250±40<br>40±10 | 6.3 | 1100±10<br>0<br>590±70 | 1.9 | 406 | 514 |
| a11c | 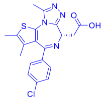 | 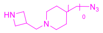 | 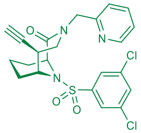 | 0.7 | 1000±20<br>0<br>190±30 | 5.3 | 2100±30<br>0<br>450±70 | 4.7 | 218 | 1229 |
| a11d | 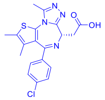 | 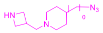 | 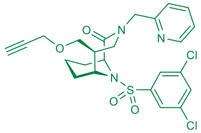 | 0.8 | 480±70<br>40±10 | 12 | 1250±15<br>0<br>700±100 | 1.8 | 165 | 687 |

### 2. Supplementary Figures

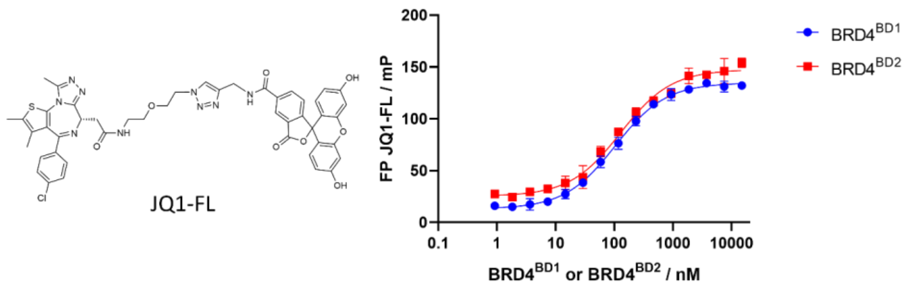

**SI Fig. 1. Characteristics of the fluorescence polarization assay for BRD4.** The binding of $BRD4^{BD1}$ and $BRD4^{BD2}$ to JQ1-FL was determined with a fluorescence polarization (FP) assay. Therefore, a serial dilution of $BRD4^{BD1}$ and $BRD4^{BD2}$ was added to 2 nM of JQ1-FL and the fluorescence polarization was determined. Data ponits are indicated as mean from three replicates and standard deviations are indicated as error bars. The solid lines represent data fitted to a binary binding model. The $K_d$-values were determined to be 100±6 nM for $BRD4^{BD1}$ and 130±10 nM for $BRD4^{BD2}$.

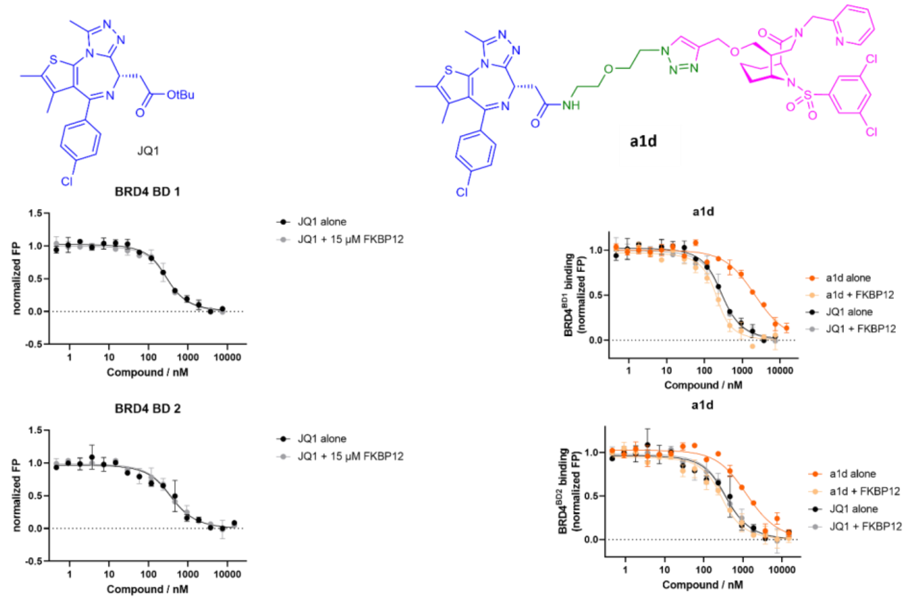

**SI Fig. 2: BRD4 binding of a1d.**

The binding of **a1d** alone or in the presence of 15 µM FKBP12 to BRD4[BD1] and BRD4B[D2] was determined in a competitive FP assay using a fluorescent JQ1 analog as tracer. Data ponits are indicated as mean from two replicates and standard deviations are indicated as error bars. The data was fitted to a competitive binding model and the fit is indicated as solid line.

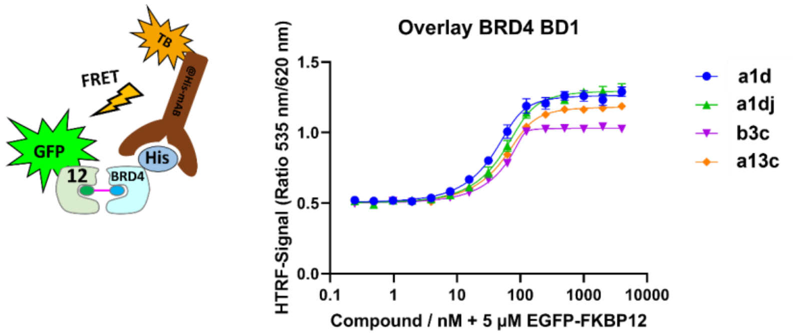

**SI Fig. 3: Ternary complex formation assessed by HTRF**

The ternary complex formation between EGFP-FKBP12 and His-BRD4$^{BD1}$ upon addition of a1d, a1dj, a13c and b3c was determined using a HTRF assay. Therefore a serial dilution of the respective compound was incubated with a mixture of 5 µM EGFP-FKBP12, 100 nM His-BRD4$^{BD1}$ (complexed with a Terbium-labelled antibody). Data points are indicated as mean from three replicates and standard deviations are indicated as error bars. The solid lines represent data fitted to a binary binding model.

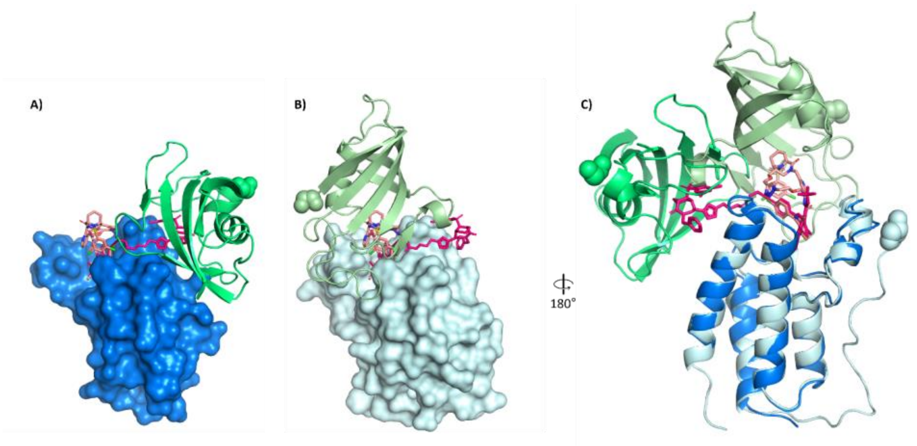

**SI Fig. 4. Ternary binding mode of b3c (9QW8) and comparison with a1d (9R5N). (A)** Ternary complex of $BRD4^{BD1}$ (marine blue surface), b3c (red pink sticks) and FKBP12 (green cartoon). The binding pose of a1d (pale magenta sticks) is shown superimposed from the FKBP12-a1d-$BRD4^{BD1}$ complex. **(B)** Ternary complex of $BRD4^{BD1}$ (pale surface), a1d (pale magenta sticks) and FKBP12 (pale green cartoon), view from the same perspective as in (A). The binding pose of b3c (red pink sticks) is shown superimposed from the FKBP12-b3c-$BRD4^{BD1}$ complex. **(C)** Superposition of the ternary complexes of b3c and a1d, with colors as in (A) and (B) and viewed as in main Fig. 4H (rotated by 180° compared to (A) and (B)).

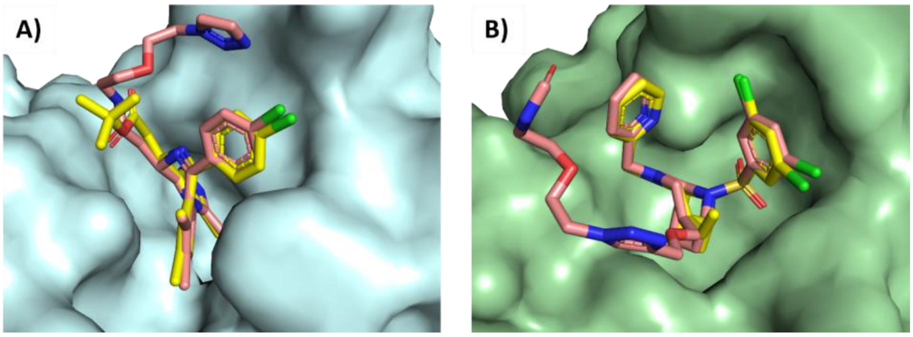

**SI Fig. 5. Binding modes of BRD4 & FKBP12 ligands are conserved: A)** JQ1 moiety of **a1d** in the FKBP12-BRD4$^{BD1}$ complex overlayed with JQ1 bound to BRD4$^{BD1}$ (PDB-ID 3MXF). The surface of BRD4$^{BD1}$ is colored cyan, compound **a1d** is colored pale pink and JQ1 is colored in yellow (FKBP12 and FKBP12 binding part of **a1d** omitted for clarity).

**B)** FKBP12-binding [4.3.1] bicyclic sulfonamide moiety of **a1d** in the FKBP12-BRD4$^{BD1}$ complex overlayed with a [4.3.1] bicyclic sulfonamide FKBP12 ligand bound to FKBP12 (PDB-ID 8CHL). The surface of FKBP12 is colored green, compound **a1d** is colored pale pink and the FKBP12 ligand is colored in yellow (BRD4$^{BD1}$ and JQ1 moiety of **a1d** omitted for clarity).

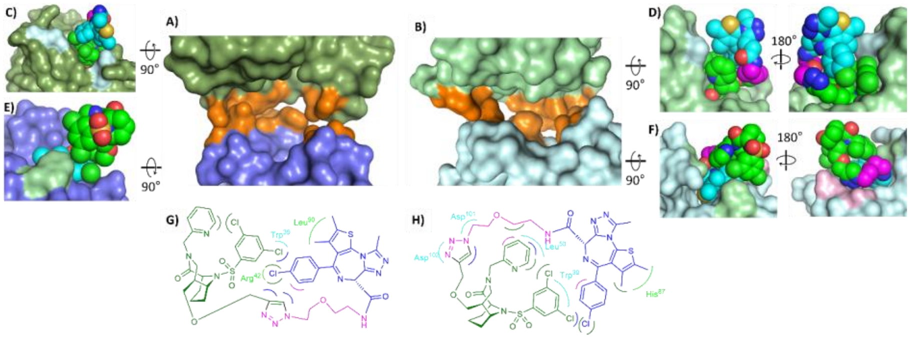

**SI Fig. 6. Analyis of ternary complex interactions.** FKBP12 and BRD4$^{BD1}$ are shown as light/dark green and cyan/dark blue surfaces, respectively. Compound **a1d** is shown as sphere model, with carbons of the FKBP12 ligands colored green, carbons of the BRD4 ligands are colored cyan, and carbons of the linker are colored magenta. The two different ternary complexes in the crystallographic unit are shown in light and dark colors, respectively. **A,B)** Amino residues engaged in direct protein-protein contacts are highlighted in orange. The ligand **a1d** has been omitted for clarity. **C,D) a1d** bound to FKBP12, BRD4$^{BD1}$ is not shown for clarity. FKBP12 residues contacting the BRD4 ligand part are colored pale cyan. **E,F) a1d** bound to BRD4$^{BD1}$, FKBP12 is not shown for clarity. BRD4$^{BD1}$ residues contacting the FKBP12 ligand part are colored pale green and BRD4$^{BD1}$ residues contacting the linker are colored pale pink. **G,H)** Lig Plot analysis of intra-ligand and 'cross-over' contacts of **a1d** bound to FKBP12 and BRD4$^{BD1}$ in the two subunits.

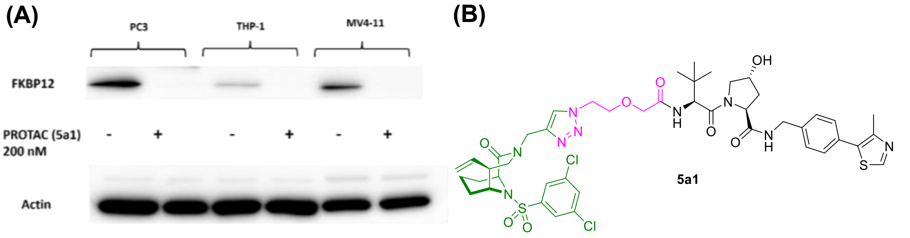

**SI Fig 7: Western Blot Analysis of FKBP12 levels and FKBP12 degradation in different cell lines.** **(A)** PC3 cells, serving as a positive control, were seeded in a 6-cm plate, while THP-1 and MV4-11 cell lines were seeded in T25 flasks for two days. Following this incubation, all cell lines were treated with 200 nM FKBP12-specific PROTAC for six hours to deplete FKBP12 protein levels. Western blot analysis was then performed to evaluate FKBP12 levels. **(B)** Chemical structure of the high-potency FKBP12 PROTAC 5a1.

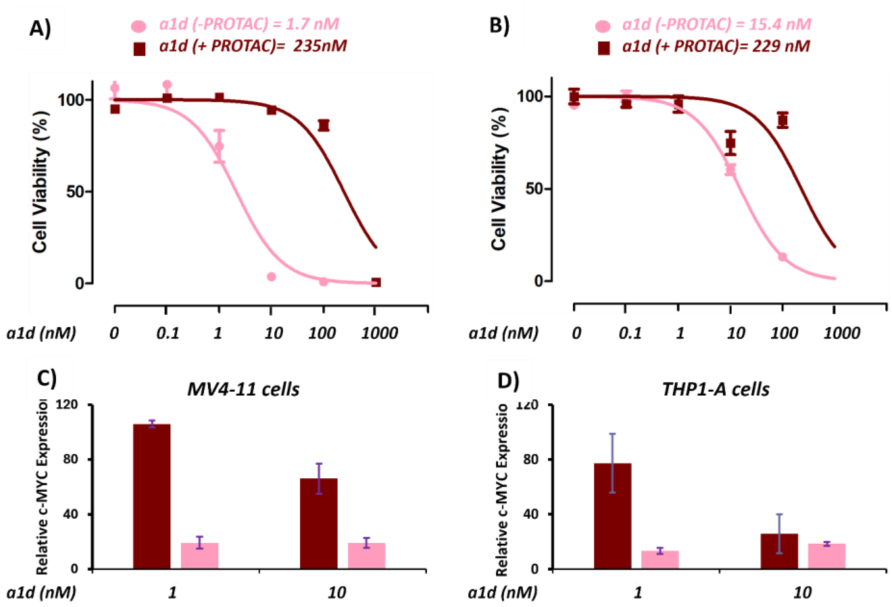

**SI Fig. 8: a1d cytotoxicity in MV4-11 and THP-1 cells.**

Cell toxicity for MV4-11 **(A)**, and THP-1 **(B)** treated for five days with the indicated concentrations of a1d in the presence or absence of PROTAC. Pink lines indicate absence of PROTAC, brown lines represent the PROTAC treatment. Estimated $IC_{50}$ values in the presence or absence of FKBP12 PROTAC are indicated above each panel. Quantitative analysis of MYC mRNA levels in MV4-11 cells **(C)**, and THP-1 **(D)** following treatment with a1d. All experiments were performed in triplicate. Error bars represent the mean ± standard deviation (SD) from three independent experiments.

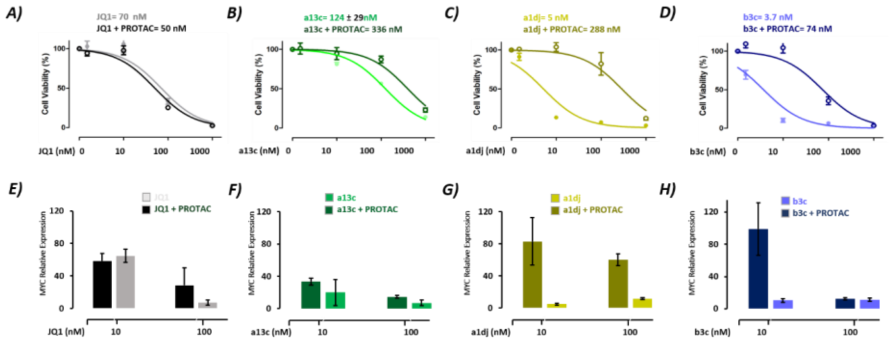

**SI Fig. 9: FKBP12-assisted toxicity by BRD4-targeting bifunctionals in THP-1 Cells.** (A-D) Cell toxicity for THP-1 cells treated for five days with the indicated concentrations of (A) JQ1, (B) a13c, (C) a1dj, and (D) b3c. All experiments were performed in triplicate, either in the presence or absence of FKBP12 PROTAC. Error bars represent the mean ± standard deviation (SD) from three independent experiments. Estimated $IC_{50}$ values in the presence or absence of FKBP12 PROTAC are indicated above each panel. (E-H) Quantitative analysis of MYC mRNA levels in THP-1 cells following treatment with (E) JQ1, (F) a13c, (G) a1dj, and (H) b3c for six hours. These experiments were conducted in the presence or absence of PROTAC. Error bars represent the mean ± standard deviation (SD) from three independent experiments.

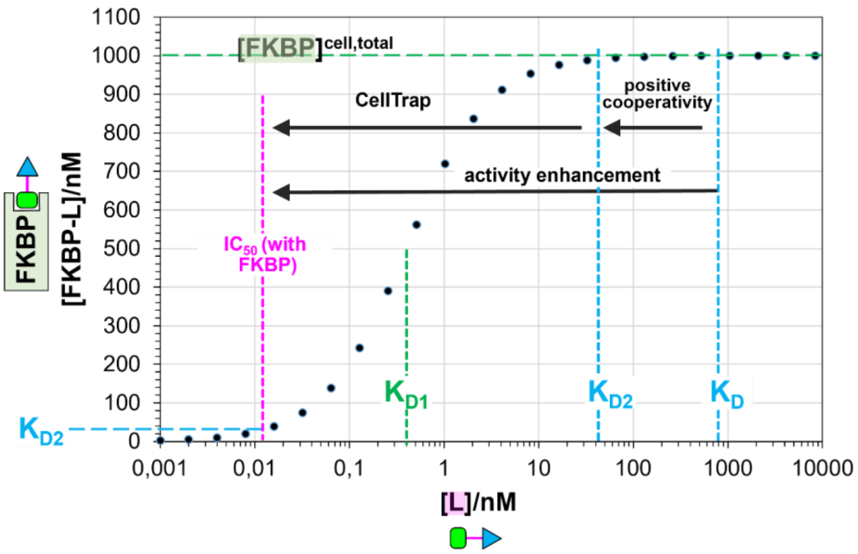

**SI Fig. 10 FKBP12-assisted activity enhancement exemplarily modelled for a1dj.**

The formation of the intracellular FKBP12-ligand complex in dependence of ligand concentration is modelled assuming a total intracellular FKBP12 concentration of 1 µM[1] and no ligand depletion (unlimited replenishment of free intracellular ligand by the extracellular ligand reservoir). Due to the high intracelluar FKBP12 concentration and the high $K_{D2}$ for ternary complex formation, only 1,4% of FKBP12 needs to the occupied to bind 50% of BRD4. Due to the high affinity to FKBP12 this is achieved at very low concentrations (12pM), representing an activity enhancement of more than 5 orders of magnitude. We note that under these extreme conditions the assumptions of ligand non-depletion in the extracellular medium might not be valid anymore.

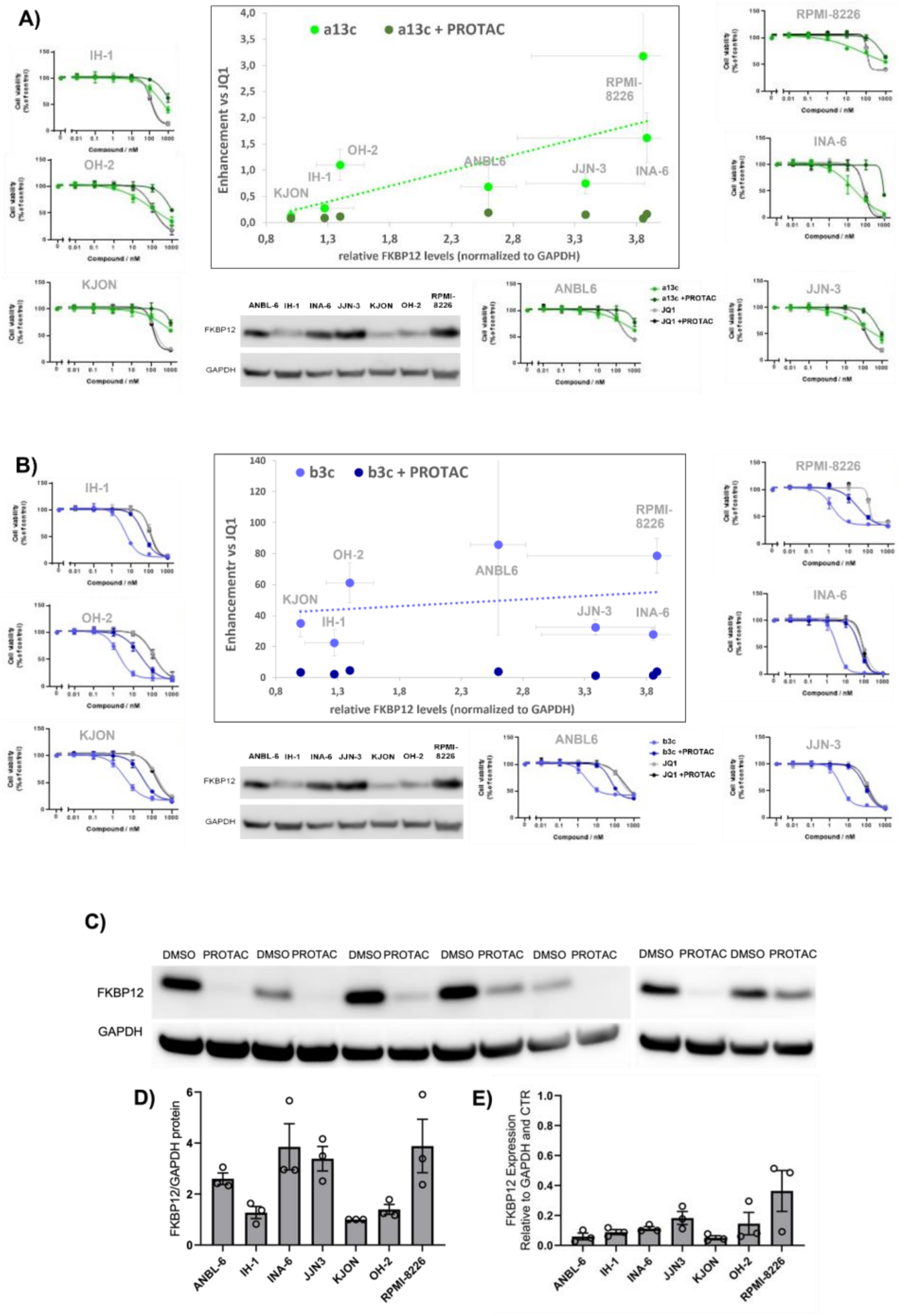

**SI Fig. 11: FKBP12-enhanced cell toxicity in myeloma cell lines. A)** Cell toxicity of a13c (green) correlates with FKBP12 expression levels and is substantially reduced after FKBP12 knockdown using a FKBP12 PROTAC. **B)** b3c (blue) is substantially enhanced compared to JQ1 in a panel of myeloma cell lines and this is abolished after FKBP12 knockdown using a FKBP12 PROTAC. C) Western Blot analysis of FKBP12 levels in the indicated myeloma cell lines in the presence or absence of FKB12 PROTAC. D) Quantification of FKBP12 levels normalized to GAPDH in the indicated myeloma cell lines. Circles represent measurements from three independent repeats, bars represent mean levels, error bar standard deviations. E) Quantification of FKBP12 depletion upon PROTAC treatment in in the indicated myeloma cell lines. Circles represent measurements from three independent repeats, bars represent mean levels, error bar standard deviations.

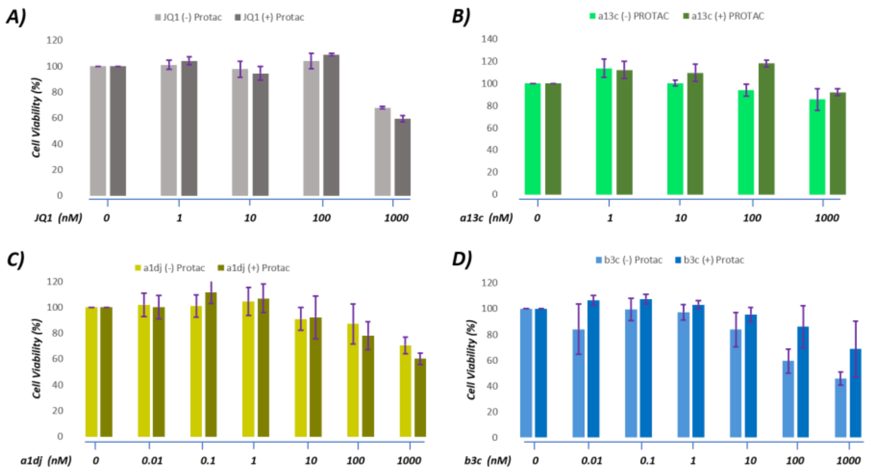

**SI Fig. 12: FKBP12-assisted toxicity by BRD4-targeting bifunctionals in PNT1-A cells.** (A-D) Cell toxicity for PNT1-A cells treated for five days with the indicated concentrations of (A) JQ1, (B) a13c, (C) a1dj, and (D) b3c. All experiments were performed in triplicate, either in the presence (lighter colors) or absence (darker color) of FKBP12 PROTAC. Error bars represent the mean ± standard deviation (SD) from three independent experiments.

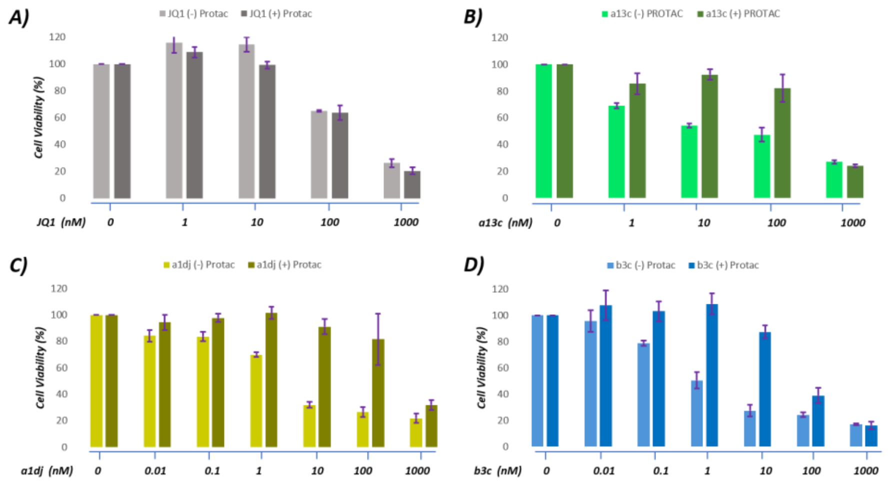

**SI Fig. 13: FKBP12-assisted toxicity by BRD4-targeting bifunctionals in PC3 cells.**

(A-D) Cell toxicity for PC3 cells treated for five days with the indicated concentrations of (A) JQ1, (B) a13c, (C) a1dj, and (D) b3c. All experiments were performed in triplicate, either in the presence (lighter colors) or absence (darker colors) of FKBP12 PROTAC. Error bars represent the mean ± standard deviation (SD) from three independent experiments.

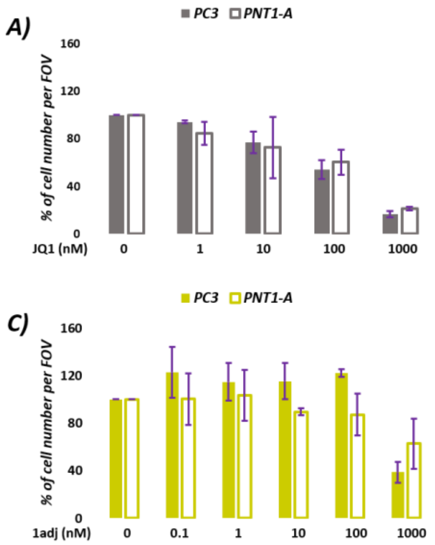

**SI Fig. 14: Cytotoxicity of co-cultured PC3 carcinoma and non-transformed PNT1A cells in the absence of FKBP12.** Microscopic cell count analysis in PNT1-A and PC3 cells following treatment for five days with various concentrations of (A) JQ1, (B) a13c, (C) a1dj, and (D) b3c. All cells were treated with the indicated compounds after FKBP12-degradation using PROTAC. Error bars represent the mean ± standard deviation (SD) from two independent live cell microscopy experiments.

### 3. Structure solution and refinement

The crystallographic experiments were performed at the BL14.1 at the Helmholtz-Zentrum BESSYII synchrotron, Berlin, Germany[2] and at the beamline ID23-1 of the European Synchrotron Radiation Facility (ESRF) in Grenoble, France and the raw data can be accessed at https://doi.org/10.15151/ESRF-DC-2127908021[3]. Diffraction data were integrated with XDS and further processed with the implemented programs of the CCP4i and CCP4i2 interface[4–8]. The data reduction was conducted with Aimless[7,9,10]. Crystal structures were solved by molecular replacement using Phaser[11]. Iterative model improvement and refinement were performed with Coot and Refmac5[12–17]. The dictionaries for the compounds were generated wit AceDRG implemented in CCP4i2[18]. Residues facing solvent channels without detectable side chain density were truncated.

| **PDB entry** | **9QW8** | **9R5N** |
|---|---|---|
| **Ligand name** | **a1d** | **b3c** |
| **Data collection** | | |
| Beamline | ESRF (ID23-1) | BESSY II (BL14.1) |
| Wavelength | $\lambda$ = 0.885603 Å | $\lambda$ = 0.9184 Å |
| Space group | P1 | $P2_12_12_1$ |
| Cell dimensions | | |
| *a*, *b*, *c* (Å) | 35.65, 35.65, 100.92 | 57.82, 90.51, 103.12 |
| $\alpha$, $\beta$, $\gamma$ (°) | 86.46, 84.22, 72.47 | 90, 90, 90 |
| Resolution (Å) | 100.35-1.80 (1.84-1.80) | 48.72-3.00 (3.18-3.00) |
| $R_{merge}$ | 0.094 (0.689) | 0.306 (0.660) |
| $R_{pim}$ | 0.061 (0.467) | 0.127 (0.269) |
| $I/\sigma(I)$ | 8.4 (1.7) | 8.9 (4.3) |
| CC1/2 | 0.977 (0.563) | 0.989 (0.936) |
| Completeness (%) | 97.0 (94.6) | 99.9 (100.0) |
| Redundancy | 3.3 (3.1) | 12.5 (13.2) |
| **Refinement** | | |
| Resolution (Å) | 100.35-1.80 | 48.77-3.00 |

**Commented [C1]:** Cite:
Gerlach, M.; Mueller, U.; Weiss, M. S., The MX beamlines BL14. 1-3 at BESSY II. *Journal of large-scale research facilities JLSRF* **2016,** *2*, 47

**Commented [AB2]:** Christian could you please check this, I could not find it

| | | |
|---|---|---|
| No. of reflections | 42396 | 11326 |
| $R_{work}/R_{free}$ (%) | 21.3/24.3 | 25.5/30.7 |
| No. of atoms | | |
| Protein | 2024/1979/1568/1563 | 1654/1543/1553/1553 |
| Ligand | 110/110 | 120/120 |
| Water | 256 | |
| *B*-factors | | |
| Protein | 20.1/21.0/27.7/35.0 | 32.2/31.3/23.7/23.3 |
| Ligand | 20.1/29.7 | 24.3/22.1 |
| Water | 27.2 | |
| R.m.s. deviations | | |
| Bond lengths (Å) | 0.0115 | 0.0194 |
| Bond angles (°) | 2.163 | 2.67 |
| Ramachandran plot | | |
| Favoured (%) | 97.0 | 93.0 |
| Allowed (%) | 3.0 | 6.0 |
| Outlier (%) | 0.0 | 1.0 |

### 4. General Synthetic Remarks

All reactions and manipulations which are sensitive towards air or moisture were performed under dry argon by using standard Schlenk techniques. All chemicals were purchased from Acros Organics, Sigma Aldrich, Alfa Aesar, TCI or ChemPUR. If not indicated otherwise, reagents and solvents were purchased from commercial suppliers and used without further treatment. All reactions were followed by TLC analysis or LC-MS. Flash silica gel column chromatography was performed with a Biotage® Isolera One system with Biotage® Sfär Silica HC D columns. Column chromatography was performed manually with silica gel 60 (0.04–0.063 mm) from Machery Nagel GmbH & Co. KG. Semi-preparative HPLC was performed with an Interchim PuriFlash 5250 system with a Luna® 5 µm C18(2) 100 Å, 250x21.2 mm column from Phenomenex. Eluents were 0.1 % TFA in water (Eluent A) and 0.1 % TFA in acetonitrile (Eluent B), methods are given in percentage B. All compounds were >95 % purity by HPLC if not noted otherwise. Preparative Separation by THAR SFC with Column Chiral Pak IE. Compound purity and low-resolution mass spectra were determined using an Agilent 1260 Infinity II system with a Poroshell 120 EC-C18 1.9 µm, 2.1 x 50 mm column from Agilent. Eluents were 0.1 % formic acid in water (Eluent A) and 0.1 % formic acid in acetonitrile (Eluent B), the used method was 5 % B to 100 % B in 2 min. MS was recorded with an Agilent InfinityLab G6125B LC/MSD. NMR spectroscopy was performed by the NMR department at TU Darmstadt. NMR spectra were recorded either on a 300 MHz Avance II NMR spectrometer from Bruker BioSpin GmbH (for $^{1}$H-NMR only), a 300 MHz Avance III NMR spectrometer from Bruker BioSpin GmbH (for $^{1}$H-, $^{13}$C-NMR), or a 500 MHz NMR spectrometer DRX 500 from Bruker BioSpin GmbH (for $^{1}$H- and $^{13}$C-NMR). NMR spectra were recorded at room temperature. The following abbreviations are used in the analysis of NMR spectra: s = singlet, d = doublet, t = triplet, q = quartet, hept = heptet, sept = septet, $s_{nr}$ = broad singlet. Combination of these abbreviations is applied whenever more than one coupling is observed. HRMS was performed by the mass spectrometry department at TU Darmstadt. Mass spectra were recorded on an Impact II, quadrupole-time-of-flight spectrometer from Bruker Daltonics. TLC was performed on TLC Silica gel 60 F254 Aluminum sheets from Merck Millipore. All final test compounds had a purity ≥95 % as determined by HPLC and UV detection at 220 nm.

### 5. Compound Synthesis

**General Procedure 1 (GP-1, HATU coupling)**

Under Argon atmosphere, the acid (1.0 eq.) and the amine (1.2 eq.) were dissolved in dry DCM [0.05M], then HATU (2.0 eq.) and DIPEA (2.5 eq.) were added, and the reaction mixture was stirred at room temperature. After 16 h, the solution was treated by brine and extracted by DCM. The organic phase was dried over anhydrous $Na_2SO_4$ and concentrated under reduced pressure.

**General Procedure 2 (GP-2,** Click reaction**)**

Azid (1.0 eq.) and alkyne (1.0 eq.) were dissolved in DMSO/$H_2O$ (v/v, 8/2), then aq. $CuSO_4$ (0.4 eq.) and sodium ascorbate (0.4 eq.) were added, and the reaction mixture was stirred at room temperature. The crude product was purified by prep. HPLC.

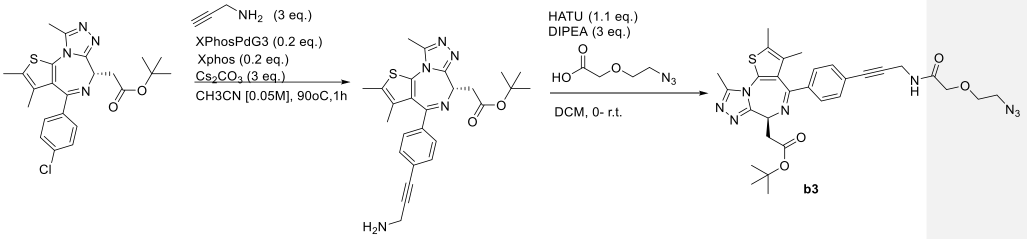

*tert*-butyl (*S*)-2-(4-(4-(3-aminoprop-1-yn-1-yl)phenyl)-2,3,9-trimethyl-6H-thieno[3,2-f][1,2,4]triazolo[4,3-a][1,4]diazepin-6-yl)acetate[19]

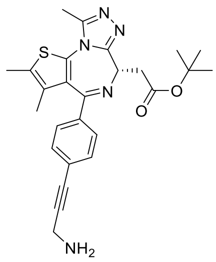

Under dry and argon atmosphere, JQ1 (228 mg, 0.5 mmol, 1.00 eq.), XPhos Pd G3 (85 mg, 0.1 mmol, 0.2 eq.), XPhos ligand (50 mg, 0.1 mmol, 0.2 eq.), propargylamine (83 mg, 1.5 mmol, 3.00 eq.) and $Cs_2CO_3$ (490 mg, 1.5 mmol, 3.00 eq.) were mixed in dry $CH_3CN$. The vial was capped and heated for 3h at 90 ºC to yield the title compound.

After completion of the reaction, the mixture was filtered through a pad of celite and purified by flash column chromatography on silica gel (0-100% MeOH in DCM). The obtained product was further purified by prep. HPLC (5-100% B), neutralized by $NaHCO_3$, extracted by *i*PrOH/$CHCl_3$ – 1/4, dried by lyophilization.

**Yield:** 28 mg (0.06 mmol, 12%).

**Physical State:** yellow solid.

**$R_f$ Value:** 0.38 (DCM/MeOH – 10/1).

**$^1$H NMR** (500 MHz, $CDCl_3$) δ 7.39 (d, *J* = 2.2 Hz, 4H), 4.55 (dd, *J* = 7.7, 6.4 Hz, 1H), 3.65 (s, 2H), 3.54 (dd, *J* = 7.0, 1.8 Hz, 2H), 2.65 (s, 3H), 2.39 (s, 3H), 1.66 (s, 3H), 1.49 (s, 9H) ppm.

**$^{13}$C NMR** (126 MHz, $CDCl_3$) δ 170.85, 164.11, 155.55, 149.79, 137.69, 132.20, 131.61, 130.94, 130.56, 130.54, 128.40, 125.42, 92.26, 82.06, 80.91, 53.94, 37.87, 32.19, 28.17, 14.33, 13.09, 11.87. ppm.

**HRMS** (ESI, m/z) calcd. for $C_{26}H_{30}N_5O_2S$: 476.21147, found: 476.21157.

*tert*-butyl (*S*)-2-(4-(4-(3-(2-(2-azidoethoxy)acetamido)prop-1-yn-1-yl)phenyl)-2,3,9-trimethyl-6H-thieno[3,2-f][1,2,4]triazolo[4,3-a][1,4]diazepin-6-yl)acetate **b3**

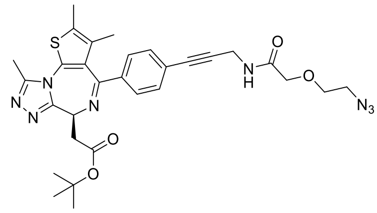

According to **GP-1**, the title compound **b3** was synthesized from *tert*-butyl (*S*)-2-(4-(4-(3-aminoprop-1-yn-1-yl)phenyl)-2,3,9-trimethyl-6H-thieno[3,2-f][1,2,4]triazolo[4,3-a][1,4]diazepin-6-yl)acetate[19](27 mg, 0.05 mmol, 1.00 eq.) and 2-(2-azidoethoxy)acetic acid (15 mg, 0.1 mmol, 2.00 eq.). The obtained product was further purified by prep. HPLC (5-100 % B).

**Yield:** 15 mg (0.025 mmol, 50%).

**Physical State:** yellow solid.

**$^1$H NMR** (500 MHz, $CDCl_3$)δ 7.41 (s, 4H), 6.90 (t, *J* = 5.3 Hz, 1H), 4.57 (dd, *J* = 7.9, 6.2 Hz, 1H), 4.33 (d, *J* = 5.4 Hz, 2H), 4.06 (s, 2H), 3.72 (dd, *J* = 5.4, 4.2 Hz, 2H), 3.55 – 3.52 (m, 2H), 3.50 – 3.45 (m, 2H), 2.70 (s, 3H), 2.41 (s, 3H), 1.66 (s, 3H), 1.49 (s, 9H).

**$^{13}$C NMR** (126 MHz, $CDCl_3$) δ 170.70, 168.80, 164.21, 155.43, 149.91, 137.86, 131.85, 131.72, 131.07, 131.05, 130.86, 128.42, 124.86, 86.52, 82.92, 81.08, 70.37, 70.34, 53.85, 50.66, 37.72, 29.49, 28.16, 14.34, 13.13, 11.67.

**HRMS** (ESI, m/z) calcd. for $C_{30}H_{35}N_8O_4S$ [M+H$^+$]: 603.24965, found: 603.24965.

*tert*-butyl 2-((6*S*)-4-(4-(3-(2-(2-(4-((((5*S*)-10-((3,5-dichlorophenyl)sulfonyl)-2-oxo-3-(pyridin-2-ylmethyl)-3,10-diazabicyclo[4.3.1]decan-5-yl)methoxy)methyl)-1H-1,2,3-triazol-1-yl)ethoxy)acetamido)prop-1-yn-1-yl)phenyl)-2,3,9-trimethyl-6H-thieno[3,2-f][1,2,4]triazolo[4,3-a][1,4]diazepin-6-yl)acetate **b3d**

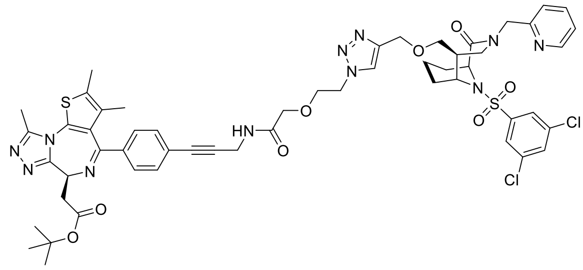

According to **GP-2**, the title compound **b3d** was synthesized synthesized from compound **b3** (6 mg, 10 µmol, 1.00 eq.) and (1S,5S,6R)-10-((3,5-dichlorophenyl)sulfonyl)-5-((prop-2-yn-1-yloxy)methyl)-3-(pyridin-2-ylmethyl)-3,10-diazabicyclo[4.3.1]decan-2-one[20] (5.2 mg, 10 µmol, 1.00 eq.). The obtained product was further purified by prep. HPLC (5-100 % B).

**Yield:** 4.5 mg (4 µmol, 40%).

**Purity:** 90 % (HPLC, UV-absorption 220 nm)

**Physical State:** yellow solid.

**HRMS** (ESI, m/z) calcd. for $C_{54}H_{61}Cl_2N_{11}O_8S_2$ [M+H$^+$]:1124.34393, found: 1124.34552.

*tert*-butyl 2-((6*S*)-4-(4-(3-(2-(2-(4-((5*S*)-10-((3,5-dichlorophenyl)sulfonyl)-2-oxo-3-(pyridin-

2-ylmethyl)-3,10-diazabicyclo[4.3.1]decan-5-yl)-1H-1,2,3-triazol-1-yl)ethoxy)acetamido)prop-1-yn-1-yl)phenyl)-2,3,9-trimethyl-6H-thieno[3,2-f][1,2,4]triazolo[4,3-a][1,4]diazepin-6-yl)acetate **b3c**

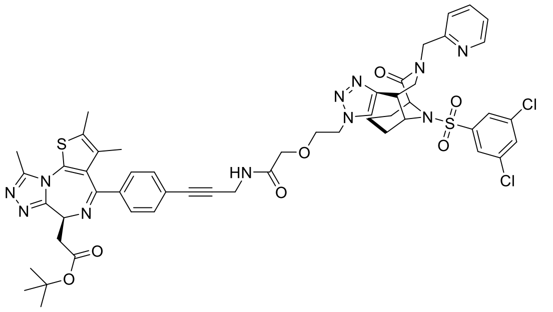

According to **GP-2**, the title compound **b3c** was synthesized from compound **b3** (6 mg, 10 µmol, 1.00 eq.) and (*1S,5S,6R*)-10-((3,5-dichlorophenyl)sulfonyl)-5-ethynyl-3-(pyridin-2-ylmethyl)-3,10-diazabicyclo[4.3.1]decan-2-one[20] (4.8 mg, 10 µmol, 1.00 eq.). The obtained product was further purified by prep. HPLC (5-100 % B).

**Yield:** 5.4 mg (4 µmol, 50%).

**Purity:** 99 % (HPLC, UV-absorption 220 nm)

**Physical State:** yellow solid.

**HRMS** (ESI, m/z) calcd. for $C_{52}H_{56}Cl_2N_{11}O_7S_2$ [M+H$^+$]:1080.31772, found: 1080.31935.

(*S*)-2-(4-(4-chlorophenyl)-2,3,9-trimethyl-6H-thieno[3,2-f][1,2,4]triazolo[4,3-a][1,4]diazepin-6-yl)acetic acid

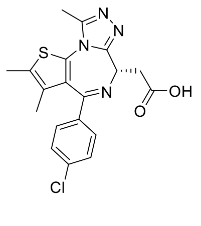

(+)-JQ1 (99 mg, 217 µmol, 1.0 eq.) was dissolved in formic acid / DCM (3 mL / 3 mL) and stirred at room temperature. After 2 hours, the solvent was evaporated *in vacuo* to afford the

title compound.

**Yield:** 86 mg (99 %)

**Purity:** 94 % (HPLC, UV-absorption 220 nm)

**Physical State:** yellow solid

**TLC:** $R_f$ = 0.11 (EA + 1 % FA)

**$^1$H-NMR** (500 MHz, $CDCl_3$) δ 7.42 (d, 2H, J = 8.5 Hz), 7.32 (d, 2H, J = 8.6 Hz), 4.60 (t, 1H, J = 6.9 Hz), 3.70 (dd, 1H, J = 16.9/6.8 Hz), 3.60 (dd, 1H, J = 16.9/7.2 Hz), 2.68 (s, 3H), 2.40 (s, 3H), 1.68 (s, 3H) ppm.

**$^{13}$C-NMR** (125 MHz, $CDCl_3$): δ 174.1, 164.2, 155.4, 150.1, 137.0, 136.5, 132.0, 131.3, 131.1, 130.7, 130.1, 128.8, 53.8, 36.9, 14.5, 13.2, 11.8 ppm.

(*S*)-N-((1-(2-(2-(2-(4-(4-chlorophenyl)-2,3,9-trimethyl-6H-thieno[3,2-f][1,2,4]triazolo[4,3-a][1,4]diazepin-6-yl)acetamido)ethoxy)ethyl)-1H-1,2,3-triazol-4-yl)methyl)-3',6'-dihydroxy-3-oxo-3H-spiro[isobenzofuran-1,9'-xanthene]-5-carboxamide

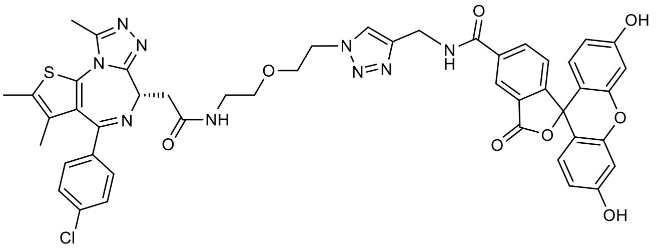

Compound **a1** (6.1 mg, 11.9 µmol, 1.0 eq.) and 3',6'-Dihydroxy-3-oxo-*N*-(prop-2-yn-1-yl)-3*H*-spiro[isobenzofuran-1,9'-xanthene]-5-carboxamide (5.0 mg, 12.1 µmol, 1.0 eq.) were dissolved in DMSO (1 mL), then $^t$BuOH (100 µL), $H_2O$ (100 µL), aq. $CuSO_4$ (1 M, 4.8 µL, 4.8 µmol, 0.4 eq.) and aq. sodium ascorbate (1 M, 4.8 µL, 4.8 µmol, 0.4 eq.) were added and the reaction mixture was stirred at room temperature. After 18 h, additional aq. $CuSO_4$ (1 M, 10 µL, 10 µmol, 0.8 eq.) and aq. sodium ascorbate (1 M, 10 µL, 10 µmol, 0.8 eq.) were added and the reaction mixture was stirred for another 4 h. Then brine was added and it was extracted with DCM, the organic phase was dried over $MgSO_4$, filtered and concentrated *in vacuo*. The crude product was purified by semi-preparative HPLC (20-70B, 9 CV) to afford the title compound.

**Yield:** 6.7 mg (61 %)

**Purity:** >99 % (HPLC, UV-absorption 220 nm)

**Physical State:** yellow solid

**HR-MS** (ESI): m/z calculated: $[M+H]^+$ = 926.24818, found: $[M+H]^+$ = 926.24934

(*S*)-*N*-(2-(2-azidoethoxy)ethyl)-2-(4-(4-chlorophenyl)-2,3,9-trimethyl-6*H*-thieno[3,2-*f*][1,2,4]triazolo[4,3-*a*][1,4]diazepin-6-yl)acetamide **a1**

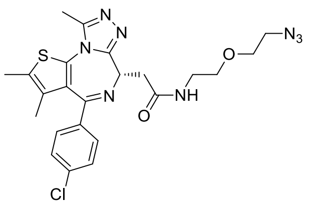

JQ1 acid (40 mg, 99.8 µmol, 1.0 eq.) and $H_2N$-PEG$_1$-$N_3$ (15.7 mg, 120 µmol, 1.2 eq.) were dissolved in dry DCM (2 mL), then HATU (77 mg, 202 µmol, 2.0 eq.) and DIPEA (42 µL, 247 µmol, 2.5 eq.) were added and the reaction mixture was stirred at room temperature. After 16 h, the solvent was evaporated *in vacuo* and the crude product was purified by semi-preparative HPLC (30-70B over 9 CV) to afford the title compound **a1**.

**Yield:** 54 mg (quant.)

**Purity:** 99 % (HPLC, UV-absorption 220 nm)

**Physical State:** yellow solid

**MS** (ESI): m/z calculated: $[M+H]^+$ = 513.2, found: $[M+H]^+$ = 513.2

**$^1$H-NMR** (500 MHz, $CDCl_3$): δ = 1.68 (s, 3H), 2.41 (s, 3H), 2.77 (s, 3H), 3.30-3.43 (m, 3H), 3.43-3.72 (m, 7H), 4.70 (t, 1H, J = 7.0 Hz), 7.02-7.10 (m, 1H), 7.34 (d, 2H, J = 8.6 Hz), 7.40 (d, 2H, J = 8.5 Hz) ppm.

**$^{13}$C-NMR** (125 MHz, $CDCl_3$): δ = 11.3, 13.2, 14.4, 38.2, 39.7, 50.7, 53.9, 69.5, 70.1, 128.9,

130.1, 130.9, 131.3, 131.5, 132.6, 135.8, 137.6, 150.4, 155.2, 164.9, 170.7 ppm.

(*S*)-*N*-(-(2-(2-(2-azidoethoxy)ethoxy)ethyl)-2-(4-(4-chlorophenyl)-2,3,9-trimethyl-6*H*-thieno[3,2-*f*][1,2,4]triazolo[4,3-*a*][1,4]diazepin-6-yl)acetamide **a2**

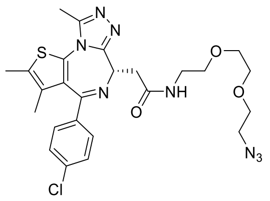

JQ1 acid (40 mg, 99.8 µmol, 1.0 eq.) and $H_2N$-$PEG_2$-$N_3$ (21.1 mg, 121 µmol, 1.2 eq.) were dissolved in dry DCM (2 mL), then HATU (77 mg, 202 µmol, 2.0 eq.) and DIPEA (42 µL, 247 µmol, 2.5 eq.) were added and the reaction mixture was stirred at room temperature. After 16 h, the solvent was evaporated *in vacuo* and the crude product was purified by semi-preparative HPLC (30-70B over 9 CV) to afford the title compound **a2**.

**Yield:** 43 mg (77 %)
**Purity:** >99 % (HPLC, UV-absorption 220 nm)
**Physical State:** yellow solid
**MS** (ESI): m/z calculated: $[M+H]^+$ = 557.2, found: $[M+H]^+$ = 557.2
**$^1$H-NMR** (500 MHz, $CDCl_3$): δ = 1.69 (s, 3H), 2.44 (s, 3H), 2.82 (s, 3H), 3.37-3.42 (m, 2H), 3.46 (m, 1H, J = 15.0/6.9 Hz), 3.49-3.73 (m, 11H), 4.78 (t, 1H, J = 7.0 Hz), 7.26-7.31 (m, 1H), 7.36 (d, 2H, J = 8.7 Hz), 7.41 (d, 2H, J = 8.5 Hz) ppm.
**$^{13}$C-NMR** (125 MHz, $CDCl_3$): δ = 11.2, 13.3, 14.4, 37.7, 40.0, 50.7, 53.6, 69.4, 70.1, 70.5, 70.6, 129.1, 130.3, 131.1, 131.2, 131.8, 133.3, 135.0, 138.2, 150.7, 154.9, 165.7, 170.9 ppm.

2-((*S*)-4-(4-chlorophenyl)-2,3,9-trimethyl-6*H*-thieno[3,2-*f*][1,2,4]triazolo[4,3-*a*][1,4]diazepin-6-yl)-*N*-(2-(2-(4-((((1*S*,5*S*,6*R*)-10-((3,5-dichlorophenyl)sulfonyl)-2-oxo-3-(pyridin-2-ylmethyl)-3,10-diazabicyclo[4.3.1]decan-5-yl)methoxy)methyl)-1*H*-1,2,3-triazol-

1-yl)ethoxy)ethyl)acetamide **a1d**

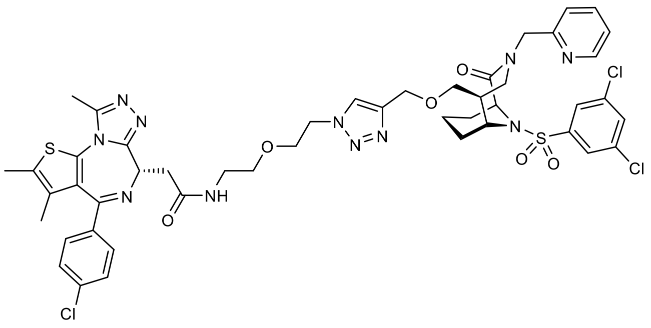

Compound **a1** (5.0 mg, 9.7 µmol, 1.0 eq.) and (*1S,5S,6R*)-10-((3,5-dichlorophenyl)sulfonyl)-5-((prop-2-yn-1-yloxy)methyl)-3-(pyridin-2-ylmethyl)-3,10-diazabicyclo[4.3.1]decan-2-one[20] (5.1 mg, 9.7 µmol, 1.0 eq.) were dissolved in DMSO (1 mL), then *ᵗ*BuOH (100 µL), $H_2O$ (100 µL), aq. $CuSO_4$ (1 M, 3.9 µL, 3.9 µmol, 0.4 eq.) and aq. sodium ascorbate (1 M, 3.9 µL, 3.9 µmol, 0.4 eq.) were added and the reaction mixture was stirred at room temperature. After 17 h, additional aq. $CuSO_4$ and sodium ascorbate (each 10 µL, 10 µmol, 1.0 eq.) were added and it was stirred for another 4 h. Brine was added and it was extracted with DCM, the organic phase was dried over $MgSO_4$, filtered and concentrated *in vacuo*. The crude product was purified by semi-preparative HPLC (30-70B, 9 CV) to afford the title compound **a1d**.

**Yield:** 7.8 mg (77 %)
**Purity:** >99 % (HPLC, UV-absorption 220 nm)
**Physical State:** yellowish solid
**HR-MS** (ESI): m/z calculated: $[M+H]^+$ = 1034.25253, found: $[M+H]^+$ = 1034.25307

2-((*S*)-4-(4-chlorophenyl)-2,3,9-trimethyl-6*H*-thieno[3,2-*f*][1,2,4]triazolo[4,3-*a*][1,4]diazepin-6-yl)-*N*-(2-(2-(4-((1*S*,5*S*,6*R*)-10-((3,5-dichlorophenyl)sulfonyl)-2-oxo-3-(pyridin-2-ylmethyl)-3,10-diazabicyclo[4.3.1]decan-5-yl)-1*H*-1,2,3-triazol-1-yl)ethoxy)ethyl)acetamide **a1c**

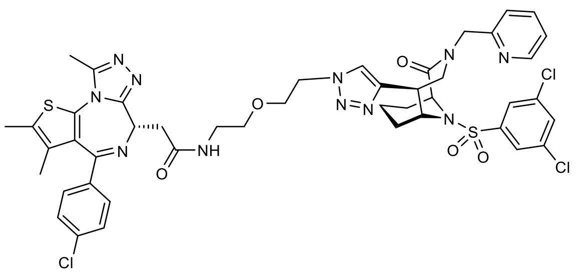

Compound **a1** (5.0 mg, 9.7 µmol, 1.0 eq.) and (*1S,5S,6R*)-10-((3,5-dichlorophenyl)sulfonyl)-5-ethynyl -3-(pyridin-2-ylmethyl)-3,10-diazabicyclo[4.3.1]decan-2-one [20] (4.8 mg, 9.9 µmol, 1.0 eq.) were dissolved in DMSO (1 mL), then $^t$BuOH (100 µL), $H_2O$ (100 µL), aq. $CuSO_4$ (1 M, 9.7 µL, 9.7 µmol, 1.0 eq.) and aq. sodium ascorbate (1 M, 9.7 µL, 9.7 µmol, 1.0 eq.) were added and the reaction mixture was stirred at room temperature. After 14 h, brine was added and it was extracted with DCM, the organic phase was dried over $MgSO_4$, filtered and concentrated *in vacuo*. The crude product was purified by semi-preparative HPLC (30-70B, 9 CV) to afford the title compound **a1c**.

**Yield:** 8.2 mg (85 %)

**Purity:** >99 % (HPLC, UV-absorption 220 nm)

**Physical State:** yellowish solid

**HR-MS** (ESI): m/z calculated: $[M+H]^+$ = 990.22632, found: $[M+H]^+$ = 990.22654

2-((*S*)-4-(4-chlorophenyl)-2,3,9-trimethyl-6*H*-thieno[3,2-*f*][1,2,4]triazolo[4,3-*a*][1,4]diazepin-6-yl)-*N*-(2-(2-(2-(4-((1*S*,5*S*,6*R*)-10-((3,5-dichlorophenyl)sulfonyl)-2-oxo-3-(pyridin-2-ylmethyl)-3,10-diazabicyclo[4.3.1]decan-5-yl)-1*H*-1,2,3-triazol-1-yl)ethoxy)ethoxy)ethyl)acetamide **a2c**

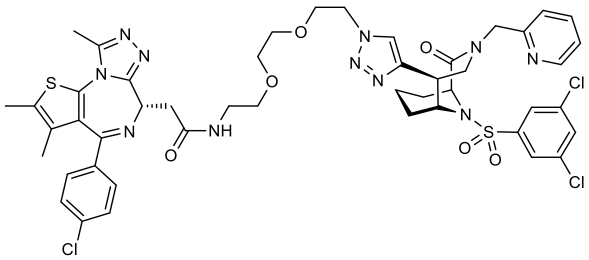

Compound **a2** (5.3 mg, 9.5 µmol, 1.0 eq.) and (*1S,5S,6R*)-10-((3,5-dichlorophenyl)sulfonyl)-5-ethynyl -3-(pyridin-2-ylmethyl)-3,10-diazabicyclo[4.3.1]decan-2-one [20] (4.4 mg, 9.2 µmol, 1.0 eq.) were dissolved in DMSO (1 mL), then $^{t}$BuOH (100 µL), $H_2O$ (100 µL), aq. $CuSO_4$ (1 M, 9.0 µL, 9.0 µmol, 1.0 eq.) and aq. sodium ascorbate (1 M, 9.0 µL, 9.0 µmol, 1.0 eq.) were added and the reaction mixture was stirred at room temperature. After 14 h, brine was added and it was extracted with DCM, the organic phase was dried over $MgSO_4$, filtered and concentrated *in vacuo*. The crude product was purified by semi-preparative HPLC (30-70B, 9 CV) to afford the title compound **a2c**.

**Yield:** 8.1 mg (85 %)

**Purity:** >99 % (HPLC, UV-absorption 220 nm)

**Physical State:** yellowish solid

**HR-MS** (ESI): m/z calculated: $[M+H]^+$ = 1034.25253, found: $[M+H]^+$ = 1034.25306

2-((*S*)-4-(4-chlorophenyl)-2,3,9-trimethyl-6*H*-thieno[3,2-*f*][1,2,4]triazolo[4,3-*a*][1,4]diazepin-6-yl)-*N*-(2-(2-(4-(((1*S*,5*S*,6*R*)-10-((3,5-dichlorophenyl)sulfonyl)-2-oxo-5-vinyl-3,10-diazabicyclo[4.3.1]decan-3-yl)methyl)-1*H*-1,2,3-triazol-1-yl)ethoxy)ethyl)acetamide **a1i**

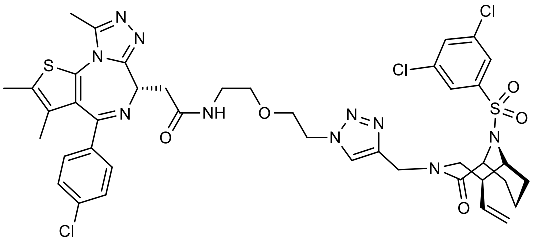

Compound **a1** (5.4 mg, 10.5 µmol, 1.1 eq.) and (1S,5S,6R)-10-((3,5-dichlorophenyl)sulfonyl)-3-(prop-2-yn-1-yl)-5-vinyl-3,10-diazabicyclo[4.3.1]decan-2-one [20] (4.3 mg, 9.9 µmol, 1.0 eq.) were dissolved in DMSO (1 mL), then $^t$BuOH (100 µL), $H_2O$ (100 µL), aq. $CuSO_4$ (1 M, 9.7 µL, 9.7 µmol, 1.0 eq.) and aq. sodium ascorbate (1 M, 9.7 µL, 9.7 µmol, 1.0 eq.) were added and the reaction mixture was stirred at room temperature. After 14 h, brine was added and it was extracted with DCM, the organic phase was dried over $MgSO_4$, filtered and concentrated *in vacuo*. The crude product was purified by semi-preparative HPLC (30-70B, 9 CV) to afford the title compound **a1i**.

**Yield:** 7.4 mg (78 %)
**Purity:** >99 % (HPLC, UV-absorption 220 nm)
**Physical State:** yellowish solid
**HR-MS** (ESI): m/z calculated: $[M+H]^+$ = 939.21542, found: $[M+H]^+$ = 939.21479

2-((*S*)-4-(4-chlorophenyl)-2,3,9-trimethyl-6*H*-thieno[3,2-*f*][1,2,4]triazolo[4,3-*a*][1,4]diazepin-6-yl)-*N*-(2-(2-(2-(4-(((1*S*,5*S*,6*R*)-10-((3,5-dichlorophenyl)sulfonyl)-2-oxo-5-vinyl-3,10-diazabicyclo[4.3.1]decan-3-yl)methyl)-1*H*-1,2,3-triazol-1-yl)ethoxy)ethoxy)ethyl)acetamide
**a2i**

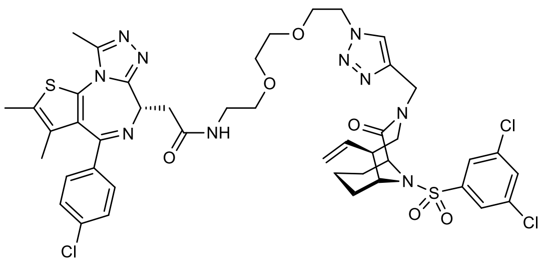

Compound **a2** (5.2 mg, 9.4 µmol, 1.0 eq.) and (*1S,5S,6R*)-10-((3,5-dichlorophenyl)sulfonyl)-3-(prop-2-yn-1-yl)-5-vinyl-3,10-diazabicyclo[4.3.1]decan-2-one [20] (3.8 mg, 9.0 µmol, 1.0 eq.) were dissolved in DMSO (1 mL), then $^{t}$BuOH (100 µL), $H_2O$ (100 µL), aq. $CuSO_4$ (1 M, 9.0 µL, 9.0 µmol, 1.0 eq.) and aq. sodium ascorbate (1 M, 9.0 µL, 9.0 µmol, 1.0 eq.) were added and the reaction mixture was stirred at room temperature. After 14 h, brine was added and it was extracted with DCM, the organic phase was dried over $MgSO_4$, filtered and concentrated *in vacuo*. The crude product was purified by semi-preparative HPLC (40-80B, 9 CV) to afford the title compound **a2i**.

**Yield:** 6.2 mg (70 %)

**Purity:** 95 % (HPLC, UV-absorption 220 nm)

**Physical State:** yellowish solid

**HR-MS** (ESI): m/z calculated: $[M+H]^+$ = 983.24163, found: $[M+H]^+$ = 953.24137

*N*-(2-(2-(4-(2-chloro-4-(((1*S*,5*S*,6*R*)-2-oxo-3-(pyridin-2-ylmethyl)-5-vinyl-3,10-diazabicyclo[4.3.1]decan-10-yl)sulfonyl)phenyl)-1*H*-1,2,3-triazol-1-yl)ethoxy)ethyl)-2-((*S*)-4-(4-chlorophenyl)-2,3,9-trimethyl-6*H*-thieno[3,2-*f*][1,2,4]triazolo[4,3-*a*][1,4]diazepin-6-yl)acetamide **a1h**

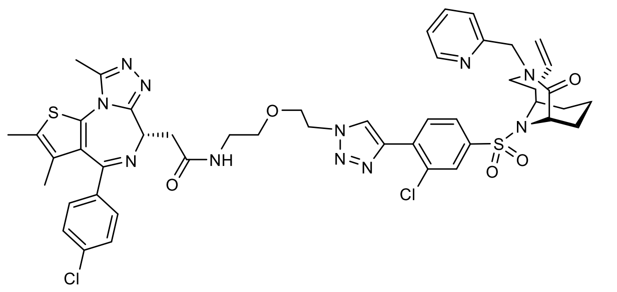

Compound **a1** (5.1 mg, 9.9 µmol, 1.1 eq.) and (*1S,5S,6R*)-10-((3-chloro-4-ethynylphenyl)sulfonyl)-3-(pyridin-2-ylmethyl)-5-vinyl-3,10-diazabicyclo[4.3.1]decan-2-one[20] (4.6 mg, 9.7 µmol, 1.0 eq.) were dissolved in DMSO (1 mL), then $^t$BuOH (100 µL), $H_2O$ (100 µL), aq. $CuSO_4$ (1 M, 9.7 µL, 9.7 µmol, 1.0 eq.) and aq. sodium ascorbate (1 M, 9.7 µL, 9.7 µmol, 1.0 eq.) were added and the reaction mixture was stirred at room temperature. After 14 h, brine was added and it was extracted with DCM, the organic phase was dried over $MgSO_4$, filtered and concentrated *in vacuo*. The crude product was purified by semi-preparative HPLC (30-70B, 9 CV) to afford the title compound **a1h**.

**Yield:** 3.6 mg (38 %)

**Purity:** 99 % (HPLC, UV-absorption 220 nm)

**Physical State:** yellowish solid

**HR-MS** (ESI): m/z calculated: $[M+H]^+$ = 982.28094, found: $[M+H]^+$ = 982.28242

*N*-(2-(2-(2-(4-(2-chloro-4-(((1*S*,5*S*,6*R*)-2-oxo-3-(pyridin-2-ylmethyl)-5-vinyl-3,10-diazabicyclo[4.3.1]decan-10-yl)sulfonyl)phenyl)-1*H*-1,2,3-triazol-1-yl)ethoxy)ethoxy)ethyl)-2-((*S*)-4-(4-chlorophenyl)-2,3,9-trimethyl-6*H*-thieno[3,2-*f*][1,2,4]triazolo[4,3-*a*][1,4]diazepin-6-yl)acetamide **a2h**

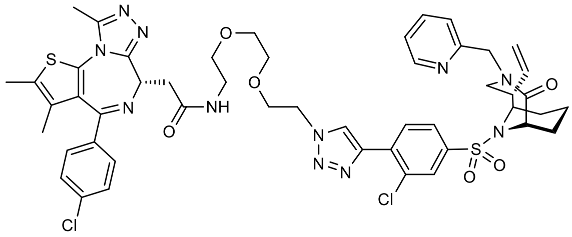

Compound **a2** (5.1 mg, 9.2 µmol, 1.0 eq.) and (*1S,5S,6R*)-10-((3-chloro-4-ethynylphenyl)sulfonyl)-3-(pyridin-2-ylmethyl)-5-vinyl-3,10-diazabicyclo[4.3.1]decan-2-one[20] (4.2 mg, 9.0 µmol, 1.0 eq.) were dissolved in DMSO (1 mL), then $^{t}$BuOH (100 µL), $H_2O$ (100 µL), aq. $CuSO_4$ (1 M, 9.0 µL, 9.0 µmol, 1.0 eq.) and aq. sodium ascorbate (1 M, 9.0 µL, 9.0 µmol, 1.0 eq.) were added and the reaction mixture was stirred at room temperature. After 14 h, brine was added and it was extracted with DCM, the organic phase was dried over $MgSO_4$, filtered and concentrated *in vacuo*. The crude product was purified by semi-preparative HPLC (30-70B, 9 CV) to afford the title compound **a2h**.

**Yield:** 3.8 mg (41 %)
**Purity:** >99 % (HPLC, UV-absorption 220 nm)
**Physical State:** yellowish solid
**HR-MS** (ESI): m/z calculated: $[M+H]^+$ = 1026.30715, found: $[M+H]^+$ = 1026.30723

*N*-(2-(2-(4-(3-chloro-5-(((1*S*,5*S*,6*R*)-2-oxo-3-(pyridin-2-ylmethyl)-5-vinyl-3,10-diazabicyclo[4.3.1]decan-10-yl)sulfonyl)phenyl)-1*H*-1,2,3-triazol-1-yl)ethoxy)ethyl)-2-((*S*)-4-(4-chlorophenyl)-2,3,9-trimethyl-6*H*-thieno[3,2-*f*][1,2,4]triazolo[4,3-*a*][1,4]diazepin-6-yl)acetamide **a1g**

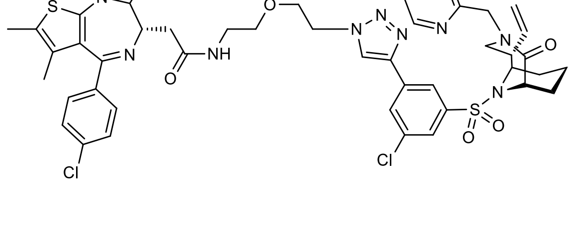

Compound **a1** (5.1 mg, 9.9 µmol, 1.1 eq.) and (*1S,5S,6R*)-10-((3-chloro-5-ethynylphenyl)sulfonyl)-3-(pyridin-2-ylmethyl)-5-vinyl-3,10-diazabicyclo[4.3.1]decan-2-one[20] (4.6 mg, 9.7 µmol, 1.0 eq.) were dissolved in DMSO (1 mL), then $^t$BuOH (100 µL), $H_2O$ (100 µL), aq. $CuSO_4$ (1 M, 9.7 µL, 9.7 µmol, 1.0 eq.) and aq. sodium ascorbate (1 M, 9.7 µL, 9.7 µmol, 1.0 eq.) were added and the reaction mixture was stirred at room temperature. After 14 h, brine was added and it was extracted with DCM, the organic phase was dried over $MgSO_4$, filtered and concentrated *in vacuo*. The crude product was purified by semi-preparative HPLC (35-75B, 9 CV) to afford the title compound **a1g**.

**Yield:** 7.3 mg (76 %)
**Purity:** 99 % (HPLC, UV-absorption 220 nm)
**Physical State:** yellowish solid
**HR-MS** (ESI): m/z calculated: $[M+H]^+$ = 982.28094, found: $[M+H]^+$ = 982.28149

*N*-(2-(2-(2-(4-(3-chloro-5-(((1*S*,5*S*,6*R*)-2-oxo-3-(pyridin-2-ylmethyl)-5-vinyl-3,10-diazabicyclo[4.3.1]decan-10-yl)sulfonyl)phenyl)-1*H*-1,2,3-triazol-1-yl)ethoxy)ethoxy)ethyl)-2-((*S*)-4-(4-chlorophenyl)-2,3,9-trimethyl-6*H*-thieno[3,2-*f*][1,2,4]triazolo[4,3-*a*][1,4]diazepin-6-yl)acetamide **a2g**

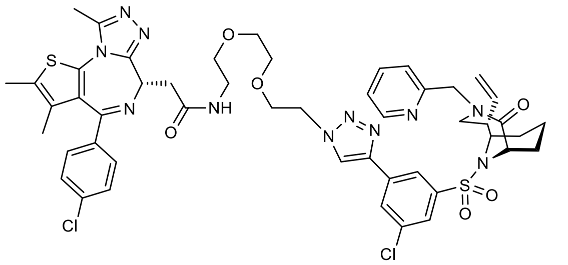

Compound **a2** (5.0 mg, 9.0 µmol, 1.0 eq.) and (*1S,5S,6R*)-10-((3-chloro-5-ethynylphenyl)sulfonyl)-3-(pyridin-2-ylmethyl)-5-vinyl-3,10-diazabicyclo[4.3.1]decan-2-one[20] (4.2 mg, 9.0 µmol, 1.0 eq.) were dissolved in DMSO (1 mL), then $^t$BuOH (100 µL), $H_2O$

(100 µL), aq. $CuSO_4$ (1 M, 9.0 µL, 9.0 µmol, 1.0 eq.) and aq. sodium ascorbate (1 M, 9.0 µL, 9.0 µmol, 1.0 eq.) were added and the reaction mixture was stirred at room temperature. After 14 h, brine was added and it was extracted with DCM, the organic phase was dried over $MgSO_4$, filtered and concentrated *in vacuo*. The crude product was purified by semi-preparative HPLC (33-73B, 9 CV) to afford the title compound **a2g**.

**Yield:** 6.1 mg (66 %)

**Purity:** >99 % (HPLC, UV-absorption 220 nm)

**Physical State:** yellowish solid

**HR-MS** (ESI): m/z calculated: $[M+H]^+$ = 1026.30715, found: $[M+H]^+$ = 1026.30718

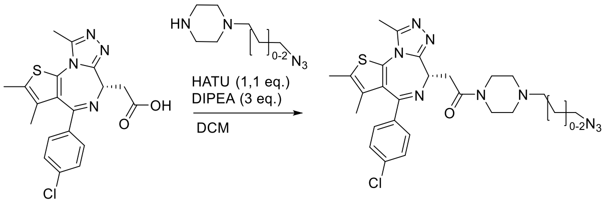

(*S*)-1-(4-(2-azidoethyl)piperazin-1-yl)-2-(4-(4-chlorophenyl)-2,3,9-trimethyl-6H-thieno[3,2-f][1,2,4]triazolo[4,3-a][1,4]diazepin-6-yl)ethan-1-on **a4**

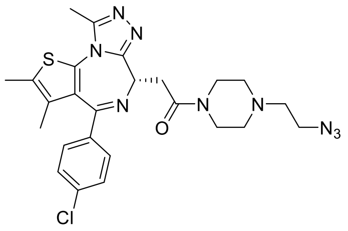

According to **GP-1**, the title compound **a4** was synthesized from JQ1 acid (30 mg, 75 µmol, 1.00 eq.) and 1-(2-azidoethyl)piperazine (19 mg, 100 µmol, 1.00 eq.). The obtained product was further purified by prep. HPLC (5-100 % B).

**Yield:** 19 mg (36 µmol, 48%).

**Physical State:** yellow solid.

**$^1$H NMR** (500 MHz, $CDCl_3$) δ 7.48 – 7.31 (m, 4H), 4.81 (dd, $J$ = 9.7, 5.2 Hz, 1H), 4.53 - 4.43 (m, 1H), 4.36 (d, $J$ = 14.9 Hz, 1H), 4.04 – 3.80 (m, 4H), 3.66 – 3.18 (m, 8H), 2.72 (s, 3H), 2.43 (s, 3H), 1.70 (s, 3H).

**$^{13}$C NMR** (126 MHz, $CDCl_3$) 168.9, 164.2, 155.4, 150.1, 137.0, 136.5, 132.0, 131.8, 131.1, 130.8, 129.8, 128.9, 55.5, 54.7, 52.1, 51.7, 45.9, 43.4, 39.4, 34.5, 14.5, 13.2, 11.5 ppm.

**HRMS** (ESI, m/z) calcd. for $C_{25}H_{29}ClN_9OS$ [M+H$^+$]:538.18988, found: 538.19029.

(*1S,5S,6R)*-5-(((1-(2-(4-(2-((S)-4-(4-chlorophenyl)-2,3,9-trimethyl-6H-thieno[3,2-f][1,2,4]triazolo[4,3-a][1,4]diazepin-6-yl)acetyl)piperazin-1-yl)ethyl)-1H-1,2,3-triazol-4-yl)oxy)methyl)-10-((3,5-dichlorophenyl)sulfonyl)-3-(pyridin-2-ylmethyl)-3,10-diazabicyclo[4.3.1]decan-2-one **a4d**

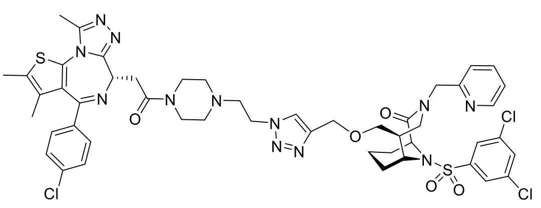

According to **GP-2**, the title compound **a4d** was synthesized from **a4** (5.4 mg, 10 µmol, 1.00 eq.) and (*1S,5S,6R*)-10-((3,5-dichlorophenyl)sulfonyl)-5-((prop-2-yn-1-yloxy)methyl)-3-(pyridin-2-ylmethyl)-3,10-diazabicyclo[4.3.1]decan-2-one[20] (5.2 mg, 10 µmol, 1.00 eq.). The obtained product was further purified by prep. HPLC (5-100 % B).

**Yield:** 5.5 mg (4 µmol, 50%).

**Purity:** 91 % (HPLC, UV-absorption 220 nm)

**Physical State:** yellow solid.

**HRMS** (ESI, m/z) calcd. for $C_{49}H_{54}Cl_3N_{12}O_5S_2$ [M+H$^+$]:1059.28416, found: 1059.28544.

(*S*)-1-(4-(3-azidopropyl)piperazin-1-yl)-2-(4-(4-chlorophenyl)-2,3,9-trimethyl-6H-thieno[3,2-f][1,2,4]triazolo[4,3-a][1,4]diazepin-6-yl)ethan-1-o **a5**

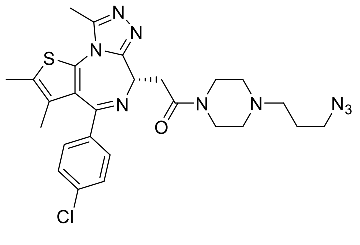

According to **GP-1**, the title compound **a5** was synthesized from JQ1 acid (40 mg, 100 µmol,

1.00 eq.) and 1-(3-azidopropyl)piperazine (19 mg, 100 µmol, 1.00 eq.). The obtained product was further purified by prep. HPLC (5-100 % B).

**Yield:** 6 mg (10 µmol, 10%).

**Physical State:** yellow solid.

**$^1$H NMR** (500 MHz, $CDCl_3$) δ 7.48 – 7.31 (m, 4H), 4.81 (dd, *J* = 9.7, 5.2 Hz, 1H), 4.53 - 4.43 (m, 1H), 4.55 – 4.41 (m, 1H), 4.04 – 3.80 (m, 4H), 3.66 – 3.18 (m, 10H), 2.72 (s, 3H), 2.43 (s, 3H), 1.70 (s, 3H).

**$^{13}$C NMR** (126 MHz, $CDCl_3$) 168.9, 164.2, 155.4, 150.1, 137.0, 136.5, 132.0, 131.8, 131.1, 130.8, 129.8, 128.9, 55.5, 54.7, 51.4, 51.6, 48.4, 43.4, 38.9, 34.5, 23.4, 14.5, 13.2, 11.5 ppm.

**HRMS** (ESI, m/z) calcd. for $C_{26}H_{31}ClN_9OS$ [M+Na$^+$]:574.18748, found: 574.18772.

(*1S,5S,6R*)-5-(((1-(3-(4-(2-((*S*)-4-(4-chlorophenyl)-2,3,9-trimethyl-6H-thieno[3,2-f][1,2,4]triazolo[4,3-a][1,4]diazepin-6-yl)acetyl)piperazin-1-yl)propyl)-1H-1,2,3-triazol-4-yl)oxy)methyl)-10-((3,5-dichlorophenyl)sulfonyl)-3-(pyridin-2-ylmethyl)-3,10-diazabicyclo[4.3.1]decan-2-one **a5d**

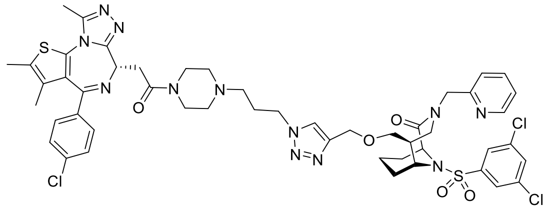

According to **GP-2**, the title compound **a5d** was synthesized from **a5** (5.4 mg, 10 µmol, 1.00 eq.) and (*1S,5S,6R*)-10-((3,5-dichlorophenyl)sulfonyl)-5-((prop-2-yn-1-yloxy)methyl)-3-(pyridin-2-ylmethyl)-3,10-diazabicyclo[4.3.1]decan-2-one[20] (5.2 mg, 10 µmol, 1.00 eq.). The obtained product was further purified by prep. HPLC (5-100 % B).

**Yield:** 5.7 mg (5.3 µmol, 53%).

**Purity:** 88 % (HPLC, UV-absorption 220 nm)

**Physical State:** yellow solid.

**HRMS** (ESI, m/z) $C_{50}H_{56}Cl_3N_{12}O_5S_2$ [M+H$^+$]:1073.29981, found: 1073.30181.

(*S*)-1-(4-(4-azidobutyl)piperazin-1-yl)-2-(4-(4-chlorophenyl)-2,3,9-trimethyl-6H-thieno[3,2-

f][1,2,4]triazolo[4,3-a][1,4]diazepin-6-yl)ethan-1-one **a6**

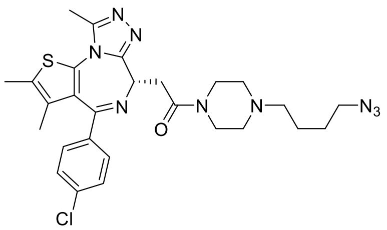

According to **GP-1**, the title compound **a6** was synthesized from JQ1 acid (20 mg, 50 µmol, 1.00 eq.) and 1-(4-azidobutyl)piperazine(18 mg, 100 µmol, 2.00 eq.). The obtained product was further purified by prep. HPLC (5-100 % B).

**Yield:** 12 mg (20 µmol, 43%).

**Physical State:** yellow solid.

**$^1$H NMR** (500 MHz, $CDCl_3$) δ 7.48 – 7.31 (m, 4H), 4.81 (dd, *J* = 9.7, 5.2 Hz, 1H), 4.53 - 4.43 (m, 1H), 4.55 – 4.41 (m, 1H), 4.04 – 3.80 (m, 4H), 3.66 – 3.18 (m, 12H), 2.72 (s, 3H), 2.43 (s, 3H), 1.70 (s, 3H).

**$^{13}$C NMR** (126 MHz, $CDCl_3$) 168.9, 164.2, 155.4, 150.1, 137.0, 136.5, 132.0, 131.8, 131.1, 130.8, 129.8, 128.9, 56.7, 51.5, 51.2, 50.5, 48.4, 43.4, 38.9, 34.5, 25.9, 21.1, 14.5, 13.2, 11.5 ppm.

**HRMS** (ESI, m/z) calcd. for $C_{27}H_{33}ClN_9O_{10}$ [M+H$^+$]:566.22118, found: 566.22120.

(*1S,5S,6R*)-5-(((1-(4-(4-(2-((*S*)-4-(4-chlorophenyl)-2,3,9-trimethyl-6H-thieno[3,2-f][1,2,4]triazolo[4,3-a][1,4]diazepin-6-yl)acetyl)piperazin-1-yl)butyl)-1H-1,2,3-triazol-4-yl)oxy)methyl)-10-((3,5-dichlorophenyl)sulfonyl)-3-(pyridin-2-ylmethyl)-3,10-diazabicyclo[4.3.1]decan-2-one **a6d**

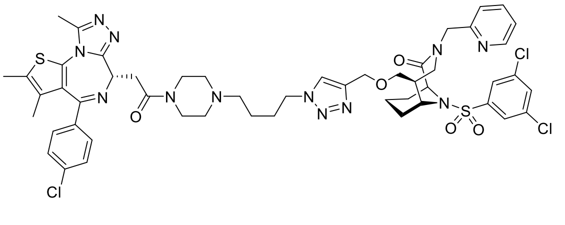

According to **GP-2**, the title compound **a6d** was synthesized from **a6** (5.7 mg, 10 µmol, 1.00 eq.) and (*1S,5S,6R*)-10-((3,5-dichlorophenyl)sulfonyl)-5-((prop-2-yn-1-yloxy)methyl)-3-(pyridin-

2-ylmethyl)-3,10-diazabicyclo[4.3.1]decan-2-one[20] (5.2 mg, 10 µmol, 1.00 eq.). The obtained product was further purified by prep. HPLC (5-100 % B).

**Yield:** 9 mg (8.3 µmol, 83%).

**Purity:** 99 % (HPLC, UV-absorption 220 nm)

**Physical State:** yellow solid.

**HRMS** (ESI, m/z) calcd. for $C_{51}H_{58}Cl_3N_{12}O_{10}S_2$ [M+H$^+$]:1087.31456, found: 1087.31535.

2-((*S*)-4-(4-chlorophenyl)-2,3,9-trimethyl-6H-thieno[3,2-f][1,2,4]triazolo[4,3-a][1,4]diazepin-6-yl)-N-(2-(2-(4-(((1S,5S,6R)-10-((3,5-dichlorophenyl)sulfonyl)-2-oxo-3-((S)-1-(pyridin-2-yl)ethyl)-3,10-diazabicyclo[4.3.1]decan-5-yl)methoxy)-1H-1,2,3-triazol-1-yl)ethoxy)ethyl)acetamide **a1dj**

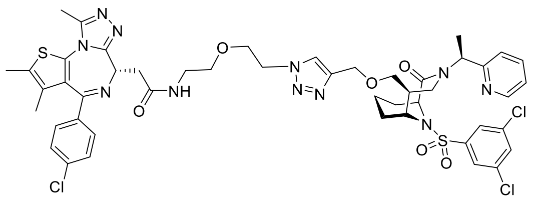

According to **GP-2**, the title compound **a1dj** was synthesized from compound **a1** (5.1 mg, 10 µmol, 1.00 eq.) and (*1S,5S,6R*)-10-((3,5-dichlorophenyl)sulfonyl)-5-((prop-2-yn-1-yloxy)methyl)-3-((*S*)-1-(pyridin-2-yl)ethyl)-3,10-diazabicyclo[4.3.1]decan-2-one [21] (5.2 mg, 10 µmol, 1.00 eq.). The obtained product was further purified by prep. HPLC (5-100 % B).

**Yield:** 6 mg (6 µmol, 60%).

**Purity:** 89 % (HPLC, UV-absorption 220 nm)

**Physical State:** yellow solid.

**HRMS** (ESI, m/z) calcd. for $C_{48}H_{53}Cl_3N_{11}O_6S_2$ [M+H$^+$]:1048.26818, found: 1048.26826.

2-((*S*)-4-(4-chlorophenyl)-2,3,9-trimethyl-6H-thieno[3,2-f][1,2,4]triazolo[4,3-a][1,4]diazepin-6-yl)-N-(2-(2-(4-(((*1S,5S,6R*)-10-((3,5-dichlorophenyl)sulfonyl)-3-(2-(1-methylpiperidin-4-yl)-2-oxoethyl)-2-oxo-3,10-diazabicyclo[4.3.1]decan-5-yl)methoxy)-1H-1,2,3-triazol-1-yl)ethoxy)ethyl)acetamide **a1dk**

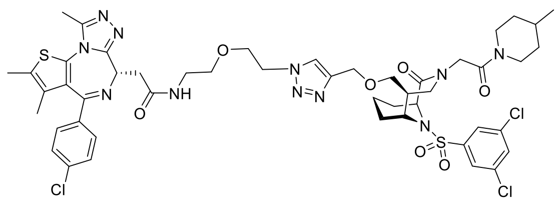

According to **GP-2**, the title compound **a1dk** was synthesized from compound **a1** (5.1 mg, 10 µmol, 1.00 eq.) and (*1S,5S,6R*)-10-((3,5-dichlorophenyl)sulfonyl)-3-(3-(4-methylpiperidin-1-

yl)-3-oxopropyl)-5-((prop-2-yn-1-yloxy)methyl)-3,10-diazabicyclo[4.3.1]decan-2-one [22] (5.7 mg, 10 µmol, 1.00 eq.). The obtained product was further purified by prep. HPLC (5-100 % B).

**Yield:** 6 mg (6 µmol, 60 %).

**Purity:** 96 % (HPLC, UV-absorption 220 nm)

**Physical State:** yellow solid.

**HRMS** (ESI, m/z) calcd. for $C_{49}H_{59}Cl_3N_{11}O_7S_2$ [M+H$^+$]:1082.31004, found: 1082.31015.

(5*S*)-10-((3,5-dichlorophenyl)sulfonyl)-5-((*R*)-1-hydroxyethyl)-3-(pyridin-2-ylmethyl)-3,10-diazabicyclo[4.3.1]decan-2-one[22]

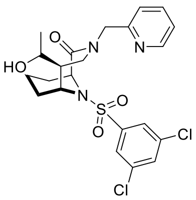

In a heatgun-dried 50 ml round flask 1.36 g of the (5*S*)-10-((3,5-dichlorophenyl)sulfonyl)-2-oxo-3-(pyridin-2-ylmethyl)-3,10-diazabicyclo[4.3.1]decane-5-carbaldehyde[22] was dissolved in 28 ml THF under argon atmosphere and cooled to 0°C. To the solution 0.91 ml (2.72 mmol, 1.2 eq) MeMgBr (3M solution in $Et_2O$) was added and the reaction mixture stirred for 17 hours at room temperature. The reaction was quenched with (100 ml) sat. $NH_4Cl$ solution. The aqueous phase was separated and extracted with DCM (3 x 60 ml). The combined organic layers were dried over $MgSO_4$ and the solvent removed under reduced pressure. The crude product was applied on (3 g) Silica and purified by column chromatography (50 g $SiO_2$, 40-100 % EA in Cy) to obtain the desired product.

**Yield:** 234 mg (6 µmol, 21 %).

**Purity:** 96 % (HPLC, UV-absorption 220 nm)

**Physical State:** off white foam.

**HRMS** (ESI, m/z) **:** calculated for $C_{22}H_{26}Cl_2N_3O_4S^+$ [M+H]$^+$: 498.1016, found: 484.1015.

**$^1$H-NMR** (500 MHz, $CDCl_3$,) $\delta$ 8.51 (dt, *J* = 5.8, 2.3 Hz, 1H), 7.74 – 7.65 (m, 3H), 7.56 (t, *J* = 1.9 Hz, 1H), 7.36 – 7.30 (m, 1H), 7.24 – 7.18 (m, 1H), 4.97 (dd, *J* = 15.0, 2.2 Hz, 1H), 4.76 (tq, *J* = 6.1, 1.9 Hz, 1H), 4.63 (dd, *J* = 15.0, 10.0 Hz, 1H), 4.20 – 4.11 (m, 1H), 3.94 (dd, *J* = 14.3, 10.6 Hz, 0.5H), 3.85 (dd, *J* = 14.6, 10.5 Hz, 0.5H), 3.75 (ddd, *J* = 10.8, 6.6, 4.9 Hz, 1H), 3.36 (ddd, *J* = 14.3, 11.3, 1.8 Hz, 1H), 2.34 – 2.25 (m, 1H), 1.58 – 1.39 (m, 5H), 1.29 – 1.21

(m, 2H), 1.16 (d, $J$ = 6.4 Hz, 1.5H), 1.12 (d, $J$ = 6.3 Hz, 1.5H) ppm.

**$^{13}$C-NMR** (125 MHz, $CDCl_3$) $\delta$ = 170.3, 170.4, 157.1, 157.1, 149.3, 149.2, 144.1, 144.1, 137.3, 137.2, 136.5, 132.9, 125.1, 125.1, 122.9, 122.8, 122.6, 66.8, 57.2, 57.2, 56.1, 52.7, 52.1, 50.8, 50.6, 48.9, 48.1, 28.6, 28.5, 28.2, 28.1, 21.0, 20.1, 15.6, 15.5 ppm.

(5*S*)-10-((3,5-dichlorophenyl)sulfonyl)-5-((*R*)-1-(prop-2-yn-1-yloxy)ethyl)-3-(pyridin-2-ylmethyl)-3,10-diazabicyclo[4.3.1]decan-2-one[22]

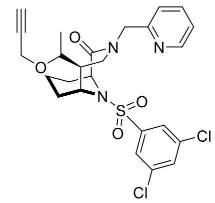

(5*S*)-10-((3,5-dichlorophenyl)sulfonyl)-5-((*R*)-1-hydroxyethyl)-3-(pyridin-2-ylmethyl)-3,10-diazabicyclo[4.3.1]decan-2-one (50 mg, 100 µmol, 1.0 eq.), was dissolved in DMF (dry, 5 mL) at 0 °C. Sodium hydride (13 mg, 300 µmol, 3.0 eq.) was added and the mixture was stirred for 30 min at 0 °C. 3-Bromoprop-1-yne (48 mg, 400 µmol, 4.0 eq.) and tetrabutylammonium iodide (1.8 mg, 5.0 µmol, 0.05 eq.) were added, the mixture was allowed to warm to room temperature and was stirred for 18 h. Saturated ammonium chloride solution (30 mL) was added and the aqueous solution was extracted with EA (3 x 30 mL). The combined organic phases were dried over MgSO4 and concentrated under reduced pressure. The obtained product was purified by chromatography (25 g SiO2, CH:EA = 1:1) and 2 times prep.HPLC (5-100 % B).

**Yield:** 25 mg (50 µmol, 25 %).

**Purity:** 95 % (HPLC, UV-absorption 220 nm)

**Physical State:** White solid.

**HRMS** (ESI, m/z) **:** calculated for $C_{25}H_{27}Cl_2N_3O_4S^+$ $[M+H]^+$: 536.1178, found: 536.1176.

The mixture was then separated by Chiral HPLC (Eluent A: $CO_2$ 70% Eluent B: Isopropanol 30%) and the separated 2 isomers were applied to click reaction.

2-((*S*)-4-(4-chlorophenyl)-2,3,9-trimethyl-6H-thieno[3,2-f][1,2,4]triazolo[4,3-a][1,4]diazepin-6-yl)-N-(2-(2-(4-(((*R*)-1-((1*S*,5*R*,6*R*)-10-((3,5-dichlorophenyl)sulfonyl)-2-oxo-3-(pyridin-2-ylmethyl)-3,10-diazabicyclo[4.3.1]decan-5-yl)ethoxy)methyl)-1H-1,2,3-triazol-1-yl)ethoxy)ethyl)acetamide **a1e** and **a1f**

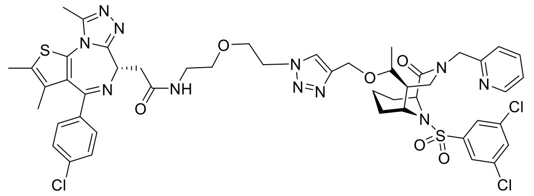

According to **GP-2**, the title compounds were synthesized from **a1** (5.7 mg, 10 µmol, 1.00 eq.) and the separated above isomers (5.2 mg, 10 µmol, 1.00 eq.) respectively. The corresponding obtained product was further purified by prep. HPLC (5-100 % B).

**Yield:** 6.6 mg (7 µmol, 70 %).

**Purity:** 91 % (HPLC, UV-absorption 220 nm)

**Physical State:** yellow solid.

**HRMS** (ESI, m/z) calcd. for $C_{48}H_{53}Cl_3N_{11}O_6S_2$ [M+H$^+$]:1048.26818, found: 1048.26923.

**Yield:** 3 mg (2.8 µmol, 28 %).

**Purity:** 70 % (HPLC, UV-absorption 220 nm)

**Physical State:** yellow solid.

**HRMS** (ESI, m/z) calcd. for $C_{48}H_{53}Cl_3N_{11}O_6S_2$ [M+H$^+$]:1048.26818, found: 1048.26923.

**2**-[(9*S*)-7-(4-chlorophenyl)-4,5,13-trimethyl-3-thia-1,8,11,12-tetraazatricyclo[8.3.0.0^{2,6}]trideca-2(6),4,7,10,12-pentaen-9-yl]-1-[4-(dimethoxymethyl)piperidin-1-yl]ethan-1-one

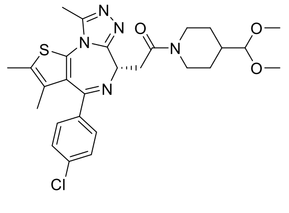

2-[(9S)-7-(4-chlorophenyl)-4,5,13-trimethyl-3-thia-1,8,11,12-

tetraazatricyclo[8.3.0.0²,6]trideca-2(6),4,7,10,12-pentaen-9-yl]acetic acid (500 mg; 1.13 mmol; 1.00 eq.) and HATU (517 mg; 1.36 mmol; 1.20 eq) were given to the vial and dissolved in N,N-Dimethylformamide (10 ml;), next N-ethyldiisopropylamine (491 µl; 2.83 mmol; 2.50 eq) was added. The mixture was stirred at RT for 15 minutes and 4-(dimethoxymethyl)piperidine (202 mg; 1.25 mmol; 1.10 eq.) added afterwards. The vial was sealed and stirred over night at room temperature

To the reaction mixture was added demineralized water and extraction carried out three times with EtOAc till there was no remaining product in water layer. The combined organic layers were washed with saturated NaCl-solution twice. Thereafter the organic layers were dried over sodium sulphate, filtered and all volatile ingredients were removed under reduced pressure. Next the crude product was purified with NP column chromatography, the pure fractions were combined and concentrated under reduced pressure to give 2-[(9S)-7-(4-chlorophenyl)-4,5,13-trimethyl-3-thia-1,8,11,12-tetraazatricyclo[8.3.0.0^{2,6}]trideca-2(6),4,7,10,12-pentaen-9-yl]-1-[4-(dimethoxymethyl)piperidin-1-yl]ethan-1-one (468 mg; 0.830 mmol; 73%).

Yield: 468 mg (0.830 mmol; 73%)

Purity: 96 % (HPLC, UV-absorption 254 nm)

Physical State: brown solid.

LC-MS (ESI, m/z) calcd. for C27H33ClN5O3S [M+H+]:542.2, found: 542.2.

1-{2-[(9*S*)-7-(4-chlorophenyl)-4,5,13-trimethyl-3-thia-1,8,11,12-tetraazatricyclo[8.3.0.0^{2,6}]trideca-2(6),4,7,10,12-pentaen-9-yl]acetyl}piperidine-4-carbaldehyde

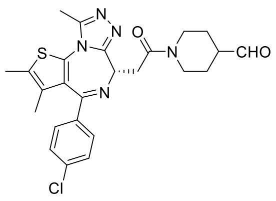

To a stirred solution of 2-[(9*S*)-7-(4-chlorophenyl)-4,5,13-trimethyl-3-thia-1,8,11,12-tetraazatricyclo[8.3.0.0^{2,6}]trideca-2(6),4,7,10,12-pentaen-9-yl]-1-[4-(dimethoxymethyl)piperidin-1-yl]ethan-1-one (200 mg; 0.353 mmol; 1.00 eq) in DCM

(2.50 ml) was added trifluoroacetic acid (1.00 ml) and water (100 µl) dropwise and the reaction mixture was stirred at room temperature for 2 h.

The reaction mixture was concentrated under reduced pressure to furnish 1-{2-[(9*S*)-7-(4-chlorophenyl)-4,5,13-trimethyl-3-thia-1,8,11,12-tetraazatricyclo[8.3.0.0^{2,6}]trideca-2(6),4,7,10,12-pentaen-9-yl]acetyl}piperidine-4-carbaldehyde (311 mg; 0.350 mmol; 100%) as crude product.

Yield: 311 mg (0.350 mmol; 100%)

Purity: 56 % (HPLC, UV-absorption 254 nm)

Physical State: yellow solid.

LC-MS (ESI, m/z) calcd. for $C_{25}H_{27}ClN_5O_2S$ [M+H+]:496.3, found: 496.1.

*tert*-butyl 4-azidopiperidine-1-carboxylate

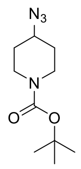

To a solution of 1-Boc-4-bromopiperidine (1.70 g; 6.43 mmol; 1.00 eq) in *N,N*-Dimethylformamide (17,0 ml) was added sodium azide (481 mg; 7,40 mmol; 1,15 eq) and the reaction mixture was stirred at 70°C and for 16 h.

The reaction mixture was poured into brine + demineralized water and extraction carried out three times with EtOAc. The organic layers were washed with brine twice, dried via sodium sulphate and concentrated under reduced pressure to give *tert*-butyl 4-azidopiperidine-1-carboxylate (1.37 g; 5.63 mmol; 88%).

Yield: 226 mg (1.28 mmol; 88%)

Purity: 93 % (HPLC, UV-absorption 220 nm)

Physical State: colourless oil

4-azidopiperidine hydrochloride

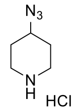

*tert*-butyl 4-azidopiperidine-1-carboxylate (310 mg; 1.28 mmol; 1.00 eq) was dissolved in D (600 µl) and 4M-HCl in Dioxane (1.59 mL; 6.38 mmol; 5.00 eq.) added dropwise to the stirred solution, the vial was purged with argon and sealed afterwards. The reaction mixture was stirred at RT for 2.5h.

The reaction mixture was evaporated under reduced pressure to give 4-azidopiperidine hydrochloride (226 mg; 1.28 mmol; 100%).

Yield: 1.37 g (5.63 mmol; 88%)

Purity: 93 % (HPLC, UV-absorption 220 nm)

Physical State: colourless oil

1-{4-[(4-azidopiperidin-1-yl)methyl]piperidin-1-yl}-2-[(9*S*)-7-(4-chlorophenyl)-4,5,13-trimethyl-3-thia-1,8,11,12-tetraazatricyclo[8.3.0.0^{2,6}]trideca-2(6),4,7,10,12-pentaen-9-yl]ethan-1-one **a7**

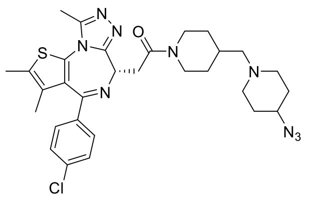

To a solution of 4-Azidopiperidine hydrochloride (60.0 mg; 0.339 mmol; 1.00 eq) and 1-{2-[(9*S*)-7-(4-chlorophenyl)-4,5,13-trimethyl-3-thia-1,8,11,12-tetraazatricyclo[8.3.0.0^{2,6}]trideca-2(6),4,7,10,12-pentaen-9-yl]acetyl}piperidine-4-carbaldehyde (311 mg; 0.353 mmol; 1.04 eq) in 1,2-DCE (1 mL) was added *N*-ethyldiisopropylamine (0.86 ml; 5.1 mmol; 15 eq) and the reaction mixture was stirred over night at RT. Sodium triacetoxyborohydride (222 mg; 1.02mmol; 3.00 eq.) was added and the reaction mixture stirred for 1.5h.

The reaction mixture was poured into demineralized water + $NaHCO_3$ and extraction carried out three times with EtOAc, till there was no remaining product in aq. layer. The combined organic layers were dried via phase separator tube concentrated under reduced pressure to give a crude material. The crude product was purified by NP column chromatography to furnish 1-{4-[(4-azidopiperidin-1-yl)methyl]piperidin-1-yl}-2-[(9S)-7-(4-chlorophenyl)-4,5,13-trimethyl-3-thia-1,8,11,12-tetraazatricyclo[8.3.0.0^{2,6}]trideca-2(6),4,7,10,12-pentaen-9-yl]ethan-1-one (91 mg; 0.15 mmol; 44%; colorless solid).

Yield: 91 mg (0.15 mmol; 45%)

Purity: > 99 % (HPLC, UV-absorption 254 nm)

Physical State: colourless solid.

LC-MS (ESI, m/z) calcd. for $C_{30}H_{37}ClN_9OS$ [M+H+]:606.3, found: 606.2.

$^1$H NMR (700 MHz, DMSO) δ 7.53 – 7.46 (m, 2H), 7.45 – 7.41 (m, 2H), 4.59 – 4.54 (m, 1H), 4.38 – 4.31 (m, 1H), 4.14 -4.09 (m, 1H), 3.63 – 3.51 (m 2H), 3.39 – 3.35 (m, 1H), 3.15 – 3.07 (m, 1H), 2.72 – 2.63 (s, 2H), 2.62 - 2.56 (m, 4H), 2.41 (s, 3H), 2.18 – 2.03 (m, 4H), 1.86 – 1.75 (m, 4H), 1.72 – 1.66 (m, 1H), 1.63 (s, 3H), 1.53 – 1.46 (m, 2H), 1.20 – 1.05 (m, 1H), 0.97 – 0.85 (m, 1H).

1-(4-{[4-(azidomethyl)piperidin-1-yl]methyl}piperidin-1-yl)-2-[(9*S*)-7-(4-chlorophenyl)-4,5,13-trimethyl-3-thia-1,8,11,12-tetraazatricyclo[8.3.0.0^{2,6}]trideca-2(6),4,7,10,12-pentaen-9-yl]ethan-1-one **a8**

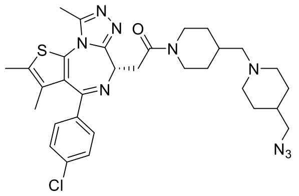

4(azidomethyl)piperidine hydrochloride (30.0 mg; 0.167 mmol; 1.00 eq), and 12-[(9*S*)-7-(4-chlorophenyl)-4,5,13-trimethyl-3-thia-1,8,11,12-tetraazatricyclo[8.3.0.0^{2,6}]trideca-2(6),4,7,10,12-pentaen-9-yl]acetyl}piperidine-4-carbaldehyde (162 mg; 0.183 mmol; 1.10 eq) and N-Ethyldiisopropylamine (0.43 ml; 2.5 mmol; 15 eq) were given into the vial, which was sealed afterwards. Next the vial was purged with argon and 1,2-dichloroethane (0.5 ml) was added. The reaction mixture was stirred at RT over night.

Next NaBH(OAc)3 (110 mg; 0.500 mmol; 3.00 eq) was added and the reaction mixture stirred for further 5 h.

Water (10 mL) and DCM (20 mL) were added to the reaction mixture, the phases were separated and the water phase was washed with DCM (2 x 20 mL). The combined organic layers were dried via phase separator tube concentrated under reduced pressure to give a crude material. The crude product was purified by NP column chromatography to furnish 1-(4-{[4-(azidomethyl)piperidin-1-yl]methyl}piperidin-1-yl)-2-[(9*S*)-7-(4-chlorophenyl)-4,5,13-trimethyl-3-thia-1,8,11,12-tetraazatricyclo[8.3.0.0^{2,6}]trideca-2(6),4,7,10,12-pentaen-9-yl]ethan-1-one (63 mg; 0.10 mmol; 56 %).

Yield: 63 mg (0.10 mmol, 56%).

Purity: 95 % (HPLC, UV-absorption 254 nm)

Physical State: white solid solid.

LC-MS (ESI, m/z) calcd. for $C_{31}H_{38}ClN_9OS$ [M+H+]:620.3, found: 620.3.

$^1$H NMR (700 MHz, DMSO) δ 7.51 – 7.46 (m, 2H), 7.46 – 7.41 (m, 2H), 4.60 – 4.55 (m, 1H), 4.38 – 4.32 (m, 1H), 4.14 – 4.09 (m, 1H), 3.60 (td, *J* = 16.9, 7.3 Hz, 1H), 3.36 (ddd, *J* = 16.4, 6.3, 2.2 Hz, 1H), 3.25 (d, *J* = 6.7 Hz, 2H), 3.15 – 3.08 (m, 1H), 2.84 (b, 2H), 2.63 – 2.54 (m, 4H), 2.42 (s, 3H), 2.13 (b, 2H), 1.91 – 1.75 (m, 2H), 1.71 – 1.61 (m, 5H), 1.53 – 1.47 (m, 1H), 1.30 – 1.06 (m, 3H), 0.95 – 0.85 (m, 1H).

(1S,5S,6R)-5-(((1-(1-((1-(2-((*S*)-4-(4-chlorophenyl)-2,3,9-trimethyl-6H-thieno[3,2-f][1,2,4]triazolo[4,3-a][1,4]diazepin-6-yl)acetyl)piperidin-4-yl)methyl)piperidin-4-yl)-1H-1,2,3-triazol-4-yl)methoxy)methyl)-10-((3,5-dichlorophenyl)sulfonyl)-3-(pyridin-2-ylmethyl)-3,10-diazabicyclo[4.3.1]decan-2-one **a7d**

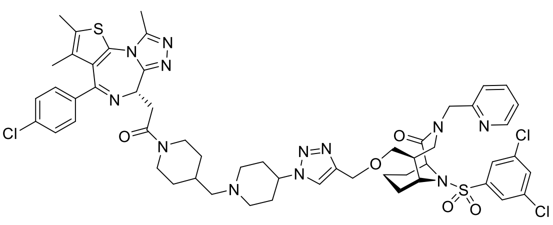

According to **GP-2**, the title compound **a7d** was synthesized from **a7** (6.1 mg, 10 µmol, 1.00 eq.) and (*1S,5S,6R*)-10-((3,5-dichlorophenyl)sulfonyl)-5-((prop-2-yn-1-yloxy)methyl)-3-(pyridin-2-ylmethyl)-3,10-diazabicyclo[4.3.1]decan-2-one [20] (5.2 mg, 10 µmol, 1.00 eq.). The obtained product was further purified by prep. HPLC (5-100 % B).

**Yield:** 10 mg (9 µmol, 90%).

**Purity:** 87 % (HPLC, UV-absorption 220 nm)

**Physical State:** yellow solid.

**HRMS** (ESI, m/z) calcd. for $C_{54}H_{62}Cl_3N_{12}O_5S_2$ [M+H$^+$]:1127.34676, found: 1127.34833.

(1S,5S,6R)-5-(1-(1-((1-(2-((*S*)-4-(4-chlorophenyl)-2,3,9-trimethyl-6H-thieno[3,2-f][1,2,4]triazolo[4,3-a][1,4]diazepin-6-yl)acetyl)piperidin-4-yl)methyl)piperidin-4-yl)-1H-1,2,3-triazol-4-yl)-10-((3,5-dichlorophenyl)sulfonyl)-3-(pyridin-2-ylmethyl)-3,10-diazabicyclo[4.3.1]decan-2-one **a7c**

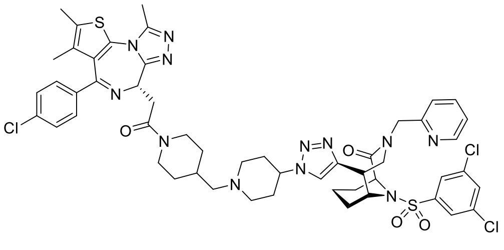

According to **GP-2**, the title compound **a7c** was synthesized from **a7** (6.1 mg, 10 µmol, 1.00 eq.) and (*1S,5S,6R*)-10-((3,5-dichlorophenyl)sulfonyl)- 5-ethynyl -3-(pyridin-2-ylmethyl)-3,10-diazabicyclo[4.3.1]decan-2-one [20] (5.2 mg, 10 µmol, 1.00 eq.). The obtained product was further purified by prep. HPLC (5-100 % B).

**Yield:** 9 mg (8.4 µmol, 84%).

**Purity:** 94 % (HPLC, UV-absorption 220 nm)

**Physical State:** yellow solid.

**HRMS** (ESI, m/z) calcd. for $C_{52}H_{58}Cl_3N_{12}O_4S_2$ [M+H$^+$]:1083.32055, found: 1083.32190.

(1S,5S,6R)-5-(((1-((1-((1-(2-((*S*)-4-(4-chlorophenyl)-2,3,9-trimethyl-6H-thieno[3,2-f][1,2,4]triazolo[4,3-a][1,4]diazepin-6-yl)acetyl)piperidin-4-yl)methyl)piperidin-4-yl)methyl)-1H-1,2,3-triazol-4-yl)methoxy)methyl)-10-((3,5-dichlorophenyl)sulfonyl)-3-(pyridin-2-ylmethyl)-3,10-diazabicyclo[4.3.1]decan-2-one **a8d**

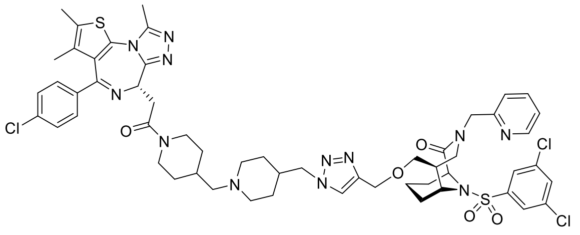

According to **GP-2**, the title compound **a8d** was synthesized from **a8** (6.2 mg, 10 µmol, 1.00 eq.) and (*1S,5S,6R*)-10-((3,5-dichlorophenyl)sulfonyl)-5-((prop-2-yn-1-yloxy)methyl)-3-(pyridin-2-ylmethyl)-3,10-diazabicyclo[4.3.1]decan-2-one [20] (5.2 mg, 10 µmol, 1.00 eq.). The obtained product was further purified by prep. HPLC (5-100 % B).

**Yield:** 6 mg (5.2 µmol, 52%).

**Purity:** 99 % (HPLC, UV-absorption 220 nm)

**Physical State:** yellow solid.

**HRMS** (ESI, m/z) calcd. for $C_{55}H_{64}Cl_3N_{12}O_5S_2$ [M+H$^+$]:1141.36241, found: 1141.36370.

(1S,5S,6R)-5-(1-((1-((1-(2-((*S*)-4-(4-chlorophenyl)-2,3,9-trimethyl-6H-thieno[3,2-f][1,2,4]triazolo[4,3-a][1,4]diazepin-6-yl)acetyl)piperidin-4-yl)methyl)piperidin-4-yl)methyl)-1H-1,2,3-triazol-4-yl)-10-((3,5-dichlorophenyl)sulfonyl)-3-(pyridin-2-ylmethyl)-3,10-diazabicyclo[4.3.1]decan-2-one **a8c**

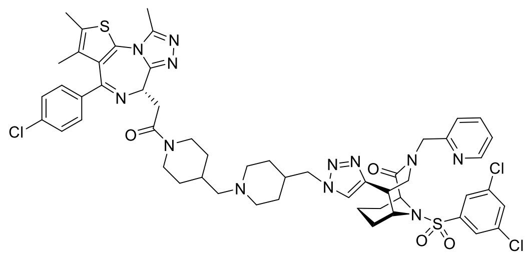

According to **GP-2**, the title compound **a8c** was synthesized from **a8** (6.2 mg, 10 µmol, 1.00 eq.) and (*1S,5S,6R*)-10-((3,5-dichlorophenyl)sulfonyl)- 5-ethynyl -3-(pyridin-2-ylmethyl)-3,10-diazabicyclo[4.3.1]decan-2-one [20] (5.2 mg, 10 µmol, 1.00 eq.). The obtained product was further purified by prep. HPLC (5-100 % B).

**Yield:** 8 mg (7.3 µmol, 73%).

**Purity:** 93 % (HPLC, UV-absorption 220 nm)

**Physical State:** yellow solid.

**HRMS** (ESI, m/z) calcd. for $C_{53}H_{60}Cl_3N_{12}O_4S_2$ [M+H$^+$]:1097.33620, found: 1097.33758.

*tert*-butyl *N*-(7-{2-[(9*S*)-7-(4-chlorophenyl)-4,5,13-trimethyl-3-thia-1,8,11,12-tetraazatricyclo[8.3.0.0^{2,6}]trideca-2(6),4,7,10,12-pentaen-9-yl]acetyl}-7-azaspiro[3.5]nonan-2-yl)carbamate

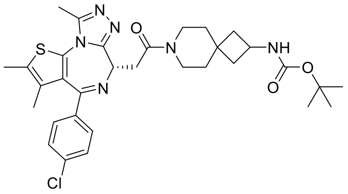

To a solution of 2-[(9*S*)-7-(4-chlorophenyl)-4,5,13-trimethyl-3-thia-1,8,11,12-tetraazatricyclo[8.3.0.0²,6]trideca-2(6),4,7,10,12-pentaen-9-yl]acetic acid (172 mg; 0.41 mmol; 0.86 eq.) and HATU (209 mg; 0.548 mmol; 1.15 eq.) in DMF (4 mL) was added *N*Ethyldiisopropylamine (207 µl; 1.19 mmol; 2.50 eq.). The mixture was stirred at RT for 15 minutes and *t*-butyl *N*-{7-azaspiro[3.5]nonan-2-yl}carbamate (133 mg; 0.53 mmol; 1.10 eq.) was added and it was stirred for 90 min at room temperature.

To the reaction mixture was added demineralized water and extraction carried out three times with DCM till there was no remaining product in water layer. Thereafter the organic layers were combined and dried via phase separator tube and concentrated in under reduced pressure. Next the crude product was purified with NP-column chromatography, the clean fractions were combined and concentrated under reduced pressure to furnish tert-butyl N-(7-{2-[(9S)-7-(4-chlorophenyl)-4,5,13-trimethyl-3-thia-1,8,11,12-tetraazatricyclo[8.3.0.0^{2,6}]trideca-2(6),4,7,10,12-pentaen-9-yl]acetyl}-7-azaspiro[3.5]nonan-2-yl)carbamate (335 mg; 0.43 mmol; 91%).

Yield: 355 mg (0.43 mmol, 91%).

Purity: 81 % (HPLC, UV-absorption 254 nm)

Physical State: pale yellow solid.

LC-MS (ESI, m/z) calcd. for $C_{32}H_{40}ClN_6O_3S$ [M+H+]:623.2, found: 623.2.

1-{2-amino-7-azaspiro[3.5]nonan-7-yl}-2-[(9*S*)-7-(4-chlorophenyl)-4,5,13-trimethyl-3-thia-1,8,11,12-tetraazatricyclo[8.3.0.0^{2,6}]trideca-2(6),4,7,10,12-pentaen-9-yl]ethan-1-one hydrochloride

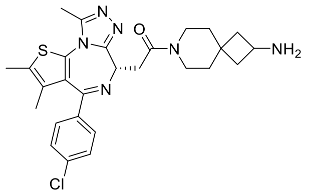

*tert*-butyl *N*-(7-{2-[(9*S*)-7-(4-chlorophenyl)-4,5,13-trimethyl-3-thia-1,8,11,12-tetraazatricyclo[8.3.0.0^{2,6}]trideca-2(6),4,7,10,12-pentaen-9-yl]acetyl}-7-azaspiro[3.5]nonan-2-yl)carbamate (335 mg; 0.433 mmol; 1.00 eq.) was dissolved in D (5 ml) and 4M-HCl in Dioxane (5 ml) added dropwise to the stirred solution. The reaction mixture was stirred at RT for 1.5h.

The reaction mixture was concentrated under reduced pressure to give a crude material, 1-{2-amino-7-azaspiro[3.5]nonan-7-yl}-2-[(9S)-7-(4-chlorophenyl)-4,5,13-trimethyl-3-thia-1,8,11,12-tetraazatricyclo[8.3.0.0^{2,6}]trideca-2(6),4,7,10,12-pentaen-9-yl]ethan-1-one hydrochloride (399 mg; 0.430 mmol; 100%).

Yield: 399 mg (0.43 mmol, 100%).

Purity: 61 % (HPLC, UV-absorption 254 nm)

Physical State: pale yellow solid.

LC-MS (ESI, m/z) calcd. for $C_{27}H_{32}ClN_6OS$ [M+H+]:523.2, found: 523.2.

1-{2-azido-7-azaspiro[3.5]nonan-7-yl}-2-[(9*S*)-7-(4-chlorophenyl)-4,5,13-trimethyl-3-thia-1,8,11,12-tetraazatricyclo[8.3.0.0^{2,6}]trideca-2(6),4,7,10,12-pentaen-9-yl]ethan-1-one **a13**

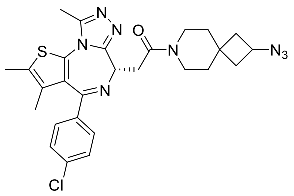

To a solution of 1-{2-amino-7-azaspiro[3.5]nonan-7-yl}-2-[(9*S*)-7-(4-chlorophenyl)-4,5,13-trimethyl-3-thia-1,8,11,12-tetraazatricyclo[8.3.0.0^{2,6}]trideca-2(6),4,7,10,12-pentaen-9-yl]ethan-1-one hydrochloride (399 mg; 0.433 mmol; 1.00 eq.) in DMSO & MTBE (1:1) (v/v) (9 ml) was added potassium hydroxide solution (aq.) (2.6 mmol; 6.00 eq.; 865.66 µl). *N*-diazosulfamoyl fluoride (5 eq) in MTBE (4 ml) and Dimethyl sulfoxide (4 ml) was added and the reaction mixture was stirred at RT for 1 h

The reaction mixture was acidified with 1M HCl till pH = 1 was met. Next the reaction mixture was poured into demineralized water and extraction carried out three times with EtOAc, till there was no remaining product in aq. layer. The combined organic layers were dried via phase separator tube and concentrated under reduced pressure to give a crude material. The crude product was fused onto Isolute H-MN and purified with RP-chromatography (see attached file), next the pure fractions were combined and concentrated. Next added demineralized water basified with a saturated sodium bicarbonate solution and extraction carried out three times with EtOAc till there was no remaining product in water layer. Thereafter the combined organic layers were concentrated under reduced pressure, resolved in MeCN/water, freezed in a dry-ice/acetone bath and lyophilized overnight in freeze dryer to give 1-{2-azido-7-

azaspiro[3.5]nonan-7-yl}-2-[(9S)-7-(4-chlorophenyl)-4,5,13-trimethyl-3-thia-1,8,11,12-tetraazatricyclo[8.3.0.0^{2,6}]trideca-2(6),4,7,10,12-pentaen-9-yl]ethan-1-one (172 mg; 0.310 mmol; 34%).

Yield: 172 mg (0.31 mmol, 34%).

Purity: 100 % (HPLC, UV-absorption 254 nm)

Physical State: pale yellow solid.

LC-MS (ESI, m/z) calcd. for $C_{27}H_{30}ClN_8OS$ [M+H+]:549.2, found: 549.2.

$^1$H NMR (500 MHz, DMSO) δ 7.51 – 7.47 (m, 2H), 7.45 – 7.41 (m, 2H), 4.58 – 4.54j (t, *J* = 6.7 Hz, 1H), 4.12 – 4.00 (m, 1H), 3.65 – 3.56 (m, 2H), 3.53 – 3.50 (m, 1H), 3.46 – 3.35 (m, 3H), 2.59 (s, 3H), 2.41 (s, 3H), 2.31 – 2.22 (m, 2H), 1.87 – 1.77 (m, 2H), 1.70 – 1.58 (m, 5H), 1.51 – 1.44 (m, 2H).

(*1S,5S,6R*)-5-(((1-(7-(2-((*S*)-4-(4-chlorophenyl)-2,3,9-trimethyl-6H-thieno[3,2-f][1,2,4]triazolo[4,3-a][1,4]diazepin-6-yl)acetyl)-7-azaspiro[3.5]nonan-2-yl)-1H-1,2,3-triazol-4-yl)methoxy)methyl)-10-((3,5-dichlorophenyl)sulfonyl)-3-(pyridin-2-ylmethyl)-3,10-diazabicyclo[4.3.1]decan-2-one **a13d**

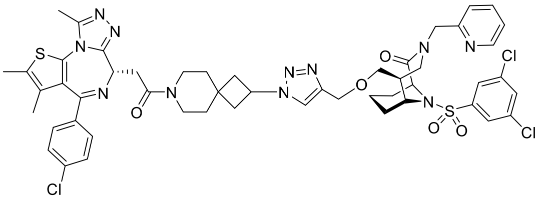

According to **GP-2**, the title compound **a13d** was synthesized from **a13** (5.5 mg, 10 µmol, 1.00 eq.) and (1S,5S,6R)-10-((3,5-dichlorophenyl)sulfonyl)-5-((prop-2-yn-1-yloxy)methyl)-3-(pyridin-2-ylmethyl)-3,10-diazabicyclo[4.3.1]decan-2-one [20] (5.2 mg, 10 µmol, 1.00 eq.). The obtained product was further purified by prep. HPLC (5-100 % B).

**Yield:** 6.4 mg (6.0 µmol, 60 %).

**Purity:** 88 % (HPLC, UV-absorption 220 nm)

**Physical State:** yellow solid.

**HRMS** (ESI, m/z) calcd. for $C_{51}H_{55}Cl_3N_{11}O_5S_2$ [M+H$^+$]:1070.28892, found: 1070.29133.

(*1S,5S,6R*)-5-(1-(7-(2-((*S*)-4-(4-chlorophenyl)-2,3,9-trimethyl-6H-thieno[3,2-f][1,2,4]triazolo[4,3-a][1,4]diazepin-6-yl)acetyl)-7-azaspiro[3.5]nonan-2-yl)-1H-1,2,3-triazol-4-yl)-10-((3,5-dichlorophenyl)sulfonyl)-3-(pyridin-2-ylmethyl)-3,10-diazabicyclo[4.3.1]decan-2-one **a13c**

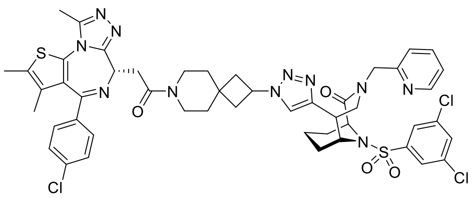

According to **GP-2**, the title compound **a13c** was synthesized from **a13** (8.3 mg, 15 µmol, 1.00 eq.) and (1S,5S,6R)-10-((3,5-dichlorophenyl)sulfonyl)- 5-ethynyl -3-(pyridin-2-ylmethyl)-3,10-diazabicyclo[4.3.1]decan-2-one [20] (7.2 mg, 15 µmol, 1.00 eq.). The obtained product was further purified by prep. HPLC (5-100 % B).

**Yield:** 11 mg (11 µmol, 73%).

**Purity:** 97 % (HPLC, UV-absorption 220 nm)

**Physical State:** yellow solid.

**HRMS** (ESI, m/z) calcd. for $C_{49}H_{51}Cl_3N_{11}O_4S_2$ [M+H$^+$]:1026.26270, found: 1026.26406.

1-{2-[(9*S*)-7-(4-chlorophenyl)-4,5,13-trimethyl-3-thia-1,8,11,12-tetraazatricyclo[8.3.0.0^{2,6}]trideca-2(6),4,7,10,12-pentaen-9-yl]acetyl}piperidin-4-one

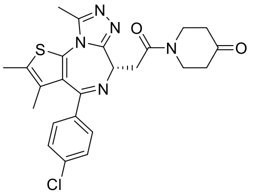

To a solution of 2-[(9S)-7-(4-chlorophenyl)-4,5,13-trimethyl-3-thia-1,8,11,12-tetraazatricyclo[8.3.0.0²,6]trideca-2(6),4,7,10,12-pentaen-9-yl]acetic acid (500 mg; 1.13 mmol; 1.00 eq.) and HATU (495 mg; 1.30 mmol; 1.15 eq.) in *N,N*-Dimethylformamide (10 ml) was added *N*-ethyldiisopropylamine (491 µl; 2.83 mmol; 2.50 eq.), the mixture was stirred at RT for 15 min and piperidin-4-one hydrochloride (169 mg; 1.25 mmol; 1.10 eq.) was added afterwards. The vial was sealed and stirred for 16 h at room temperature.

To the reaction mixture was added demineralized water and extraction carried out three times with EtOAc till there was no remaining product in water layer. Thereafter the organic layers were combined and dried via phase separator tube and concentrated under reduced pressure. Next the crude product was purified with NP-column chromatography, the clean fractions were combined and concentrated under reduced pressure to furnish 1-{2-[(9*S*)-7-(4-chlorophenyl)-4,5,13-trimethyl-3-thia-1,8,11,12-tetraazatricyclo[8.3.0.0^{2,6}]trideca-2(6),4,7,10,12-pentaen-9-yl]acetyl}piperidin-4-one (483 mg; 1.00 mmol; 89%).

Yield: 483 mg (1.00 mmol, 89%).

Purity: 100 % (HPLC, UV-absorption 254 nm)

Physical State: pale yellow solid.

LC-MS (ESI, m/z) calcd. for $C_{24}H_{25}ClN_5O_2S$ [M+H+] 482.1, found: 482.1.

1-[4-(azidomethyl)-[1,4'-bipiperidin]-1'-yl]-2-[(9*S*)-7-(4-chlorophenyl)-4,5,13-trimethyl-3-thia-1,8,11,12-tetraazatricyclo[8.3.0.0^{2,6}]trideca-2(6),4,7,10,12-pentaen-9-yl]ethan-1-one **a10**

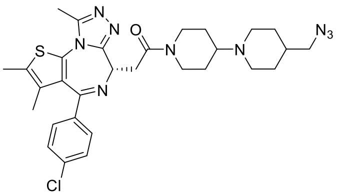

4-(azidomethyl)piperidine hydrochloride (60.0 mg; 0.329 mmol; 1.00 eq.), 1-{2-[(9*S*)-7-(4-chlorophenyl)-4,5,13-trimethyl-3-thia-1,8,11,12-tetraazatricyclo[8.3.0.0^{2,6}]trideca-2(6),4,7,10,12-pentaen-9-yl]acetyl}piperidin-4-one (175 mg; 0.362 mmol; 1.10 eq.) and *N*-Ethyldiisopropylamine (0.84 ml; 4.9 mmol; 15 eq.) were given into the vial, which was purged with argon and 1,2-dichloroethane (0.90 ml) was added. The reaction mixture was warmed up to 50°C and stirred at these temperature for 18h, therafter the reaction mixture was allowed to cool down to ambient temperature and stirred for the rest of weekend.

Next NaBH(OAc)$_3$ (218 mg; 0.987 mmol; 3.00 eq. -> repeated after several hours) was added and the reaction mixture stirred again for 48h at room temperature.

To the reaction mixture was added demineralized water and extraction carried out three times with EtOAc till there was no remaining product in water layer. Thereafter the organic layers were combined and dried via phase separator tube and concentrated under reduced pressure. Next the crude product was purified with RP-column chromatography, the clean fractions were combined and concentrated under reduced pressure to furnish 1-[4-(azidomethyl)-[1,4'-bipiperidin]-1'-yl]-2-[(9S)-7-(4-chlorophenyl)-4,5,13-trimethyl-3-thia-1,8,11,12-tetraazatricyclo[8.3.0.0^{2,6}]trideca-2(6),4,7,10,12-pentaen-9-yl]ethan-1-one (24 mg; 0.039 mmol; 12%).

Yield: 23.7 mg (0.04 mmol, 12%).

Purity: 99 % (HPLC, UV-absorption 254 nm)

Physical State: pale yellow solid.

LC-MS (ESI, m/z) calcd. for $C_{30}H_{37}ClN_9OS$ [M+H+]:606.3, found: 606.3.

$^1$H NMR (500 MHz, DMSO) δ 7.50 – 7.47 (m, 2H), 7.46 – 7.41 (m, 2H), 4.56 (td, *J* = 6.5, 1.5 Hz, 1H), 4.41 – 4.35 (m, *J* = 12.9 Hz, 1H), 4.19 – 4.13 (d, *J* = 13.6 Hz, 1H), 3.64 – 3.56 (m, 1H), 3.42 – 3.33 (m, 1H), 3.24 (dd, *J* = 6.7, 3.6 Hz, 2H), 3.14 – 3.06 (m, 1H), 2.89 – 2.83 (m, 1H), 2.59 (s, 3H), 2.42 (s, 3H), 2.16 – 2.08 (m, 2H), 1.84 – 1.68 (m, 2H), 1.67 – 1.61 (m, 5H), 1.57 – 1.43 (m, 2H), 1.30 – 1.11 (m, 4H).

1-{4-azido-[1,4'-bipiperidin]-1'-yl}-2-[(9*S*)-7-(4-chlorophenyl)-4,5,13-trimethyl-3-thia-1,8,11,12-tetraazatricyclo[8.3.0.0^{2,6}]trideca-2(6),4,7,10,12-pentaen-9-yl]ethan-1-one **a9**

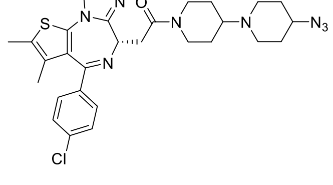

To a solution of 1-{2-[(9*S*)-7-(4-chlorophenyl)-4,5,13-trimethyl-3-thia-1,8,11,12-tetraazatricyclo[8.3.0.0^{2,6}]trideca-2(6),4,7,10,12-pentaen-9-yl]acetyl}piperidin-4-one

(200 mg; 415 µmol; 1.00 eq.) and *N*Ethyldiisopropylamine for synthesis (353 µl; 2.07 mmol; 5.00 eq.) in 1,2-Dichloroethane (5.00 ml) was added 4-azidopiperidine hydrochloride (142 mg; 830 µmol; 2.00 eq.) and the reaction mixture was stirred for 1 h at 80°C. Sodium triacetoxyborohydride (177 mg; 833 µmol; 3.00 eq.) was added and it was stirred for 1 h room temperature.

The reaction mixture was evapurated on celite and NP-column chromatography furnished 1-{4-azido-[1,4'-bipiperidin]-1'-yl}-2-[(9*S*)-7-(4-chlorophenyl)-4,5,13-trimethyl-3-thia-1,8,11,12-tetraazatricyclo[8.3.0.0^{2,6}]trideca-2(6),4,7,10,12-pentaen-9-yl]ethan-1-one (174 mg; 297 µmol) as a pale yellow foam.

Yield: 174 mg (0.294 mmol, 71%).

Purity: 100 % (HPLC, UV-absorption 254 nm)

Physical State: pale yellow foam.

LC-MS (ESI, m/z) calcd. for $C_{29}H_{35}ClN_9OS$ [M+H+]:592.2, found: 592.2.

$^1$H NMR (500 MHz, DMSO) δ 7.50 – 7.47 (m, 2H), 7.46 – 7.42 (m, 2H), 4.57 (t, *J* = 6.7 Hz, 1H), 3.86 (tt, *J* = 8.8, 3.9 Hz, 1H), 3.68 – 3.57 (m, 1H), 3.23 – 3.18 (m, 2H), 2.97 (ddd, *J* = 13.1, 9.8, 3.5 Hz, 2H), 2.60 (s, 3H), 2.42 (s, 3H), 2.02 (m, 2H), 1.71 – 1.64 (m, 2H), 1.63 (s, 3H), 1.30 – 1.22 (m, 6H).

(*1S,5S,6R*)-5-(1-((1'-(2-((*S*)-4-(4-chlorophenyl)-2,3,9-trimethyl-6H-thieno[3,2-f][1,2,4]triazolo[4,3-a][1,4]diazepin-6-yl)acetyl)-[1,4'-bipiperidin]-4-yl)methyl)-1H-1,2,3-triazol-4-yl)-10-((3,5-dichlorophenyl)sulfonyl)-3-(pyridin-2-ylmethyl)-3,10-diazabicyclo[4.3.1]decan-2-one **a10c**

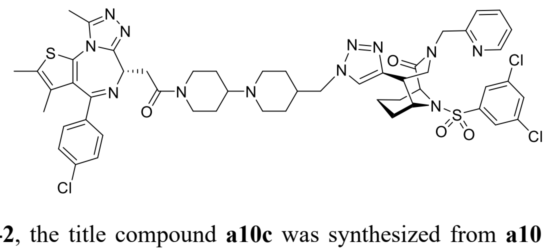

According to **GP-2**, the title compound **a10c** was synthesized from **a10** (6.3 mg, 10 µmol, 1.00 eq.) and (1S,5S,6R)-10-((3,5-dichlorophenyl)sulfonyl)- 5-ethynyl -3-(pyridin-2-ylmethyl)-3,10-diazabicyclo[4.3.1]decan-2-one [20] (4.8 mg, 10 µmol, 1.00 eq.). The obtained product was further purified by prep. HPLC (5-100 % B).

**Yield:** 9.2 mg (9.0 µmol, 90 %).

**Purity:** 98 % (HPLC, UV-absorption 220 nm)

**Physical State:** yellow solid.

**HRMS** (ESI, m/z) calcd. for $C_{52}H_{58}Cl_3N_{12}O_4S_2$ [M+H$^+$]:1083.32055, found: 1083.32267.

(*1S,5S,6R*)-5-(((1-((1'-(2-((*S*)-4-(4-chlorophenyl)-2,3,9-trimethyl-6H-thieno[3,2-f][1,2,4]triazolo[4,3-a][1,4]diazepin-6-yl)acetyl)-[1,4'-bipiperidin]-4-yl)methyl)-1H-1,2,3-triazol-4-yl)methoxy)methyl)-10-((3,5-dichlorophenyl)sulfonyl)-3-(pyridin-2-ylmethyl)-3,10-diazabicyclo[4.3.1]decan-2-one **a9d**

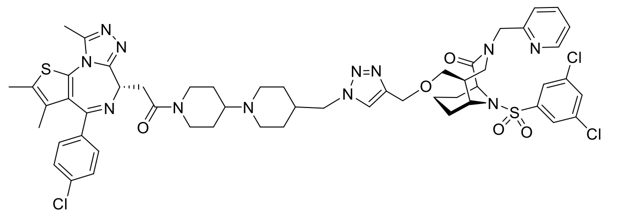

According to **GP-2**, the title compound **a9d** was synthesized from **a9** (6.1 mg, 10 µmol, 1.00 eq.) and (1S,5S,6R)-10-((3,5-dichlorophenyl)sulfonyl)-5-((prop-2-yn-1-yloxy)methyl)-3-(pyridin-2-ylmethyl)-3,10-diazabicyclo[4.3.1]decan-2-one [20] (5.2 mg, 10 µmol, 1.00 eq.). The obtained product was further purified by prep. HPLC (5-100 % B).

**Yield:** 8 mg (7.0 µmol, 71%).

**Purity:** 97 % (HPLC, UV-absorption 220 nm)

**Physical State:** yellow solid.

**HRMS** (ESI, m/z) calcd. for $C_{54}H_{62}Cl_3N_{12}O_5S_2$ [M+H$^+$]:1127.34676, found: 1127.34866.

(*1S,5S,6R*)-5-(1-(1'-(2-((*S*)-4-(4-chlorophenyl)-2,3,9-trimethyl-6H-thieno[3,2-f][1,2,4]triazolo[4,3-a][1,4]diazepin-6-yl)acetyl)-[1,4'-bipiperidin]-4-yl)-1H-1,2,3-triazol-4-yl)-10-((3,5-dichlorophenyl)sulfonyl)-3-(pyridin-2-ylmethyl)-3,10-diazabicyclo[4.3.1]decan-2-one **a9c**

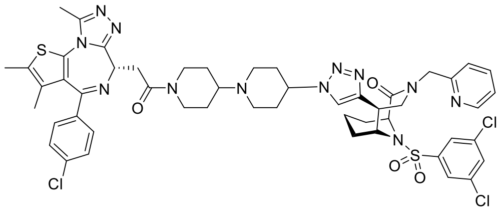

According to **GP-2**, the title compound **a9c** was synthesized from **a9** (5.9 mg, 10 µmol, 1.00 eq.) and (1S,5S,6R)-10-((3,5-dichlorophenyl)sulfonyl)- 5-ethynyl -3-(pyridin-2-ylmethyl)-3,10-diazabicyclo[4.3.1]decan-2-one [20] (4.8 mg, 10 µmol, 1.00 eq.). The obtained product was further purified by prep. HPLC (5-100 % B).

**Yield:** 6.7 mg (7.0 µmol, 70%).

**Purity:** 98 % (HPLC, UV-absorption 220 nm)

**Physical State:** yellow solid.

**HRMS** (ESI, m/z) calcd. for $C_{51}H_{56}Cl_3N_{12}O_4S_2$ [M+H$^+$]:1069.30490, found: 1069.30667.

(*1S,5S,6R*)-5-(((1-(1'-(2-((*S*)-4-(4-chlorophenyl)-2,3,9-trimethyl-6H-thieno[3,2-f][1,2,4]triazolo[4,3-a][1,4]diazepin-6-yl)acetyl)-[1,4'-bipiperidin]-4-yl)-1H-1,2,3-triazol-4-yl)methoxy)methyl)-10-((3,5-dichlorophenyl)sulfonyl)-3-(pyridin-2-ylmethyl)-3,10-diazabicyclo[4.3.1]decan-2-one **a9d**

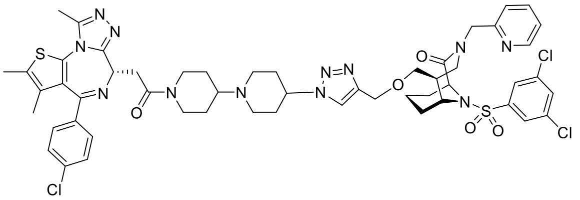

According to **GP-2**, the title compound **a9d** was synthesized from **a9** (12 mg, 20 µmol, 1.00 eq.) and (1S,5S,6R)-10-((3,5-dichlorophenyl)sulfonyl)-5-((prop-2-yn-1-yloxy)methyl)-3-(pyridin-2-ylmethyl)-3,10-diazabicyclo[4.3.1]decan-2-one [20] (11 mg, 20 µmol, 1.00 eq.). The obtained product was further purified by prep. HPLC (5-100 % B).

**Yield:** 11 mg (10 µmol, 50%).

**Purity:** 92 % (HPLC, UV-absorption 220 nm)

**Physical State:** yellow solid.

**HRMS** (ESI, m/z) calcd. for $C_{53}H_{60}Cl_3N_{12}O_5S_2$ [M+H$^+$]:1113.33111, found: 1113.33254.

2-[(9*S*)-7-(4-chlorophenyl)-4,5,13-trimethyl-3-thia-1,8,11,12-tetraazatricyclo[8.3.0.0^{2,6}]trideca-2(6),4,7,10,12-pentaen-9-yl]-1-[3-(hydroxymethyl)azetidin-1-yl]ethan-1-one

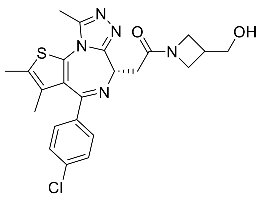

To a solution of 2-[(9*S*)-7-(4-chlorophenyl)-4,5,13-trimethyl-3-thia-1,8,11,12-tetraazatricyclo[8.3.0.0²,6]trideca-2(6),4,7,10,12-pentaen-9-yl]acetic acid (450 mg; 1.07 mmol; 1.00 eq.) and *N*-ethyldiisopropylamine (464 µl; 2.67 mmol; 2.50 eq.) in *N,N*-Dimethylformamide (5.00 ml) was added HATU (468 mg; 1.23 mmol; 1.15 eq.) and the mixture was stirred for 30 min at room temperature. azetidin-3-ylmethanol hydrochloride (153 mg; 1.18 mmol; 1.1 eq.) was added and it was stirred over night.

EtOAc (100 mL) and NaOH-solution (100 mL; 1 M aq.) were added, the phases were separated and the water phase was washed with EtOAc (2 x 40 mL). the combined organic phases were dried over $Na_2SO_4$ and all volatile ingredients were removed under reduced pressure to furnish crude product 2-[(9*S*)-7-(4-chlorophenyl)-4,5,13-trimethyl-3-thia-1,8,11,12-tetraazatricyclo[8.3.0.0^{2,6}]trideca-2(6),4,7,10,12-pentaen-9-yl]-1-[3-(hydroxymethyl)azetidin-1-yl]ethan-1-one (570 mg; 1.08 mmol) as a pale yellow solid.

Yield: 570 mg (1.80 mmol, 100%).

Purity: 89 % (HPLC, UV-absorption 254 nm)

Physical State: pale yellow solid.

LC-MS (ESI, m/z) calcd. for $C_{23}H_{25}ClN_5O_2S$ [M+H+]:470.1, found: 470.1.

1-{2-[(9*S*)-7-(4-chlorophenyl)-4,5,13-trimethyl-3-thia-1,8,11,12-tetraazatricyclo[8.3.0.0^{2,6}]trideca-2(6),4,7,10,12-pentaen-9-yl]acetyl}azetidine-3-carbaldehyde

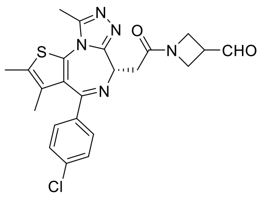

To a suspension of 2-[(9*S*)-7-(4-chlorophenyl)-4,5,13-trimethyl-3-thia-1,8,11,12-tetraazatricyclo[8.3.0.0^{2,6}]trideca-2(6),4,7,10,12-pentaen-9-yl]-1-[3-(hydroxymethyl)azetidin-1-yl]ethan-1-one (570 mg; 1.09 mmol; 1.00 eq.) and sodium hydrogen carbonate (367 mg; 4.37 mmol; 4.00 eq.) in D (5.00 ml) was added Dess-Martin periodinane (509 mg; 1.20 mol; 1.10 eq.) and the reaction mixture was stirred for 2 h at room temperature.

Water (10 ml) and $Na_2S_2O_3$-solution (aq) was added and the mixture was stirred for 30 min. The phases were separated and the water phase was washed with EtOAc (2 x 20 ml). The combined organic phases were dried over $Na_2SO_4$ and all volatile ingredients were removed under reduced pressure. NP-column chromatography furnished 1-{2-[(9*S*)-7-(4-chlorophenyl)-4,5,13-trimethyl-3-thia-1,8,11,12-tetraazatricyclo[8.3.0.0^{2,6}]trideca-2(6),4,7,10,12-pentaen-9-yl]acetyl}azetidine-3-carbaldehyde (268 mg; 573 mmol) as colourless oil.

Yield: 268 mg (0.573 mmol, 53%).

Purity: 100 % (HPLC, UV-absorption 254 nm)

Physical State: colourless oil.

LC-MS (ESI, m/z) calcd. for $C_{23}H_{23}ClN_5O_2S$ [M+H+]:468.1, found: 468.1.

1-{3-[(4-azidopiperidin-1-yl)methyl]azetidin-1-yl}-2-[(9*S*)-7-(4-chlorophenyl)-4,5,13-trimethyl-3-thia-1,8,11,12-tetraazatricyclo[8.3.0.0^{2,6}]trideca-2(6),4,7,10,12-pentaen-9-yl]ethan-1-one **a11**

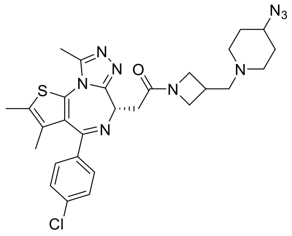

To a solution of 1-{2-[(9*S*)-7-(4-chlorophenyl)-4,5,13-trimethyl-3-thia-1,8,11,12-tetraazatricyclo[8.3.0.0^{2,6}]trideca-2(6),4,7,10,12-pentaen-9-yl]acetyl}azetidine-3-carbaldehyde (110 mg; 235 µmol; 1.00 eq.) and N-ethyldiisopropylamine (200 µl; 1.18 mmol; 5.00 eq.) in DCM (1 ml) was added 4azidopiperidine hydrochloride (80.4 mg; 470 µmol; 2.00 eq.) and the reaction mixture was stirred for 1 h at room temperature. Sodium triacetoxyborohydride (149 mg; 705 mmol; 3.00 eq.) was added and it was stirred 1 h.

The reaction mixture was evapurated on celite and NP-column chromatography furnished 1-{3-[(4-azidopiperidin-1-yl)methyl]azetidin-1-yl}-2-[(9S)-7-(4-chlorophenyl)-4,5,13-trimethyl-3-thia-1,8,11,12-tetraazatricyclo[8.3.0.0^{2,6}]trideca-2(6),4,7,10,12-pentaen-9-yl]ethan-1-one (150 mg; 252 µmol) as a colourless oil.

Yield: 150 mg (0.252 mmol, 107%).

Purity: 97 % (HPLC, UV-absorption 254 nm)

Physical State: colourless oil.

LC-MS (ESI, m/z) calcd. for $C_{28}H_{33}ClN_9OS$ [M+H+]:578.2, found: 578.3.

1-(3-{[4-(azidomethyl)piperidin-1-yl]methyl}azetidin-1-yl)-2-[(9*S*)-7-(4-chlorophenyl)-4,5,13-trimethyl-3-thia-1,8,11,12-tetraazatricyclo[8.3.0.0^{2,6}]trideca-2(6),4,7,10,12-pentaen-9-yl]ethan-1-one **a12**

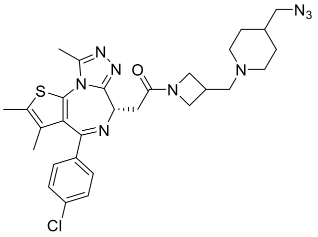

To a solution of 1-{2-[(9*S*)-7-(4-chlorophenyl)-4,5,13-trimethyl-3-thia-1,8,11,12-tetraazatricyclo[8.3.0.0^{2,6}]trideca-2(6),4,7,10,12-pentaen-9-yl]acetyl}azetidine-3-carbaldehyde (130 mg; 278 µmol; 1.00 eq.) and *N*-ethyldiisopropylamine (236 µl; 1.39 mmol; 5.00 eq.) in 1,2-dichloroethane (3.0 ml) and the reaction mixture was stirred for 1 h at room temperature. Sodium triacetoxyborohydride (177 mg; 833 µmol; 3.00 eq.) was added and it was stirred 1 h.

The reaction mixture was evapurated on celite and NP-column chromatography furnished 1-(3-{[4-(azidomethyl)piperidin-1-yl]methyl}azetidin-1-yl)-2-[(9S)-7-(4-chlorophenyl)-4,5,13-trimethyl-3-thia-1,8,11,12-tetraazatricyclo[8.3.0.0^{2,6}]trideca-2(6),4,7,10,12-pentaen-9-yl]ethan-1-one (160 mg; 270 µmol) as a colourless oil.

Yield: 160 mg (0.270 mmol, 97%).

Purity: 100 % (HPLC, UV-absorption 254 nm)

Physical State: colourless oil.

LC-MS (ESI, m/z) calcd. for $C_{29}H_{35}ClN_9OS$ [M+H+]:592.2, found: 592.2.

(*1S,5S,6R*)-5-(1-((1-((1-(2-((*S*)-4-(4-chlorophenyl)-2,3,9-trimethyl-6H-thieno[3,2-f][1,2,4]triazolo[4,3-a][1,4]diazepin-6-yl)acetyl)azetidin-3-yl)methyl)piperidin-4-yl)methyl)-1H-1,2,3-triazol-4-yl)-10-((3,5-dichlorophenyl)sulfonyl)-3-(pyridin-2-ylmethyl)-3,10-diazabicyclo[4.3.1]decan-2-one **a12c**

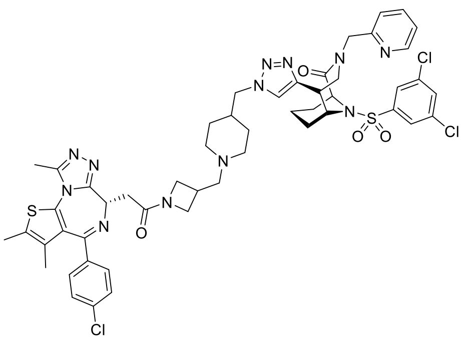

According to **GP-2**, the title compound **a12c** was synthesized from **a12** (6.8 mg, 12 µmol, 1.00 eq.) and (1S,5S,6R)-10-((3,5-dichlorophenyl)sulfonyl)- 5-ethynyl -3-(pyridin-2-ylmethyl)-3,10-diazabicyclo[4.3.1]decan-2-one [20] (5.0 mg, 12 µmol, 1.00 eq.). The obtained product was further purified by prep. HPLC (5-100 % B).

**Yield:** 12 mg (14 µmol, 95%).

**Purity:** 98 % (HPLC, UV-absorption 220 nm)

**Physical State:** yellow solid.

**HRMS** (ESI, m/z) calcd. for $C_{51}H_{56}Cl_3N_{12}O_4S_2$ [M+H$^+$]:1069.30490, found: 1069.30695.

(*1S,5S,6R*)-5-(((1-((1-((1-(2-((*S*)-4-(4-chlorophenyl)-2,3,9-trimethyl-6H-thieno[3,2-f][1,2,4]triazolo[4,3-a][1,4]diazepin-6-yl)acetyl)azetidin-3-yl)methyl)piperidin-4-yl)methyl)-1H-1,2,3-triazol-4-yl)methoxy)methyl)-10-((3,5-dichlorophenyl)sulfonyl)-3-(pyridin-2-ylmethyl)-3,10-diazabicyclo[4.3.1]decan-2-one **a12d**

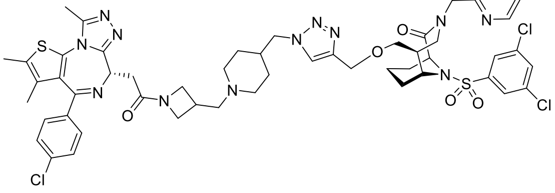

According to **GP-2**, the title compound **a12d** was synthesized from **a12** (7.1 mg, 12 µmol, 1.00 eq.) and (1S,5S,6R)-10-((3,5-dichlorophenyl)sulfonyl)-5-((prop-2-yn-1-yloxy)methyl)-3-(pyridin-2-ylmethyl)-3,10-diazabicyclo[4.3.1]decan-2-one [20] (6.2 mg, 12 µmol, 1.00 eq.). The obtained product was further purified by prep. HPLC (5-100 % B).

**Yield:** 11 mg (10 µmol, 84%).

**Purity:** 98 % (HPLC, UV-absorption 220 nm)

**Physical State:** yellow solid.

**HRMS** (ESI, m/z) calcd. for $C_{53}H_{60}Cl_3N_{12}O_5S_2$ [M+H$^+$]:1113.33111, found: 1113.33305.

(*1S,5S,6R*)-5-(1-(1-((1-(2-((*R*)-4-(4-chlorophenyl)-2,3,9-trimethyl-6H-thieno[3,2-f][1,2,4]triazolo[4,3-a]azepin-6-yl)acetyl)azetidin-3-yl)methyl)piperidin-4-yl)-1H-1,2,3-triazol-4-yl)-10-((3,5-dichlorophenyl)sulfonyl)-3-(pyridin-2-ylmethyl)-3,10-diazabicyclo[4.3.1]decan-2-one **a11c**

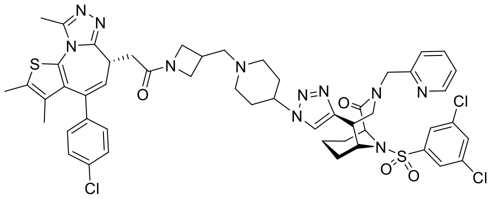

According to **GP-2**, the title compound **a11c** was synthesized from **a11** (6.3 mg, 10 µmol, 1.00 eq.) and (1S,5S,6R)-10-((3,5-dichlorophenyl)sulfonyl)- 5-ethynyl -3-(pyridin-2-ylmethyl)-3,10-diazabicyclo[4.3.1]decan-2-one [20] (5.0 mg, 10 µmol, 1.00 eq.). The obtained product was further purified by prep. HPLC (5-100 % B).

**Yield:** 7 mg (7.0 µmol, 70%).

**Purity:** 99 % (HPLC, UV-absorption 220 nm)

**Physical State:** yellow solid.

**HRMS** (ESI, m/z) calcd. for $C_{50}H_{54}Cl_3N_{12}O_4S_2$ [M+H$^+$]:1055.28925, found: 1055.29127.

(*1S,5S,6R*)-5-(((1-(1-((1-(2-((*S*)-4-(4-chlorophenyl)-2,3,9-trimethyl-6H-thieno[3,2-f][1,2,4]triazolo[4,3-a][1,4]diazepin-6-yl)acetyl)azetidin-3-yl)methyl)piperidin-4-yl)-1H-1,2,3-triazol-4-yl)methoxy)methyl)-10-((3,5-dichlorophenyl)sulfonyl)-3-(pyridin-2-ylmethyl)-3,10-diazabicyclo[4.3.1]decan-2-one **a11d**

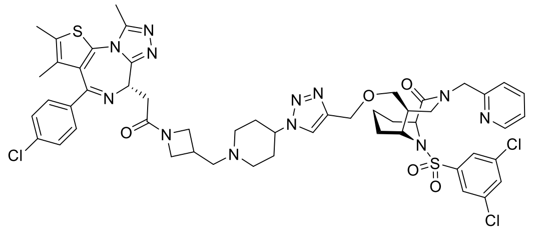

According to **GP-2**, the title compound **a11d** was synthesized from **a11** (6.1 mg, 10 µmol, 1.00 eq.) and (1S,5S,6R)-10-((3,5-dichlorophenyl)sulfonyl)-5-((prop-2-yn-1-yloxy)methyl)-3-(pyridin-2-ylmethyl)-3,10-diazabicyclo[4.3.1]decan-2-one [20] (5.2 mg, 10 µmol, 1.00 eq.). The obtained product was further purified by prep. HPLC (5-100 % B).

**Yield:** 7.4 mg (6.7 µmol, 67 %).

**Purity:** 98 % (HPLC, UV-absorption 220 nm)

**Physical State:** yellow solid.

**HRMS** (ESI, m/z) calcd. for $C_{52}H_{58}Cl_3N_{12}O_5S_2$ [M+H$^+$]:1099.31546, found: 1099.31775.
